# Proteomic and functional mapping of cardiac Na_V_1.5 channel phosphorylation reveals multisite regulation of surface expression and gating

**DOI:** 10.1101/2020.04.29.067835

**Authors:** Maxime Lorenzini, Sophie Burel, Adrien Lesage, Emily Wagner, Camille Charrière, Pierre-Marie Chevillard, Bérangère Evrard, Dan Maloney, Kiersten M. Ruff, Rohit V. Pappu, Stefan Wagner, Jeanne M. Nerbonne, Jonathan R. Silva, R. Reid Townsend, Lars S. Maier, Céline Marionneau

## Abstract

Phosphorylation of Na_V_1.5 channels regulates cardiac excitability, yet the phosphorylation sites regulating channel function and the underlying mechanisms remain largely unknown. Using a systematic quantitative phosphoproteomic approach, we analyzed Na_V_1.5 channel complexes purified from non-failing and failing mouse left ventricles, and we identified 42 phosphorylation sites on Na_V_1.5. Most sites are clustered, and three of these clusters are highly phosphorylated. Analyses of phosphosilent and phosphomimetic Na_V_1.5 mutants revealed the roles of three phosphosites in regulating Na_V_1.5 channel expression and gating. The phosphorylated serines-664 and -667 regulate the voltage-dependence of channel activation in a cumulative manner, whereas phosphorylation of the nearby serine-671, which is increased in failing hearts, decreases cell surface Na_V_1.5 expression and peak Na^+^ current. No additional roles could be assigned to the other clusters of phosphosites. Taken together, the results demonstrate that ventricular Na_V_1.5 is highly phosphorylated, and that the phosphorylation-dependent regulation of Na_V_1.5-encoded channels is highly complex, site-specific and dynamic.

**Abbreviations:** A, alanine; E, glutamate; HEK-293, Human Embryonic Kidney 293 cells; I_Na_, peak Na^+^ current; I_NaL_, late Na^+^ current; IP, immunoprecipitation; mαNa_V_PAN, anti-Na_V_ channel subunit mouse monoclonal antibody; MS, Mass Spectrometry; MS1, mass spectrum of peptide precursors; MS2 or MS/MS, fragmentation mass spectrum of peptides selected in narrow mass range (2 Da) from MS1 scan; Na_V_, voltage-gated Na^+^ channel; pS, phosphoserine; pT, phosphothreonine; S, serine; T, threonine; TAC, Transverse Aortic Constriction; TMT, Tandem Mass Tag.

## Introduction

Voltage-gated Na^+^ (Na_V_) channels are key determinants of myocardial excitability, and defects in Na_V_ channel expression or functioning in the context of inherited or acquired cardiac disease increase propensity to develop lethal arrhythmias (1). Ventricular Na_V_ channels, composed primarily of the Na_V_1.5 channel pore-forming subunit, in association with several accessory/regulatory proteins, generate the transient, peak Na^+^ current (I_Na_) responsible for the action potential upstroke and rapid intercellular conduction. While cardiac myocyte Na_V_ channels inactivate quickly, there is a finite probability (∼0.5%) of channels remaining open, resulting in the late component of the Na^+^ current (I_NaL_), which contributes to determining action potential duration. In the ventricular myocardium, the Na_V_1.5 protein is subject to many post-translational modifications, each of which fine-tunes channel expression and functioning in various physiological and disease contexts. Among the eleven different post-translational modifications previously shown to regulate cardiac Na_V_1.5 channels, phosphorylation at serine, threonine and tyrosine residues is certainly the best characterized (reviewed in (2), (3–5)).

A role for phosphorylation in regulating cardiac Na_V_1.5 channels was first suggested in a pioneering study demonstrating that β-adrenergic receptors couple to Na_V_ channels not only through a direct G-protein pathway, but also through an indirect, Protein Kinase A (PKA)-dependent pathway (6). The involvement of several additional kinases and phosphatases in regulating both I_Na_ and/or I_NaL_ later spotlighted the functional relevance of cardiac Na_V_1.5 channel phosphorylation. Perhaps most strikingly, progress in mass spectrometry (MS)-based phosphoproteomic analyses recently buttressed the field by revealing the existence of multiple phosphorylation sites on native ventricular (7, 8) and heterologously-expressed (9) Na_V_1.5 channels. Yet, little is known about the roles and detailed molecular mechanisms that underlie phosphorylation-dependent regulations of cardiac Na_V_1.5 channels.

Phosphorylation of Na_V_1.5 channels has also recently been suggested as an arrhythmogenic mechanism in heart failure (10–15). The Na_V_ channel defects associated with heart failure are most often characterized by increased I_NaL_ and/or decreased I_Na_, contributing to action potential prolongation and conduction slowing, respectively (10,14,16–19). The increase in I_NaL_ has reportedly been linked to the activation of kinases, mainly the Ca^2+^/Calmodulin-dependent protein Kinase II (CaMKII) (10,13–15), and several studies have focused on identifying the CaMKII-dependent Na_V_1.5 phosphorylation sites (7,9,20,21). Notably, increased CaMKII-dependent Na_V_1.5 phosphorylation at serine-571 has been reported and suggested to increase I_NaL_ in non-ischemic human heart failure (12) and in animal models of heart disease (11,12,14). Nevertheless, Na_V_1.5 channel phosphorylation may not be the sole mechanism involved in the observed pathophysiological defects, as other evidence suggests roles for upregulation of the neuronal Na_V_1.1 (18, 22), Na_V_1.6 (18) or Na_V_1.8 (23) channels. Intensive investigations were also undertaken to understand the causes of the reduced I_Na_, yet the detailed underlying molecular mechanisms remain unclear. While most studies failed to detect any changes in Na_V_1.5 transcript or total protein expression in failing human hearts (24) or in animal models of heart failure (17, 19), several mechanisms have been suggested to contribute to reduced I_Na_, including the generation of a C-terminal truncation splicing variant switch in Na_V_1.5 transcripts (25, 26), elevated NADH and reactive oxygen species production (27), or increased intracellular Ca^2+^ concentration and subsequent increased expression of the E3 ubiquitin ligase Nedd4-2 (28). In line with reduced I_Na_, a recent study using high-resolution imaging and functional techniques showed a reduction in Na_V_1.5 cluster size and a corresponding decreased number of open channels at the lateral membranes of ventricular myocytes from mice subjected to Transverse Aortic Constriction (TAC), without any changes in Na_V_1.5 transcript or total protein expression (29).

In this study, we investigated the patterns of phosphorylation of native mouse left ventricular Na_V_1.5 channels and the roles of identified phosphorylation sites in regulating Na_V_1.5 channel expression and functioning. Using quantitative MS-based phosphoproteomic analyses, we identified and quantified *in situ* the native phosphorylation sites of the Na_V_1.5 in a mouse model of pressure overload-induced heart failure produced by TAC. By analyzing the expression and the functional properties of phosphosilent and phosphomimetic Na_V_1.5 mutant channels in human embryonic kidney (HEK-293) cells, as well as simulating the consequences of phosphorylation on Na_V_1.5 peptide segment expansion, we identified phosphorylation hot spots for regulation of both channel cell surface expression and gating.

## Results

### Purification and characterization of Na_V_ channel complexes from Sham and TAC mouse left ventricles

Na_V_ channel complexes from four Sham-operated and five TAC mouse left ventricles were purified by immunoprecipitation (IP) using an anti-Na_V_PAN mouse monoclonal (mαNa_V_PAN) antibody, and characterized using the quantitative isobaric tandem mass tag (TMT)-based analysis. As illustrated in **Figure 1 - Table Supplement 1**, and consistent with previous findings (14), the echocardiographic analysis confirmed increased left ventricular masses (LVM/BW ratios), reduced ejection fractions, but unaltered left ventricular end-diastolic diameters (LVID;d) five weeks after the TAC surgery, demonstrating left ventricular concentric hypertrophy and systolic contractile dysfunction or heart failure in the TAC animals. Western blot analyses of total lysates showed similar total Na_V_1.5 protein expression in Sham and TAC left ventricles, which resulted in similar Na_V_1.5 immunoprecipitation yields in the nine samples (**Figure 1 - Figure Supplement 1A**). Isolated Na_V_ channel complexes were then digested with trypsin, and peptide mixtures were labeled with different TMT tags and combined in the same TMT set for multiplexed MS/MS analysis. As illustrated in **Table 1**, the Na_V_1.5 protein was the most represented protein in the mαNa_V_PAN-IPs, with 310 unique and Na_V_1.5-specific peptides identified and 56% amino acid sequence coverage (70% with the transmembrane domains removed, **Figure 2**).

**Figure 1.**
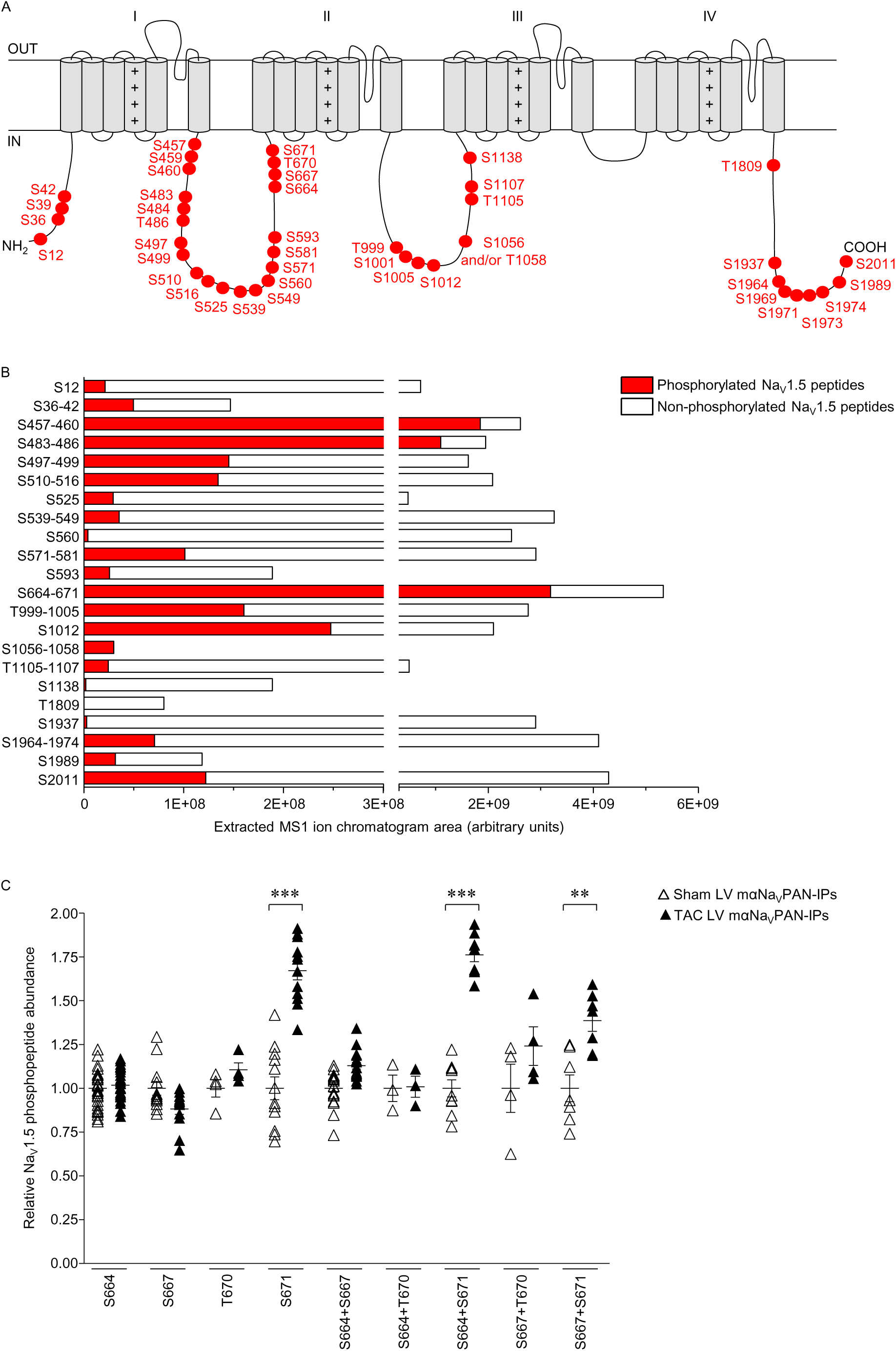
Localization and quantification of 42 MS-identified Na_V_1.5 phosphorylation sites in mαNa_V_PAN-IPs from Sham and TAC mouse left ventricles. (**A**) Schematic representation of phosphorylation sites on the Na_V_1.5 protein (UniProt reference sequence K3W4N7). Two phosphorylation site locations are possible at amino acids S1056-T1058. (**B**) The areas of extracted MS1 ion chromatograms, corresponding to MS2 spectra assigning phosphorylated (in red) and non-phosphorylated (in white) Na_V_1.5 peptides at indicated phosphorylation site(s), in mαNa_V_PAN-IPs from Sham and TAC left ventricles are indicated. (**C**) Distributions and mean ± SEM relative abundances of individual Na_V_1.5 phosphopeptides allowing assignments of indicated phosphorylation site(s), in TAC LV (n=5, in black), *versus* Sham LV (n=4, in white), mαNa_V_PAN-IPs, were obtained using TMT reporter ion intensities. The relative abundances of Na_V_1.5 phosphopeptides exhibiting phosphorylation(s) on serine-671 (S671, n=12 peptides) alone, or in combination with serines-664 (S664 + S671, n=9 peptides) or -667 (S667 + S671, n=7 peptides) are increased (\*\**p*<0.01, \*\*\**p*<0.001, Mann-Whitney test) in TAC LV, *versus* Sham LV, mαNa_V_PAN-IPs.

**Figure 2.**
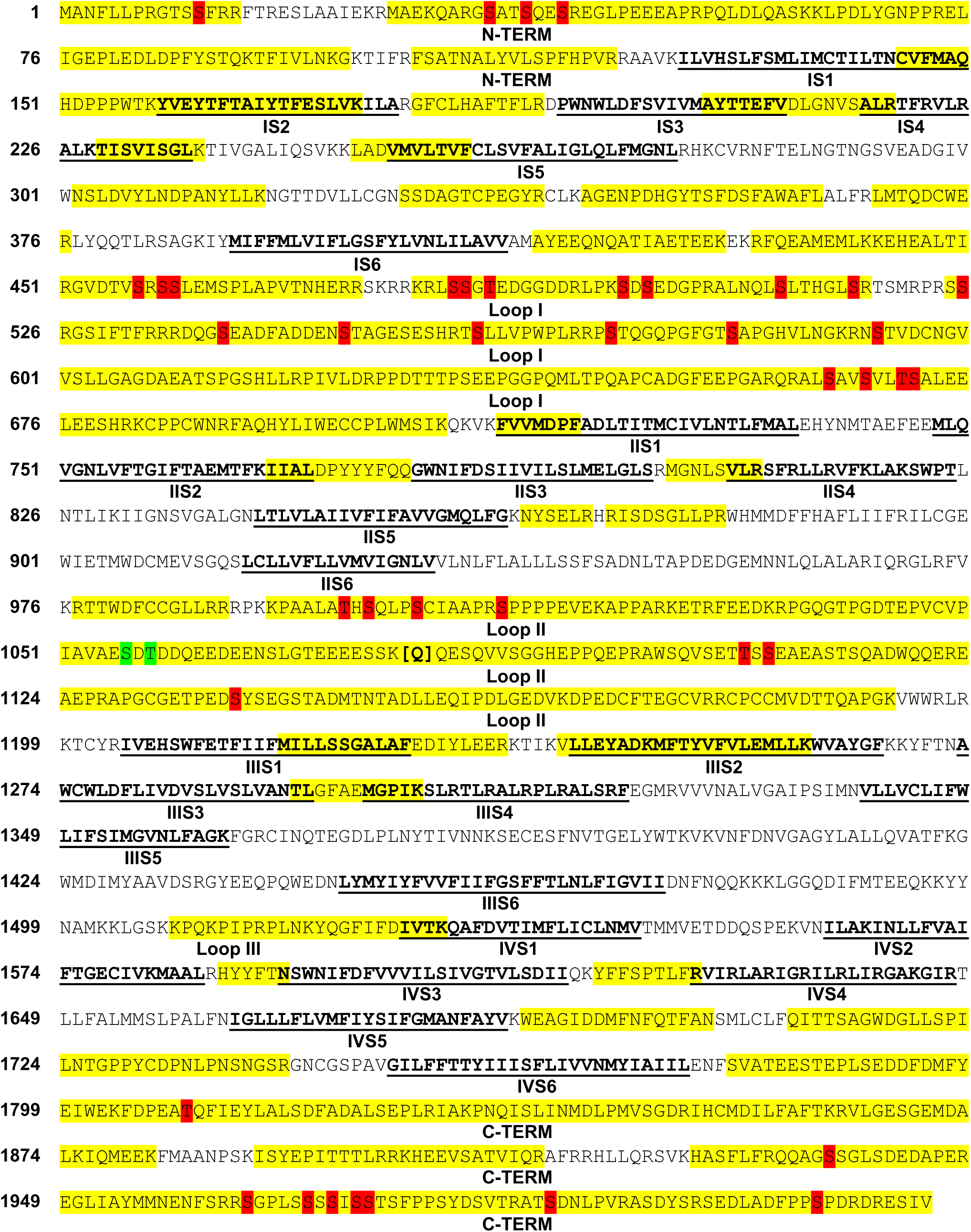
Na_V_1.5 amino acid sequence coverage and localization of 42 Na_V_1.5 phosphorylation sites in mαNa_V_PAN-IPs from Sham and TAC mouse left ventricles. Covered sequence and MS-identified phosphorylation sites are highlighted in yellow and red, respectively; transmembrane segments (S1-S6) in each domain (I-IV) are in bold and underlined in black; and loops I, II and III correspond to interdomains I-II, II-III and III-IV, respectively. Two peptides differing by the presence or absence of a glutamine (Q) at position-1080 were detected, reflecting the expression of two Na_V_1.5 variants (the Q1080 variant corresponds to UniProt reference sequence K3W4N7; and the Q1080del variant corresponds to Q9JJV9). Two phosphorylation site locations are possible at amino acids S1056-T1058 (in green).

**Table 1.** Proteins identified in immunoprecipitated NaV channel complexes from Sham and TAC mouse left ventricles using MS

| Protein | UniProt accession number | Number of exclusive unique peptides | % Amino acid sequence coverage | TAC/Sham ratio | Protein | UniProt accession number | Number of exclusive unique peptides | % Amino acid sequence coverage | TAC/Sham ratio |
| --- | --- | --- | --- | --- | --- | --- | --- | --- | --- |
| Na<sub>v</sub>1.5 (Q1080)<br>Na<sub>v</sub>1.5 (Q1080del) | K3W4N7<br>Q9JJV9 | 310 | 56% | 1.0 | N-cadherin | P15116 | 13 | 24% | 0.8 |
| Na<sub>v</sub>1.4 | G3X8T7 | 86 | 40% | 0.8 | Plakophilin-2 | Q9CQ73 | 13 | 20% | 0.7 |
| Na<sub>v</sub>1.3 | A2ASI5 | 24 | 17% | 0.9 | Plakophilin-1 | P97350 | 5 | 8% | 0.8 |
| Na<sub>v</sub>1.8 | Q6QIY3 | 13 | 9% | 0.9 | Telethonin | O70548 | 5 | 44% | 0.8 |
| Na<sub>v</sub>1.7 | Q62205 | 3 | 2% | 0.7 | Desmoglein-2 | O55111 | 13 | 19% | 0.7 |
| Na<sub>v</sub>1.9 | Q9R053 | 1 | 1% | 1.2 | Flotillin-1 | O08917 | 22 | 59% | 0.9 |
| Na<sub>v</sub>1.1 | A2APX8 | 1 | 0.5% | 1.3 | Flotillin-2 | Q60634 | 21 | 54% | 0.8 |
| Na<sub>v</sub>1.2 | B1AWN6 | 1 | 0.5% | 1.2 | 14-3-3 zeta/delta | P63101 | 5 | 27% | 0.9 |
| Na<sub>v</sub>1.6 | F6U329 | 1 | 0.6% | 1.0 | 14-3-3 epsilon | P62259 | 3 | 17% | 1.0 |
| Navβ4 | Q7M729 | 11 | 44% | 0.7 | 14-3-3 gamma | P61982 | 3 | 14% | 1.0 |
| Navβ1 | P97952 | 9 | 41% | 1.0 | 14-3-3 beta/alpha | Q9CQV8 | 1 | 6% | 1.1 |
| Navβ2 | Q56A07 | 6 | 37% | 0.8 | 14-3-3 sigma | O70456 | 1 | 3% | 0.9 |
| Calmodulin | Q3UKW2 | 12 | 53% | 0.8 | Slmap | Q3URD3 | 1 | 1% | 1.1 |
| FHF2-VY | P70377 | 37 | 84% | 0.8 | αB-crystallin | P23927 | 14 | 71% | 0.7 |
| FHF4 | P70379 | 1 | 7% | 0.7 | Cx43 | P23242 | 15 | 50% | 0.7 |
| Ankyrin-G | G5E8K5 | 47 | 32% | 1.0 | Kir2.1 | P35561 | 3 | 11% | 0.8 |
| Ankyrin-R | Q02357 | 2 | 2% | 1.0 | CaMKII<sub>γ</sub> | Q923T9 | 11 | 26% | 0.8 |
| Ankyrin-B | S4R245 | 1 | 0.4% | 0.8 | CaMKIIδ | E9Q1T1 | 2 | 6% | 1.1 |
| Dystrophin | P11531 | 88 | 29% | 0.9 | CaMKIIα | P11798 | 2 | 4% | 0.8 |
| α-1-syntrophin | A2AKD7 | 20 | 53% | 0.8 | CaMKIIβ | Q5SVI2 | 1 | 3% | 0.9 |
| β-2-syntrophin | Q61235 | 19 | 35% | 0.9 | PKA, catalytic subunit, α | P05132 | 2 | 9% | 1.0 |
| α-actinin-2 | Q9JI91 | 50 | 60% | 0.5 | PKAβ-2 | Q6PAM0 | 1 | 4% | 1.1 |
| Caveolin-1 | P49817 | 6 | 39% | 0.8 | PKCβ | P68404 | 2 | 3% | 1.6 |
| Caveolin-2 | Q9WVC3 | 3 | 28% | 0.9 | Fyn | P39688 | 1 | 2% | 1.1 |
| Caveolin-3 | P51637 | 2 | 11% | 0.8 | PP2A-C | P63330 | 1 | 3% | 1.1 |
| Vinculin | Q64727 | 25 | 29% | 0.7 |  |  |  |  |  |

The numbers of exclusive unique peptides, the percent (%) amino acid sequence coverages and the fold change abundance ratios in TAC LV (n=5) versus Sham LV (n=4) mNa<sub>v</sub>PAN-IPs for each identified protein are presented. No significant differences in protein abundance were observed between TAC and Sham IPs. Abbreviations: Na<sub>v</sub>, voltage-gated Na<sup>+</sup> channel; FHF, Fibroblast growth factor Homologous Factor; Slmap, sarcolemmal membrane-associated protein; Cx, connexin; Kir, inward rectifier K<sup>+</sup> channel; CaMKII, Ca<sup>2+</sup>/Calmodulin-dependent protein Kinase II; PKA, Protein Kinase A; PKC, Protein Kinase C. PP2A-C, Catalytic subunit of Protein Phosphatase 2A.

Consistent with the homogenous yields in the Na_V_1.5 immunoprecipitation, the relative abundance of the Na_V_1.5 peptides detected by MS in the nine samples was similar, and used for normalization of each single protein and peptide abundance (**Figure 1 - Figure Supplement 1B**). Accordingly, the distribution of normalized abundance ratios of Na_V_1.5 peptides (in log_2_) in TAC, *versus* Sham, mαNa_V_PAN-IPs was centered on zero (**Figure 1 - Figure Supplement 1C**). Altogether, therefore, these observations attest to a high reproducibility across biological replicates, and a low technical variability inherent to experimental procedures. Of note, and as described previously (30), two Na_V_1.5 peptides differing by the presence or absence of a glutamine (Q) at position-1080 were detected (**Table 1 & Figure 2**), reflecting the expression of at least two distinct Na_V_1.5 splice variants in mouse left ventricles; Q1080del corresponding to the commonly reported hH1C variant. Interestingly, these analyses also allowed the identification of eight additional Na_V_ channel pore-forming subunits, among which Na_V_1.4 is the most abundant, with 86 unique Na_V_1.4-specific peptides detected (**Table 1**). In addition, several previously identified Na_V_1.5 channel associated/regulatory proteins, including calmodulin, the VY variant of Fibroblast growth factor Homologous Factor 2 (FHF2-VY) and ankyrin-G, were detected, with no significant differences in abundance between Sham and TAC mαNa_V_PAN-IPs (**Table 1**).

### Identification and quantification of 42 Na_V_1.5 phosphorylation sites in Sham and TAC mouse left ventricles

The phosphoproteomic analysis of the mαNa_V_PAN-IPs from Sham and TAC mouse left ventricles allowed the unambiguous identification of 42 native phosphorylation sites in the Na_V_1.5 protein, 22 of which have never, to our knowledge, been previously described in native cardiac tissues (**Figures 1A & 2**). **Table 2** lists the phosphopeptides enabling the best phosphorylation site assignment(s) for each phosphorylation site; and corresponding MS/MS spectra are presented in **Table 2 - Figure Supplement 1**. Interestingly, the vast majority of these phosphorylation sites are clustered, with the first intracellular linker loop of Na_V_1.5 revealed as a hot spot for phosphorylation, with a total of 21 sites identified. Further label-free quantitative analysis of the areas of extracted MS1 peptide ion chromatograms revealed large differences in the relative abundances of the individual phosphopeptides, and the existence of three highly phosphorylated clusters at positions S457 to S460, S483 to T486, and S664 to S671 (**Figure 1B**). In addition, and in contrast to the other phosphorylation sites, the phosphorylated peptides assigning these three phosphorylation clusters are more abundant than their non-phosphorylated counterparts, suggesting that these sites are mostly phosphorylated in native Na_V_1.5 channels in wild-type mouse left ventricles. Looking into the detailed quantification of single phosphorylation sites inside each of these clusters, however, major differences in phosphopeptide abundance are evident (**Figure 1 - Figure Supplement 2**). This is the case, for example, of phosphorylation at S664 or S667, which is about 10-fold more abundant than at residues T670 or S671.

**Table 2.**
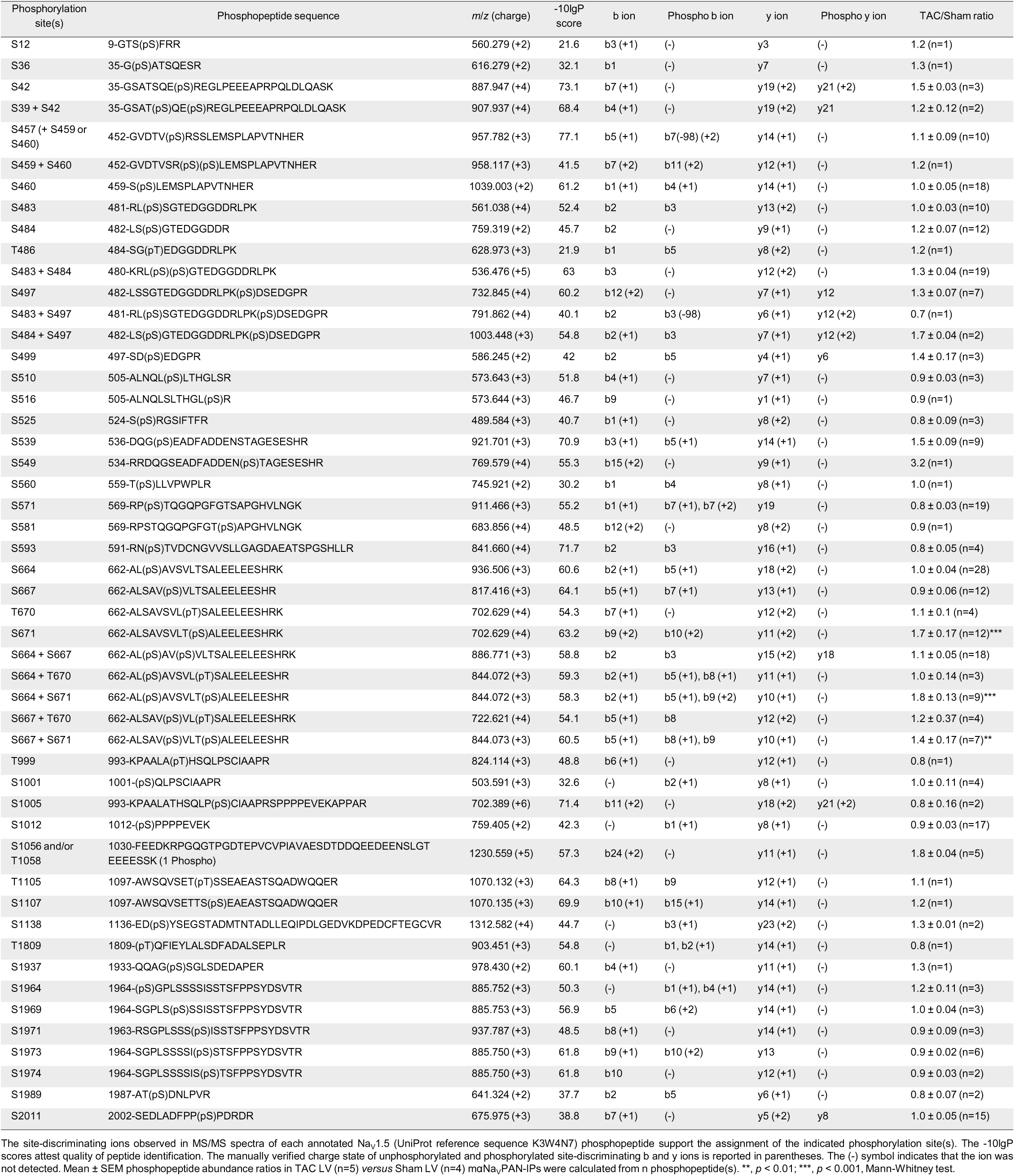
Phosphorylation sites, phosphopeptides and site-discriminating ions identified in immunoprecipitated NaV1.5 proteins from Sham and TAC mouse left ventricles using MS

To determine whether phosphorylation of Na_V_1.5 is regulated in heart failure, the relative abundance of each Na_V_1.5 phosphopeptide in TAC, *versus* Sham, mαNa_V_PAN-IPs was calculated using the relative abundance of TMT reporter ions. As illustrated in **Figure 1C**, peptides exhibiting phosphorylation(s) on serine-671 (S671) alone or in combination with serines-664 (S664 + S671) or -667 (S667 + S671) are significantly more abundant in the TAC, compared with the Sham, mαNa_V_PAN-IPs. The relative abundances of their non-phosphorylated counterparts, however, are similar in Sham and TAC mαNa_V_PAN-IPs (data not shown). Additionally, none of the other Na_V_1.5 phosphopeptides showed any significant differences in the Sham and TAC mαNa_V_PAN-IPs (**Table 2**). In addition to Na_V_1.5, four phosphorylation sites on Na_V_1.4 and one on Na_V_1.3 could also be detected (**Table 2 - Table Supplement 1 & Figure Supplement 1**). Taken together, these quantitative phosphoproteomic analyses identified 42 native phosphorylation sites on Na_V_1.5, among which three clusters of phosphorylation in the first loop of the channel are highly phosphorylated, and one serine at position-671 shows increased phosphorylation in TAC left ventricles.

### Functional mapping of Na_V_1.5 channel phosphorylation clusters

The identification of several clusters of phosphorylation sites on Na_V_1.5 suggests that these sites may be involved in the coordinated regulation of channel expression and/or function. Out of the eight clusters of phosphorylation identified in the mouse Na_V_1.5 protein, seven are conserved in the human Na_V_1.5 protein sequence; only the mouse T1105 is not conserved (**Figure 3 - Figure Supplement 1**). In order to investigate the functional roles of these (seven) phosphorylation clusters, phosphosilent and phosphomimetic Na_V_1.5 channel constructs in the human Na_V_1.5 hH1C cDNA sequence were generated, transiently expressed in HEK-293 cells, and characterized in whole-cell voltage-clamp recordings. In the phosphosilent constructs, mutations were introduced to replace serines/threonines with alanines, whereas in the phosphomimetic constructs, mutations were introduced to substitute glutamates for serines/threonines, to mimic phosphorylation.

As illustrated in **Figure 3B**, these whole-cell voltage-clamp analyses demonstrated that the voltage-dependence of activation of Na_V_1.5-S664-671A phosphosilent channels is significantly (*p*<0.001) shifted towards depolarized potentials, compared to WT channels (see distributions, detailed properties and statistics in **Figure 3 - Figure Supplement 2A & Table Supplement 1**). The activation curve of the Na_V_1.5-S664-671E phosphomimetic channel was also significantly (*p*<0.001) shifted, although to a lesser extent, when compared with the phosphosilent channel. Together, therefore, these findings suggest that the S664-671 cluster is phosphorylated in HEK-293 cells, and that disruption of phosphorylation at these sites shifts the voltage-dependence of channel activation towards depolarized potentials. In addition, the time to peak Na^+^ current (**Figure 3D**), as well as the inactivation time constants, τ_fast_ and τ_slow_ (**Figures 3E & 3F**), were shifted towards depolarized potentials until reaching full activation at ∼0 mV. The peak Na^+^ current density of the Na_V_1.5-S664-671E phosphomimetic channel was significantly (*p*<0.05) reduced compared to the WT channel, whereas no significant changes were observed with the Na_V_1.5-S664-671A phosphosilent channel (**Figure 3C,** see distributions at -20 mV and statistics in **Figure 3 - Figure Supplement 2B & Table Supplement 1**). In contrast, the voltage-dependence of steady-state inactivation (**Figure 3B**) and the kinetic of recovery from inactivation (**Figure 3 - Figure Supplement 2D & Table Supplement 1**) of both S664-671 phosphomutant channels were not changed. Additionally, and to our surprise, no differences in current densities, or in the kinetics or voltage-dependences of current activation and inactivation, or in the kinetics of recovery from inactivation were observed for any of the six other heterologously-expressed (in HEK-293 cells) paired phosphosilent or phosphomimetic Na_V_1.5 channels (**Figure 3 - Figure Supplement 2 & Table Supplement 1**). Taken together, therefore, these analyses revealed a key role for phosphorylation at S664-671 in regulating the voltage-dependence of Na_V_1.5 channel activation and peak Na^+^ current density, whereas regulation mediated by the other phosphorylation sites investigated most likely involve more complex mechanisms.

**Figure 3.**
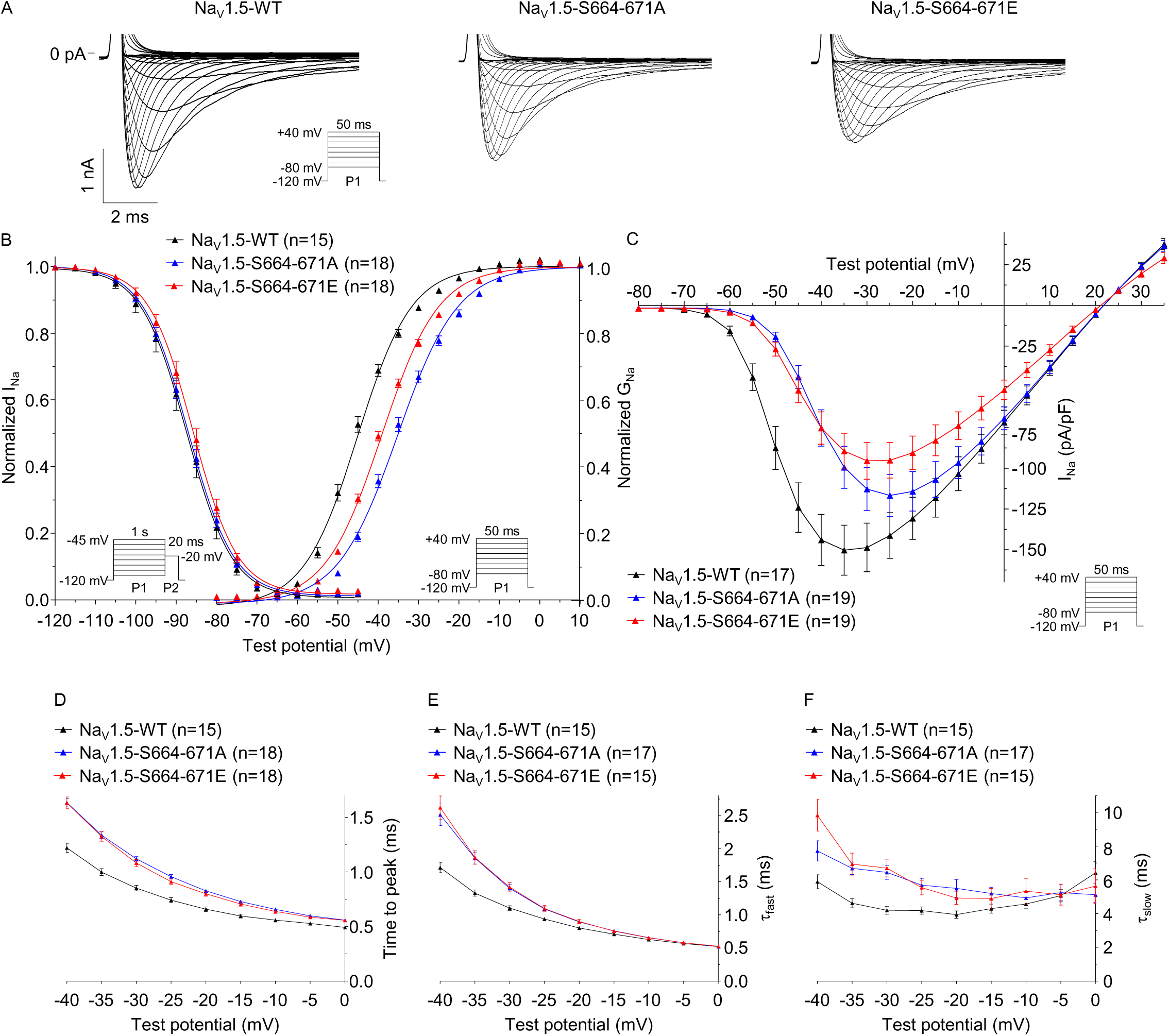
Phosphorylation at positions S664-671 regulates the voltage-dependence of current activation and peak Na^+^ current density. (**A**) Representative whole-cell voltage-gated Na^+^ currents recorded forty-eight hours following transfection of HEK-293 cells with Na_V_1.5-WT + Na_V_β1 (black), Na_V_1.5-S664-671A + Na_V_β1 (blue) or Na_V_1.5-S664-671E + Na_V_β1 (red) using the protocols illustrated in each panel. Scale bars are 1 nA and 2 ms. (**B**) Voltage-dependences of current activation and steady-state inactivation. The voltage-dependence of current activation is shifted towards depolarized potentials in cells expressing the Na_V_1.5-S664-671A or Na_V_1.5-S664-671E mutants, compared to cells expressing Na_V_1.5-WT. (**C**) Mean ± SEM peak Na^+^ current (I_Na_) densities plotted as a function of test potential. The peak I_Na_ density is reduced in cells expressing the Na_V_1.5-S664-671E mutant, compared to cells expressing Na_V_1.5-WT. Mean ± SEM times to peak (**D**), and fast (τ_fast_, **E**) and slow (τ_slow_, **F**) inactivation time constants plotted as a function of test potential. The time to peak, τ_fast_ and τ_slow_ are higher in cells expressing the Na_V_1.5-S664-671A or Na_V_1.5-S664-671E mutants, compared to cells expressing Na_V_1.5-WT. Current densities, time- and voltage-dependent properties, as well as statistical comparisons across groups, are provided in **Figure 3 - Figure Supplement 2 & Table Supplement 1**.

### Phosphorylation at S664 and S667 shifts the voltage-dependence of current activation towards hyperpolarized potentials whereas phosphorylation at S671 decreases the peak Na^+^ current density

To decipher the respective contributions of the S664, S667, T670 and S671 phosphorylation sites in regulating the voltage-dependence of current activation and peak Na^+^ current density, each of these serines/threonine was mutated individually to alanine or glutamate, and the densities and properties of Na^+^ currents from single phosphosilent or phosphomimetic channels were examined in transiently transfected HEK-293 cells. These analyses showed that the voltage-dependences of activation of the Na_V_1.5-S664 (**Figure 4A**) and Na_V_1.5-S667 (**Figure 4B**) phosphomutant channels are significantly (*p*<0.001) shifted towards depolarized potentials, compared to the WT channels, whereas no changes were observed with the Na_V_1.5-T670 or Na_V_1.5-S671 phosphomutant channels (**Figures 4C & 4D**, see detailed properties and statistics in **Figure 4 - Table Supplement 1**). Of note, the ∼6 mV shifts observed with the single Na_V_1.5-S664 and Na_V_1.5-S667 phosphomutant channels were two-fold smaller than the ∼10 mV shift obtained with the quadruple Na_V_1.5-S664-671A phosphosilent channel, suggesting that the effects at S664 and S667 are additive. Additionally, these analyses revealed that sole the Na_V_1.5-S671E phosphomimetic channel shows a significant (*p*<0.05) decrease in peak Na^+^ current density (**Figure 4H**), whereas none of the other single phosphomutant channels showed any significant differences (**Figures 4E, 4F & 4G**).

**Figure 4.**
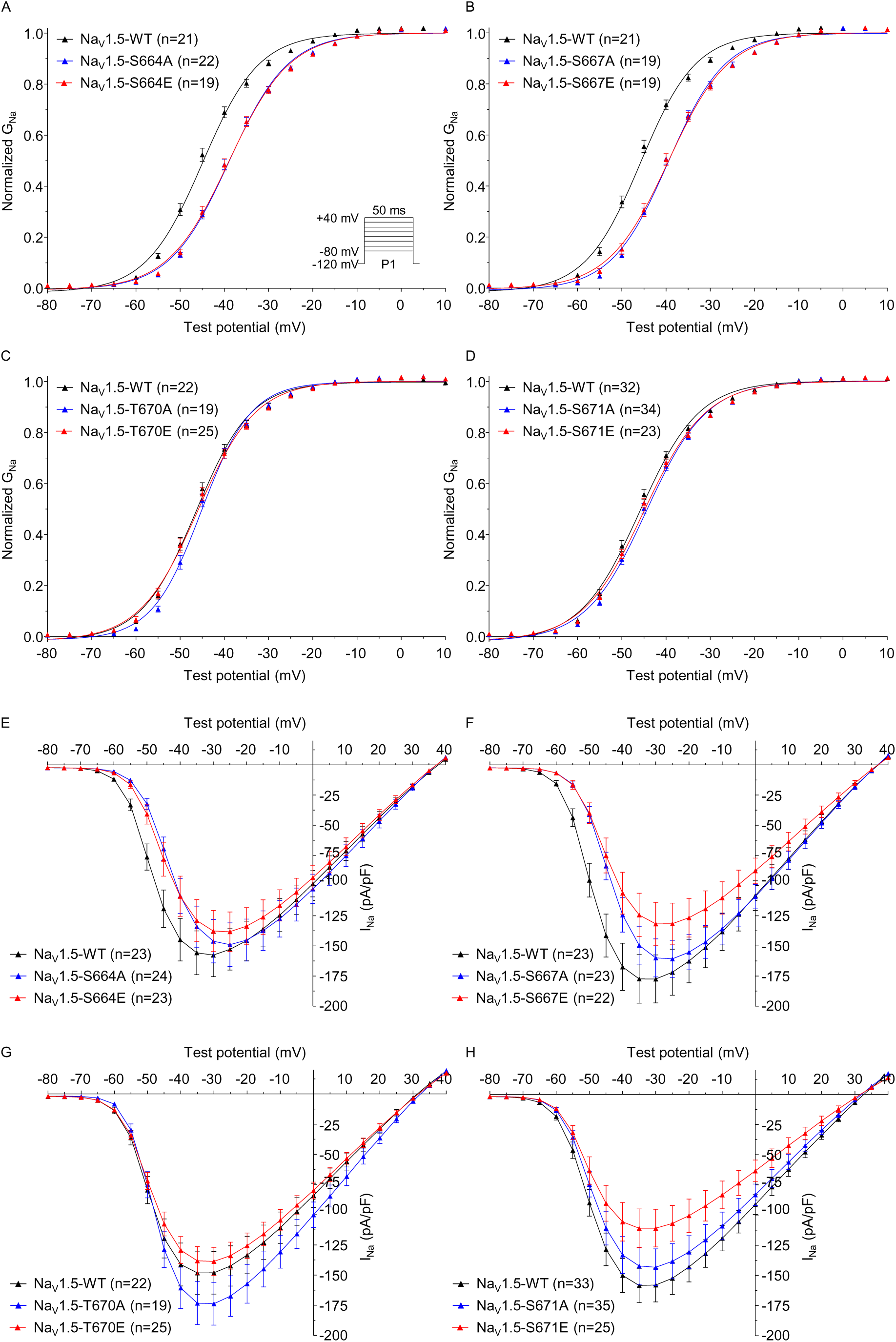
Phosphorylation at S664 and S667 regulates the voltage-dependence of current activation, whereas phosphorylation at S671 regulates the peak Na^+^ current density. Currents were recorded as described in the legend to Figure 3. The voltage-dependence of current activation is shifted towards more depolarized potentials in cells expressing Na_V_1.5-S664A (**A**), Na_V_1.5-S664E (**A**), Na_V_1.5-S667A (**B**) or Na_V_1.5-S667E (**B**), compared to cells expressing Na_V_1.5-WT, whereas no significant differences are observed with the Na_V_1.5-T670 (**C**) or Na_V_1.5-S671 (**D**) phosphomutants. (**E to H**) The mean ± SEM peak Na^+^ current (I_Na_) densities are plotted as a function of test potential. The peak I_Na_ density is reduced in cells expressing Na_V_1.5-S671E (**H**), compared to cells expressing Na_V_1.5-WT, whereas no significant differences are observed with the other phosphomutants. Current densities, time- and voltage-dependent properties, as well as statistical comparisons across groups, are provided in **Figure 4 - Table Supplement 1**.

Because phosphorylation at S671 was found to be increased in the TAC, compared with the Sham, mαNa_V_PAN-IPs (**Figure 1C**), and because it was previously suggested that phosphorylation of Na_V_1.5 may mediate increased I_NaL_ in heart failure (10–15), additional voltage-clamp experiments were designed to test whether phosphorylation at S671 regulates I_NaL_. These analyses showed that none of the single mutations at S671, or quadruple mutations at S664-671 affect TTX-sensitive I_NaL_ density in HEK-293 cells (**Figure 4 - Figure Supplement 1**). Altogether, therefore, these analyses suggest that phosphorylation at S664 and S667 shifts the voltage-dependence of current activation towards hyperpolarized potentials in a cumulative manner, whereas phosphorylation at S671 decreases the peak Na^+^ current density.

### Phosphorylation at S671 decreases the cell surface expression of Na_V_1.5 channels

Additional cell surface biotinylation experiments in transiently transfected HEK-293 cells were designed to determine whether phosphorylation at S671 regulates the cell surface expression of the Na_V_1.5 channel protein. Interestingly, these experiments revealed that the cell surface expression of the Na_V_1.5-S671E phosphomimetic channel is significantly (*p*<0.001) decreased, compared with the WT or the Na_V_1.5-S671A phosphosilent channels, whereas no differences in total Na_V_1.5 protein expression were observed (**Figures 5A & 5B**). Importantly, the decrease observed with the phosphomimetic mutant, compared with the phosphosilent mutant, suggests that not only this channel locus, but most probably phosphorylation at this particular site, underlies the observed decrease in cell surface expression. Together with the electrophysiological findings, therefore, these biochemical analyses demonstrate a key role for S671 in regulating the cell surface expression of Na_V_1.5, and suggest that phosphorylation at this site decreases the cell surface expression of Na_V_1.5-encoded channels.

**Figure 5.**
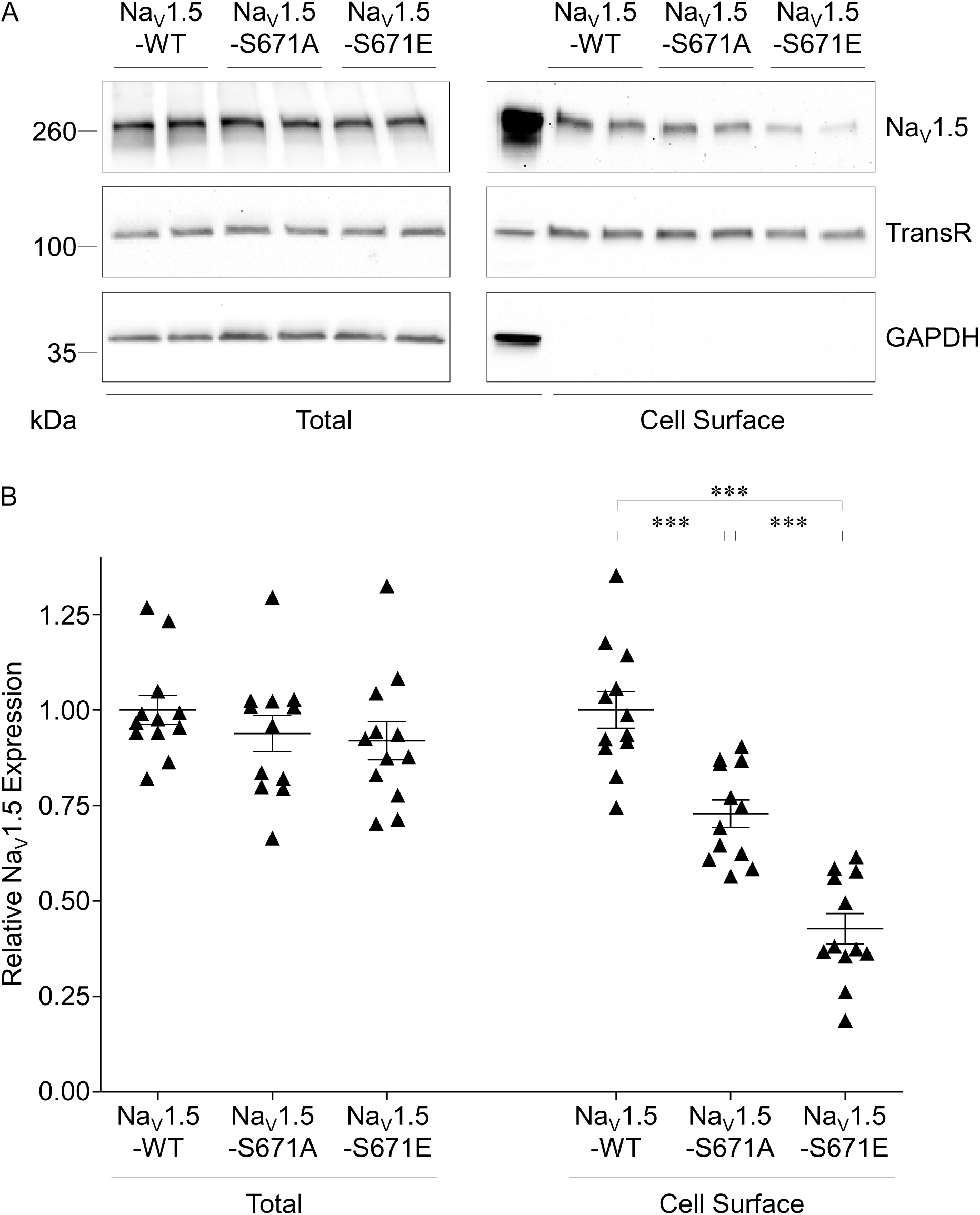
Phosphorylation at S671 regulates the cell surface expression of Na_V_1.5. (**A**) Representative western blots of total (left panel) and cell surface (right panel) Na_V_1.5 from HEK-293 cells transiently transfected with Na_V_1.5-WT + Na_V_β1, Na_V_1.5-S671A + Na_V_β1 or Na_V_1.5-S671E + Na_V_β1. Samples were probed in parallel with the anti-transferrin receptor (TransR) and anti-glyceraldehyde 3-phosphate dehydrogenase (GAPDH) antibodies. (**B**) Mean ± SEM total and cell surface Na_V_1.5 protein expression in transiently transfected HEK-293 cells (n=12 in 6 different experiments). Expression of Na_V_1.5 in each sample was first normalized to the TransR protein in the same blot and then expressed relative to Na_V_1.5 protein expression (total or cell surface) in cells transfected with Na_V_1.5-WT + Navβ1. Relative (mean ± SEM) Na_V_1.5 cell surface expression is different (\*\*\**p*<0.001, one-way ANOVA followed by the Dunnett’s post-hoc test) in cells expressing Na_V_1.5-WT, Na_V_1.5-S671A and Na_V_1.5-S671E channels.

### Simulated consequences of phosphorylation on the first intracellular linker loop of Na_V_1.5

Like many heavily phosphorylated protein segments (31, 32), the first two intracellular linker loops of Na_V_1.5 are predicted to be intrinsically disordered. Conformational heterogeneity is one of the defining hallmarks of intrinsically disordered regions (IDRs). Heterogeneity is manifest in the amplitude of fluctuations of overall size, shape, and local secondary structural preferences. There is growing recognition of sequence-specificity whereby the ensembles accessible to an IDR are governed by the amino acid composition, extent of phosphorylation, and patterning of residues within the linear sequence. These sequence-ensemble relationships can be uncovered using all atom simulations. Given the disparate timescales and length scales involved, a robust and efficient approach is to use Markov Chain Metropolis Monte Carlo (MC) simulations based on the ABSINTH implicit solvent model as implemented in the CAMPARI simulation (33–35). Here, we used simulations to quantify sequence-ensemble relationships for the first intracellular linker loop of human Na_V_1.5 containing the phosphorylation clusters S457-460, S483-486, S497-499, and S664-671 identified by mass spectrometry. For our simulations, we used segments between thirty and forty residues in length, containing each cluster in an approximately central position (441-480 for S457-460, 465-501 for S483-486, 481-515 for S497-499, 651-684 for S664-671). For each cluster, we performed simulations for the WT sequence, as well as phosphomimetic mutations where serine(s)/threonine(s) are replaced with glutamate(s). The results of simulations were analyzed using the device of internal scaling plots. These plots quantify the variation of ensemble-averaged distances between residues i and j as a function of sequence separation |j-i|. Multiple pairs of residues contribute to a given sequence separation |j-i|. The internal scaling profiles can be calibrated against reference profiled that pertain to two kinds of random-coil ensembles. These are designated as EV for excluded volume, which pertains to profiles extracted for self-avoiding walks, and FRC for Flory random coil, which pertains to profiles extracted Flory random coils. Details of these reference ensembles have been published elsewhere.

Simulations of the 441-480 segment showed that the conformational preferences of the unphosphorylated (WT) peptide are akin to those of the FRC reference (**Figure 6A**). This implies that sequence encodes a conformational averaging whereby the peptide-solvent and peptide-peptide interactions are mutually compensatory, thereby giving rise to an ensemble that is maximally heterogeneous. Introduction of phosphomimetic substitutions S457E, S459E and/or S460E did not have a large effect on the ensemble-averaged internal scaling profiles when compared to the unmodified sequence. We obtained similar results for the 465-501 segment, which is also largely unaffected by the introduction of the phosphomimetic mutation(s) of the S483-486 cluster (**Figure 6B**). Conversely, the 481-515 and 651-684 segments were noticeably sensitive to the addition of the negative charges (**Figures 6C & 6D**). When unphosphorylated, these segments preferred conformations that are considerably more compact than the FRC reference. Upon the introduction of cumulative phosphomimetic mutations, these segments gradually expanded in the direction of the EV limit.

**Figure 6.**
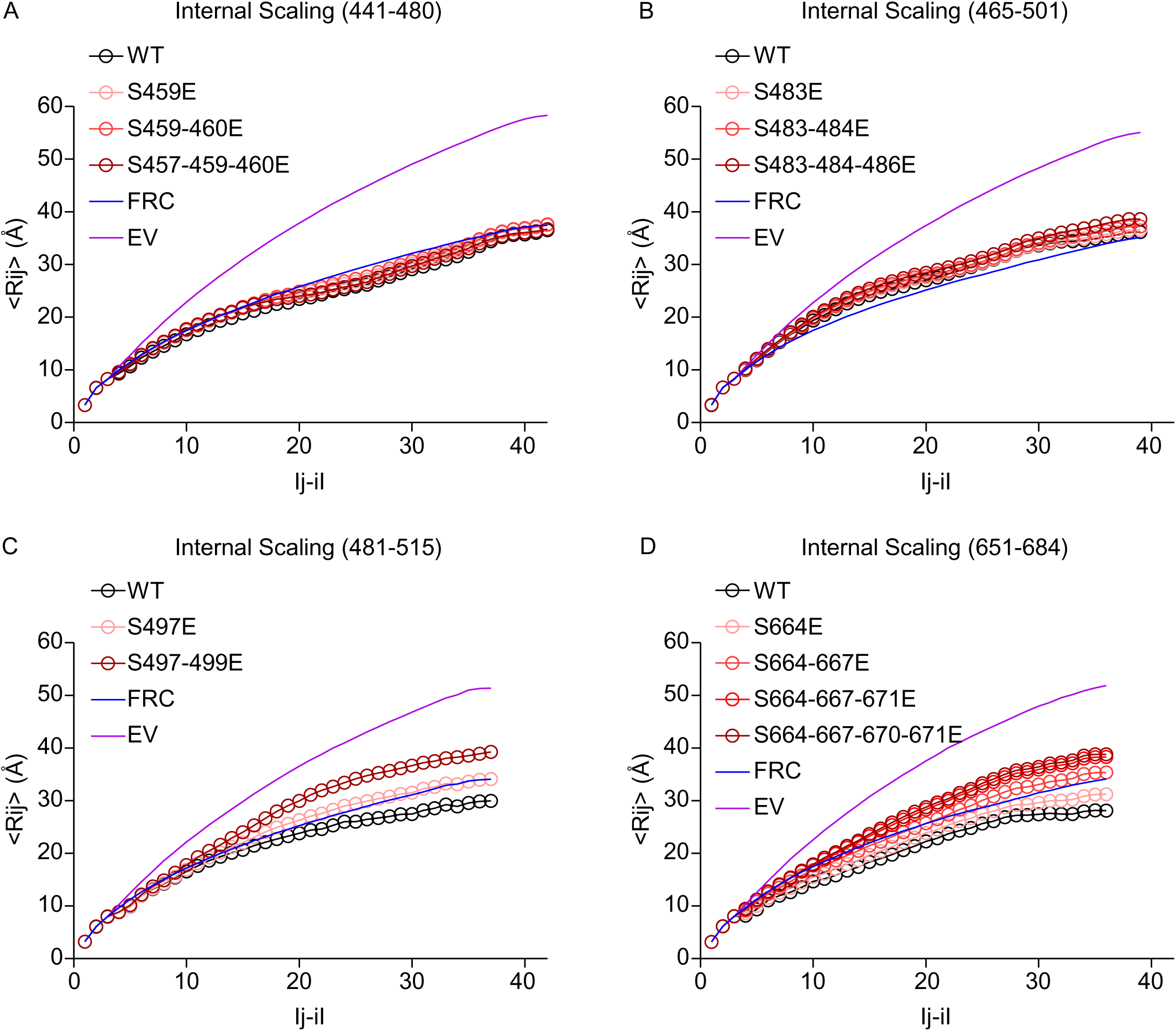
Simulations of phosphorylation of segments of the first intracellular linker loop of Na_V_1.5. The sequential distance between pair of residues is on the x-axis, and the average spatial distance between a pair of residues separated by the specified sequential distance is on the y-axis. <RIJ> is the average simulated spatial distance between all residue pairs separated in the amino acid sequence by |j-i| residues. The WT sequences are plotted in black. Phosphorylation is simulated by single or multiple replacement of serines/threonines with glutamates (E), and resulting simulations are plotted in gradation of reds. The Flory Random Coil (FRC, in blue) and Excluded Volume (EV, in purple) limits are plotted for reference (see text). (**A**) Sequence 441-480 contains the phosphosites S457, S459 and S460; (**B**) sequence 465-501 contains the phosphosites S483, S484 and T486; (**C**) sequence 481-515 contains the phosphosites S497 and S499; and (**D**) sequence 651-684 contains the phosphosites S664, S667, T670 and S671.

Taken together, the results suggest that the intrinsic conformational preferences of the WT sequence dictate the extent of responsiveness of the conformational ensemble to multisite phosphorylation. Sequence stretches that have an intrinsic preference for FRC-like conformations are relatively insensitive to phosphomimetic substitutions of serine/threonine residues. This insensitivity has been quantified for IDRs that undergo multisite phosphorylation (36). In contrast, sequences that have an intrinsic preference for compact conformations become responsive to phosphomimetic substitutions. This would appear to derive from the increased fraction of charged residues (which engenders preferential solvation) and electrostatic repulsions (34). The fraction of charged residues (FCR) and the net charge per residue (NCPR) are known to be direct determinants of the conformational preferences of IDRs (37). Both the 441-480 and 465-501 segments have a higher FCR than the 481-515 and 651-684 segments. The addition of a single negative charge would lead to a greater percent increase in the FCR of the 481-515 and 651-684 segments than it would for the 441-480 and 464-501 segments. As the latter are already expanded, additional charges do not have a large impact on the conformational preference. The more compact starting point of the former allows the phosphomimetic mutations to have a greater effect. While the results for three of the clusters were consistent with the experimental data, those for the S497-499 cluster present an apparent inconsistency. These observations suggest that the ability of this segment to expand due to the addition of charge is not connected to channel gating. Together with the electrophysiological analyses, therefore, these simulations suggest that the effect of phosphorylation at S664 and S667 on the voltage-dependence of channel activation is mediated by the expansion of the area containing the phosphorylation sites, and that this expansion is likely to regulate channel activation allosterically.

## Discussion

The results presented here provide a novel, detailed phosphorylation map of the native mouse left ventricular Na_V_1.5 channel protein, and identify the functional roles of three of these phosphorylation sites in regulating the expression and gating properties of Na_V_1.5-encoded channels. The highly phosphorylated S664 and S667 shift the voltage-dependence of channel activation towards hyperpolarized potentials in an additive manner, whereas phosphorylation at S671, which is increased in TAC mouse left ventricles, decreases Na_V_1.5 cell surface expression and peak Na^+^ current density. No additional roles could be assigned to the other clusters of Na_V_1.5 phosphorylation sites, suggesting additional complexity in the mechanisms mediating the phosphorylation-dependent regulation of cardiac Na_V_1.5 channels.

### Phosphorylation map of native mouse left ventricular Na_V_1.5 channels

The present phosphoproteomic analysis confidently identified a total of 42 native phosphorylation sites in the Na_V_1.5 channel protein purified from mouse left ventricles, of which 22 are novel. Seventeen of these sites were also found to be phosphorylated in heterologously-expressed Na_V_1.5 channels (9), suggesting that about half of this phosphorylation pattern is conserved among species (mouse and human) and cellular systems (native channels in left ventricles and recombinant channels in HEK-293 cells), whereas the other half may be associated with more specific and/or localized regulation. Among the sites identified, only six were previously suggested to be the targets for specific kinases using *in silico* and/or *in vitro* analyses: S36 and S525 were attributed to the regulation by PKA, S484 and S664 were assigned to the Serum- and Glucocorticoid-inducible Kinase 3 (SGK3), and S516 and S571 were ascribed to CaMKII (reviewed in (2)). In marked contrast, several previously described phosphorylation sites were not detected in the present study, including the PKA-dependent S528, the CaMKII-associated T594, the Protein Kinase C (PKC)-dependent S1503, the Adenosine Monophosphate-activated Protein Kinase (AMPK)-dependent T101 (38), and the six Fyn-dependent tyrosines (39, 40).

Strikingly, and consistent with previous studies from our laboratory (7, 8) and the Bers group (9), the results obtained and presented here again revealed that the first intracellular linker loop of Na_V_1.5 is a hotspot for phosphorylation, with a total of 21 sites identified. Comparisons of the relative abundances of the phosphopeptides identified three highly abundant (and highly phosphorylated) clusters of phosphorylation sites in the first intracellular linker loop of Na_V_1.5 in mouse left ventricles. The simplest interpretation of this finding is that these three phosphorylation clusters, at positions S457 to S460, S483 to T486, and S664 to S671, are likely involved in regulating the basal and/or gating properties of native cardiac Na_V_1.5 channels. Conversely, the other phosphorylation sites, with lower stoichiometries, may play spatially- or temporally-distinct roles in the physiological or more pathophysiological regulation of channel expression or gating. This suggestion is highlighted for residue S671, for example, which is substantially (10-fold) less phosphorylated than the nearby S664 and S667 residues in WT mouse left ventricles, but is (2-fold) upregulated in TAC left ventricles. Remarkably, this mass spectrometry analysis also revealed that the vast majority of identified phosphorylation sites (at least 26) are clustered, suggesting concomitant phosphorylation and roles in regulating channel expression and/or function. Unexpectedly, however, except for S664, S667 and S671, no apparent effects of phosphomimetic or phosphosilent mutations were observed on heterologously-expressed (in HEK-293 cells) Na_V_1.5 current densities or biophysical properties, suggesting a greater complexity than anticipated in the mechanisms contributing to phosphorylation-dependent regulation of Na_V_1.5 channels.

### Phosphorylation at S664 and S667 shifts the voltage-dependence of Na_V_1.5 channel activation towards hyperpolarized potentials

The electrophysiological analyses presented here identified key roles of S664 and S667 in regulating the voltage-dependence of Na_V_1.5 channel activation. Indeed, the data demonstrate that the voltage-dependence of activation of quadruple phosphosilent channels at positions S664-671 is shifted towards depolarized potentials, compared to WT channels, whereas phosphomimetic channels display a smaller shift. These findings are consistent with WT channels being phosphorylated at S664 and S667 in HEK-293 cells, as previously reported (9), and suggest that disruption of phosphorylation at these sites impact channel gating. Confounding this simple interpretation of the data is the fact that glutamate substitution only partially mimics phosphorylation.

Further analyses of the roles of each of the four phosphorylation sites in this cluster revealed the specific involvement of S664 and S667 in regulating gating, whereas modifying T670 or S671 was without effects. Single glutamate mutations at S664 and S667, however, produce the same effects as the single phosphosilent channels. These findings could be attributed to the fact that the side chain of the glutamate only has a single negative charge and a small hydrated shell, which is quite distinct from the covalently attached phosphate group characterized by a doubly negative charge and a large hydrated shell (41). It is likely, therefore, that one glutamate (in single phosphomimetic channels) is not sufficient to mimic phosphorylation at this locus, and that two glutamates (in the quadruple phosphomimetic channel) only partially mimic phosphorylation. The fact that the shifts induced by the single phosphosilent mutations are half the shift generated by the quadruple mutation further supports this hypothesis, and suggests that regulation involving these two sites is cumulative and most likely concomitant. Nevertheless, further investigations, aimed at demonstrating the role of phosphorylation, rather than any other structural determinants associated with this locus, are certainly warranted. In this regard, our findings are also in accordance with previous data reporting the role of SGK3 in shifting the voltage-dependence of channel activation towards more hyperpolarized potentials in *Xenopus* oocytes, whereas the opposite effect was observed with the Na_V_1.5-S664A phosphosilent channel (42). Although the involvement of SGK3 and S664 in a shared regulation was not directly shown in this previous study, it is tempting to speculate that SGK3 may constitute the kinase phosphorylating S664 and S667 and mediating this regulation.

The effects of phosphorylation were also analyzed using all-atom simulations approach to determine how the introduction of negative charges affects the conformational ensemble of the segments containing the phosphorylation clusters identified by mass spectrometry. These simulations demonstrate that the introduction of negative charges at positions S497-S499 and S664-671 could expand the structure of the containing segments, whereas no effects are likely with the segments containing the S457-460 and S483-486 phosphorylation clusters. Furthermore, for both of the affected segments, the expansion likely gradually increases with the cumulative addition of charges. Interestingly, the simulation findings are consistent with the additive roles of S664 and S667 in regulating the voltage-dependence of channel activation observed in the electrophysiological analyses. Consistent with the proximity of the S664-671 phosphorylation cluster to the DII voltage-sensing domain (DII-VSD) of Na_V_1.5, which is tightly linked to channel activation (43), our findings suggest that phosphorylation at S664 and S667 regulates channel activation through the expansion of the C-terminal extremity of the first intracellular linker loop of the channel. However, no effects on channel gating were observed with the S497-499 phosphomimetic mutant, even though the simulation showed an effect on its ability to expand. This result suggests that the expansion of this segment, which is more distal to the DII-VSD, does not regulate channel gating.

### Phosphorylation at S671 decreases Na_V_1.5 channel cell surface expression and peak Na^+^ current density

The functional analyses also demonstrate that mimicking phosphorylation at S671 decreases the expression of the Na_V_1.5 protein at the cell surface, as well as peak Na^+^ current density in HEK-293 cells. These results suggest that S671 is not phosphorylated in HEK-293 cells, which is in agreement with the previously published mass spectrometric analyses (9). While the phosphomimetic mutation greatly decreases the cell surface expression of Na_V_1.5, the phosphosilent mutation also reduces Na_V_1.5 surface expression, albeit to a much smaller extent. These confounding results suggest that the regulation mediated by this locus highly depends on structural changes, and that the phosphomimetic mutation affects the cell surface expression of the channel in part through a change in the structure of the locus. One could further suggest that the greater effect of the phosphomimetic channel may be caused by additional attributes common to the phosphate group and the glutamate side chain. Together, therefore, these findings highlight the novel role of this locus, and of phosphorylation at this site, in regulating the cell surface expression of Na_V_1.5 channels.

Interestingly, the mass spectrometric analyses also revealed that phosphorylation at this site is increased in the left ventricles of TAC mice, suggesting a role in mediating the Na_V_ channel defects associated with heart failure. Because previous studies have suggested that CaMKII-dependent phosphorylation of Na_V_1.5 may constitute one of the molecular mechanisms mediating the increased late Na^+^ current in heart failure (10–15), this finding prompted us to examine the late Na^+^ current generated by the phosphosilent and phosphomimetic Na_V_1.5 mutants at position-671. Our results herein appeared negative, although it cannot be excluded that this regulation may require a specific molecular and cellular environment which is not recapitulated in HEK-293 cells. Additionally, and to our surprise, no changes in phosphorylation at S571 were observed in our TAC model, in contrast with previous findings in nonischemic human heart failure (12) and in several animal models of heart disease (11,12,14). These seemingly disparate findings may reflect technical and/or experimental differences, including differences in the models used and/or stages of disease.

The results presented here raise the interesting and novel possibility that increased phosphorylation at S671 participates in decreasing the peak Na^+^ current often observed in heart failure. Consistent with this suggestion, a recent study by the Remme group, using superresolution microscopy, showed a reduction in the size of Na_V_1.5 clusters in TAC ventricular myocytes without any changes in Na_V_1.5 transcript or total protein expression (29). Although further studies will be required to determine directly whether these observations are causally linked to increased phosphorylation at S671, the results here provide new hints towards understanding the molecular basis of the decreased peak Na^+^ current in heart failure.

Altogether, the results presented here demonstrate that native mouse ventricular Na_V_1.5 is highly phosphorylated, and that the mechanisms mediating the phosphorylation-dependent regulation of Na_V_1.5- encoded channels are site-specific, complex, dynamic, and lead to diverse physiological and/or pathological consequences on both channel gating and expression.

## Materials and methods

### Statement on the use of murine tissue

All investigations conformed to directive 2010/63/EU of the European Parliament, to the Guide for the Care and Use of Laboratory Animals published by the US National Institutes of Health (NIH Publication No. 85-23, revised 1985) and to local institutional guidelines.

### Animal model of heart failure

Heart failure was induced by transverse aortic constriction (TAC) as described previously (14). Eight-week-old male C57/BL6J mice were anesthetized using intraperitoneal injections of medetomidine (0.5 mg/kg), midazolam (5 mg/kg) and fentanyl (0.05 mg/kg body weight). A horizontal incision (1-1.5 cm) at the jugulum was used to display the transverse aorta, and a 27-gauge needle was tied against the aorta using a 6.0 non-absorbable suture. After removal of the 27-gauge needle, the skin was closed, and the mice were kept on a heating plate until recovered from the anesthesia. Sham animals underwent the same procedure except for the banding of the transverse aorta. At the end of the surgery, anesthesia was antagonized using intraperitoneal injections of atipamezol (2.5 mg/kg), flumazenil (0.5 mg/kg) and buprenorphine (0.1 mg/kg body weight). For analgesia, metamizole (1.33 mg/ml) was added to the drinking water 2 days before surgery, and supplied for 7 days after operation. In addition, buprenorphine (60 µg/kg body weight) was administered s.c. 1 hr before surgery. A TAC with a mean gradient of less than 5 mmHg was deemed insufficient to induce heart failure and, if observed, the animal was excluded from later analysis. Mice were sacrificed 5 weeks after TAC by cervical dislocation, and left ventricles were harvested, flash-frozen and stored for further analyses.

### Mouse echocardiography

Transthoracic echocardiography was performed blinded before and 5 weeks after TAC using a Vevo3100 system (VisualSonics, Toronto, Canada) equipped with a 30-MHz center frequency transducer, as described previously (14). The animals were initially anesthetized with 3% isoflurane, while temperature-, respiration-, and electrocardiogram-controlled anesthesia was maintained with 1.5% isoflurane. Two-dimensional cine loops with frame rates of >200 frames/sec of a long axis view and a short axis view at mid-level of the papillary muscles, as well as M-mode loops of the short axis view were recorded. Thicknesses of the anterior (LVAW) and posterior (LVPW) walls of the left ventricle, the inner diameter of the left ventricle (LVID), and the area of the left ventricular cavity were measured in systole (s) and diastole (d) from the short axis view according to standard procedures (44). Maximal left ventricular length was measured from the long axis view. Systolic and diastolic left ventricular volumes (LV vol) were calculated using the area-length method, and the ejection fraction (EF) was derived. Left ventricular mass (LVM) was calculated from anterior and posterior wall thicknesses using Vevo LAB Software (VisualSonics). PW Doppler ultrasound was used to assess mean gradients (MG) 3 days after the TAC procedure.

### Immunoprecipitation of Na_V_ channel complexes

Flash-frozen left ventricles from 4 Sham and 5 TAC mice were homogenized individually in ice-cold lysis buffer containing 20 mM HEPES (pH 7.4), 150 mM NaCl, 0.5% amidosulfobetaine, 1X complete protease inhibitor cocktail tablet, 1 mM phenylmethylsulfonyl fluoride (PMSF), 0.7 μg/ml pepstatin A (Thermo Fisher Scientific, Waltham, MA) and 1X Halt phosphatase inhibitor cocktail (Thermo Fisher Scientific) as described previously (8). All reagents were from Sigma (Saint Louis, MO) unless otherwise noted. After 15-min rotation at 4°C, 8 mg of the soluble protein fractions were pre-cleared with 200 μL of protein G-magnetic Dynabeads (Thermo Fisher Scientific) for 1 hr, and subsequently used for immunoprecipitations (IP) with 48 μg of an anti-Na_V_PAN mouse monoclonal antibody (mαNa_V_PAN, Sigma, #S8809), raised against the SP19 epitope (45) located in the third intracellular linker loop and common to all Na_V_ channel pore-forming subunits. Prior to the IP, antibodies were cross-linked to 200 μl of protein G-magnetic Dynabeads using 20 mM dimethyl pimelimidate (Thermo Fisher Scientific) (46). Protein samples and antibody-coupled beads were mixed for 2 hrs at 4°C. Magnetic beads were then collected, washed rapidly four times with ice-cold lysis buffer, and isolated protein complexes were eluted from the beads in 1X SDS sample buffer (Bio-Rad Laboratories, Hercules, CA) at 60°C for 10 min. Ninety-nine percent of the immunoprecipitated mouse left ventricular Na_V_ channel protein complexes were analyzed by MS, and the remaining one percent was used to verify IP yields by western blotting using a rabbit polyclonal anti-Na_V_1.5 antibody (RbαNa_V_1.5, 1:1000, Alomone labs, Jerusalem, Israel, #ASC-005).

### Peptide preparation and isobaric labeling for LC-MS

The IP eluates were thawed on ice, reduced, and denatured by heating for 10 min at 95°C. The Cys residues were alkylated with iodoacetamide (10 mM) for 45 min at room temperature in the dark. The peptides were prepared using a modification (47) of the filter-aided sample preparation method (48). After the addition of 300 µL of 100 mM Tris buffer (pH 8.5) containing 8 M urea (UT) and vortexing, the samples were transferred to YM-30 filter units (Millipore, MRCF0R030) and spun for 14 min at 10,000 rcf (Eppendorf, Model No. 5424). The filters were washed with 200 µl of UT buffer, and the spin-wash cycle was repeated twice. The samples were then exchanged into digest buffer with the addition of 200 µL of 50 mM Tris buffer, pH 8.0, followed by centrifugation (10,000 rcf for 10 min). After transferring the upper filter units to new collection tubes, 80 µL of digest buffer was added, and the samples were digested with trypsin (1 µg) for 4 h at 37°C. The digestion was continued overnight after adding another aliquot of trypsin. The filter units were then spun for 10 min (10,000 rcf) in an Eppendorf microcentrifuge. The filter was washed with 50 µL of Tris buffer (100 mM, pH 8.0), followed by centrifugation. The digests were extracted three times with 1 ml of ethyl acetate, and acidified to 1% trifluoroacetic acid (TFA) using a 50% aqueous solution. The pH was < 2.0 by checking with pH paper. The solid phase extraction of the peptides was performed using porous graphite carbon micro-tips (49). The peptides were eluted with 60% acetonitrile in 0.1% TFA, and pooled for drying in a Speed-Vac (Thermo Scientific, Model No. Savant DNA 120 concentrator) after adding TFA to 5%. The peptides were dissolved in 20 µL of 1% acetonitrile in water. An aliquot (10%) was removed for quantification using the Pierce Quantitative Fluorometric Peptide Assay kit (Thermo Scientific, Cat. No. 23290). The remainder of the peptides from each IP samples (∼0.5-3.5 µg) and 1.16 µg of reference pool peptide were transferred into a new 0.5 mL Eppendorf tube, dried in the Speed-Vac, and dissolved in 12 µL of HEPES buffer (100 mM, pH 8.0, Sigma, H3537).

The samples were labeled with tandem mass tag reagents (TMT11, Thermo Scientific) according to manufacturer’s protocol. The labeled samples were pooled, dried, and resuspended in 120 µL of 1% formic acid (FA). The TMT11 labeled sample was desalted as described above for the unlabeled peptides. The eluates were transferred to autosampler vials (Sun-Sri, Cat. No. 200046), dried, and stored at −80°C for capillary liquid chromatography interfaced to a mass spectrometer (nano-LC-MS).

### Nano-LC-MS

The samples in formic acid (1%) were loaded (2.5 µL) onto a 75 µm i.d. × 50 cm Acclaim^®^ PepMap 100 C18 RSLC column (Thermo-Fisher Scientific) on an EASY *nano-*LC (Thermo Fisher Scientific). The column was equilibrated using constant pressure (700 bar) with 20 μL of solvent A (0.1% FA). The peptides were eluted using the following gradient program with a flow rate of 300 nL/min and using solvents A and B (acetonitrile with 0.1% FA): solvent A containing 5% B for 1 min, increased to 25% B over 87 min, to 35% B over 40 min, to 70% B in 6 min and constant 70% B for 6 min, to 95% B over 2 min and constant 95% B for 18 min. The data were acquired in data-dependent acquisition (DDA) mode. The MS1 scans were acquired with the Orbitrap™ mass analyzer over *m/z* = 375 to 1500 and resolution set to 70,000. Twelve data-dependent high-energy collisional dissociation spectra (MS2) were acquired from each MS1 scan with a mass resolving power set to 35,000, a range of *m/z* = 100 - 1500, an isolation width of 2 Th, and a normalized collision energy setting of 32%. The maximum injection time was 60 ms for parent-ion analysis and 120 ms for product-ion analysis. The ions that were selected for MS2 were dynamically excluded for 20 sec. The automatic gain control (AGC) was set at a target value of 3e6 ions for MS1 scans and 1e5 ions for MS2. Peptide ions with charge states of one or ≥ 7 were excluded for higher-energy collision-induced dissociation (HCD) acquisition.

### MS data analysis

Peptide identification from raw MS data was performed using PEAKS Studio 8.5 (Bioinformatics Solutions Inc., Waterloo, Canada) (50). The Uni-mouse-Reference-20131008 protein database was used for spectral matching. The precursor and product ion mass tolerances were set to 20 ppm and 0.05 Da, respectively, and the enzyme cleavage specificity was set to trypsin, with a maximum of three missed cleavages allowed. Carbamidomethylation (Cys) and TMT tags (Lys and/or peptide N-terminus) were treated as fixed modifications, while oxidation (Met), pyro-glutamination (Gln), deamidation (Asn and/or Gln), methylation (Lys and/or Arg), dimethylation (Lys and/or Arg), acetylation (Lys) and phosphorylation (Ser, Thr and/or Tyr) were considered variable modifications. The definitive annotation of each Na_V_1.5 phosphopeptide-spectrum match was obtained by manual verification and interpretation. The phosphorylation site assignments were based on the presence or absence of the unphosphorylated and phosphorylated b- and y-ions flanking the site(s) of phosphorylation, ions referred to as site-discriminating ions throughout this study. When site-discriminating ions were not all detected, the assignment of phosphorylation sites was narrowed down to several possibilities by elimination (for example, pS1056 and/or pT1058). Representative MS/MS spectra, PEAKS -10lgP scores, mass errors of parent ions (in ppm) and charge state confirmations of site-discriminating b- and y-ions are presented in **Table 2, Table 2 - Table Supplement 1 & Figure Supplement 1**.

The protein and peptide relative abundances in TAC, *versus* Sham, mαNa_V_PAN-IPs were calculated using quantification of TMT reporter ions. Reporter ion intensities in each TMT channel were normalized to the mean reporter ion intensities of Na_V_1.5-derived peptides (normalization to spike) to correct for differences in IP yields and technical variabilities. Normalization factors are presented in **Figure 1 - Figure Supplement 1B**. Quantification values of each peptide-spectrum match were exported into Excel, and the mean peptide abundance ratios were calculated from the abundance ratios of all manually verified peptide-spectrum matches assigning to the phosphorylation site(s) of interest. Label-free quantitative analysis of the areas of extracted MS1 chromatograms of phosphorylated and non-phosphorylated peptide ions covering the phosphorylation site(s) of interest was used to evaluate the proportion of phosphorylated to non-phosphorylated peptides at each position, as well as the relative abundances of phosphopeptides.

### Plasmids

The Na_V_1.5 phosphomutant constructs were generated by mutating the serine(s)/threonine(s) to alanine(s) (A) or glutamate(s) (E) by site-directed mutagenesis of a pCI-Na_V_1.5 plasmid containing the human Na_V_1.5 hH1C cDNA (30) (NCBI Reference Sequence NM_000335) using the QuikChange II XL Site-Directed Mutagenesis kit (Agilent, Sant Clara, CA) or the Q5 Site-Directed Mutagenesis kit (New England Biolabs, Ipswich, MA). The mutated constructs were then digested with restriction endonucleases to excise the mutated fragments, which were then subcloned into the original pCI-Na_V_1.5 plasmid. The human Na_V_β1 (NM_001037, a gift from A. L. George) cDNAs was subcloned into pRc/CMV. All constructs were sequenced to ensure that no unintentional mutations were introduced.

### Culture and transient transfections

Human Embryonic Kidney 293 (HEK-293) cells were maintained in Dulbecco’s Modified Eagle’s Medium (DMEM, Thermo Fisher Scientific), supplemented with 10% fetal bovine serum, 100 U/ml penicillin and 100 μg/ml streptomycin, in 37°C, 5% CO_2_: 95% air incubator. Cells were transiently transfected at 70-80% confluence in 35 mm dishes with 0.6 μg of the WT or phosphomutant Na_V_1.5 plasmid and 1.2 μg of the Na_V_β1 plasmid using 2 μL of Lipofectamine 2000 (Thermo Fisher Scientific) following the manufacturer’s instructions. For whole-cell recordings, transfections also contained 0.2 μg of the pEGFP plasmid (Enhanced Green Fluorescent Protein plasmid, Clontech), and EGFP expression served as a marker of transfection. The absolute amounts of the various constructs were calculated and the empty pcDNA3.1 plasmid was used as a filler plasmid to keep the total DNA constant at 2 μg in each transfection.

### Electrophysiological recordings

Whole-cell Na_V_ currents were recorded at room temperature from transiently transfected HEK-293 cells using an Axopatch 200A amplifier (Axon Instruments, Molecular Devices, San Jose, CA) 48 hours after transfection. Voltage-clamp protocols were applied using the pClamp 10.2 software package (Axon Instruments) interfaced to the electrophysiological equipment using a Digidata 1440A digitizer (Axon Instruments). Current signals were filtered at 10 kHz prior to digitization at 50 kHz and storage. Patch-clamp pipettes were fabricated from borosilicate glass (OD: 1.5 mm, ID: 0.86 mm, Sutter Instrument, Novato, CA) using a P-97 micropipette puller (Sutter Instrument), coated with wax, and fire-polished to a resistance between 1.5 and 2.5 MΩ when filled with internal solution. The internal solution contained (in mM): NaCl 5, CsF 115, CsCl 20, HEPES 10, EGTA 10 (pH 7.35 with CsOH, ∼300 mosM). The external solution contained (in mM): NaCl 10 (NaCl 20 for analysis of single phosphomutants), CsCl 103, TEA-Cl (tetraethylammonium chloride) 25, HEPES 10, Glucose 5, CaCl_2_ 1, MgCl_2_ 2 (pH 7.4 with CsOH, ∼300 mosM). All chemicals were purchased from Sigma. After establishing the whole-cell configuration, three minutes were allowed to ensure stabilization of voltage-dependence of activation and inactivation properties, at which time 25 ms voltage steps to ± 10 mV from a holding potential (HP) of - 70 mV were applied to allow measurement of whole-cell membrane capacitances, input and series resistances. Only cells with access resistance < 7 MΩ were used, and input resistances were typically > 5 GΩ. After compensation of series resistance (80%), the membrane was held at a HP of -120 mV, and the voltage-clamp protocols were carried out as indicated below. Leak currents were always < 200 pA at HP (−120 mV), and were corrected offline. Cells exhibiting peak current amplitudes < 500 or > 5000 pA were excluded from analyses of biophysical properties because of leak or voltage-clamp issues, respectively, but conserved in analyses of peak current density to avoid bias in evaluation of current densities.

Data were compiled and analyzed using ClampFit 10.2 (Axon Instruments), Microsoft Excel, and Prism (GraphPad Software, San Diego, CA). Whole-cell membrane capacitances (Cm) were determined by analyzing the decays of capacitive transients elicited by brief (25 ms) voltage steps to ± 10 mV from the HP (−70 mV). Input resistances were calculated from the steady-state currents elicited by the same ±10 mV steps (from the HP). Series resistances were calculated by dividing the decay time constants of the capacitive transients (fitted with single exponentials) by the Cm. To determine peak Na^+^ current-voltage relationships, currents were elicited by 50-ms depolarizing pulses to potentials ranging from −80 to +40 mV (presented at 5-s intervals in 5-mV increments) from a HP of -120 mV. Peak current amplitudes were defined as the maximal currents evoked at each voltage. Current amplitudes were leak-corrected, normalized to the Cm, and current densities are presented.

To analyze voltage-dependence of current activation properties, conductances (G) were calculated, and conductance-voltage relationships were fitted with the Boltzmann equation G = G_max_ / (1 + exp (- (V_m_ - V_1/2_) / k)), in which V_1/2_ is the membrane potential of half-activation and k is the slope factor. The time courses of inactivation of macroscopic currents were fitted with bi-exponential functions, I(*t*) = A_fast_ x exp (-*t*/τ_fast_) + A_slow_ x exp (-*t*/τ_slow_) + A_0_, in which A_fast_ and A_slow_ are the amplitudes of the fast and slow inactivating current components, respectively, and τ_fast_ and τ_slow_ are the decay time constants of A_fast_ and A_slow_, respectively. A standard two-pulse protocol was used to examine the voltage-dependences of steady-state inactivation. From a HP of -120 mV, 1-s conditioning pulses to potentials ranging from -120 to -45 mV (in 5-mV increments) were followed by 20-ms test depolarizations to -20 mV (interpulse intervals were 5-s). Current amplitudes evoked from each conditioning voltage were measured and normalized to the maximal current (I_max_) evoked from -120 mV, and normalized currents were plotted as a function of the conditioning voltage. The resulting steady-state inactivation curves were fitted with the Boltzmann equation I = I_max_ / (1 + exp ((V_m_ - V_1/2_) / k)), in which V_1/2_ is the membrane potential of half-inactivation and k is the slope factor. To examine the rates of recovery from inactivation, a three-pulse protocol was used. Cells were first depolarized to -20 mV (from a HP of -120 mV) to inactivate the channels, and subsequently repolarized to -120 mV for varying times (ranging from 1 to 200 ms), followed by test depolarizations to -20 mV to assess the extent of recovery (interpulse intervals were 5-s). The current amplitudes at -20 mV, measured following each recovery period, were normalized to the maximal current amplitude and plotted as function of the recovery time. The resulting plot was fitted with a single exponential function I(*t*) = A x (1 - exp (-*t* / τ_rec_)) to determine the recovery time constant. For each of these biophysical properties, data from individual cells were first fitted and then averaged.

The currents generated on expression of each phosphosilent and phosphomimetic Na_V_1.5 mutant were recorded and compared to currents generated by the Na_V_1.5-WT construct obtained on the same days of patch-clamp analyses. The densities and properties of Na_V_1.5-WT currents in each data set were similar, and for the sake of clarity, a single representative Na_V_1.5-WT channel data set was chosen and presented in **Figure 3 - Figure Supplement 2 and Table Supplement 1 and Figure 4 - Table Supplement 1**.

In experiments aimed at recording the tetrodotoxin (TTX)-sensitive late Na^+^ current, cells were bathed in external solution containing (in mM): NaCl 120, TEA-Cl 25, HEPES 10, Glucose 5, CaCl_2_ 1, MgCl_2_ 2 (pH 7.4 with CsOH, ∼300 mosM). Repetitive 350-ms test pulses to -20 mV from a HP of -120 mV (at 5-s intervals) were applied to cells to record Na^+^ currents in the absence of TTX. Cells were then superfused locally with the external solution supplemented with 30 μM TTX (Bio-Techne SAS, Rennes, France). Only cells exhibiting peak current amplitudes > 4000 pA were used (those with peak currents < 4000 pA did not show measurable late Na^+^ current), and cells with difference in leak current amplitudes before and after TTX application > 5 pA at -20 mV (calculated from leak currents at -120 mV) were excluded from analyses. TTX-sensitive currents from individual cells were determined by offline digital subtraction of average leak-subtracted currents obtained from 5 recordings in the absence and in the presence of TTX after achieving steady state. The amplitude of TTX-sensitive late Na^+^ current was defined as the steady-state current amplitude (A_0_) obtained by fitting the inactivation decay of macroscopic TTX-sensitive current with the double exponential function I(*t*) = A_fast_ x exp (-*t*/τ_fast_) + A_slow_ x exp (-*t*/τ_slow_) + A_0_. For each cell, the TTX-sensitive late Na^+^ current amplitude was normalized to the peak Na^+^ current amplitude, and expressed as a percentage of the peak Na^+^ current.

### Cell surface biotinylation and western blot analyses

Surface biotinylation of HEK-293 cells was completed as described previously (51). Briefly, cells were incubated with the cleavable EZ-Link Sulfo-NHS-SS-Biotin (0.5 mg/ml, Thermo Fisher Scientific) in ice-cold PBS (pH 7.4) for 30 min at 4°C. Free biotin was quenched with Tris-saline (10 mM Tris (pH 7.4), 120 mM NaCl), and detergent-soluble cell lysates were prepared. Biotinylated cell surface proteins were affinity-purified using NeutrAvidin-conjugated agarose beads (Thermo Fisher Scientific), and purified cell surface proteins were analyzed by western blot using the mαNa_V_PAN antibody (1:2000, Sigma, #S8809), the anti-transferrin receptor mouse monoclonal antibody (TransR, 1:1000, Thermo Fisher Scientific), and the anti-glyceraldehyde 3-phosphate dehydrogenase mouse monoclonal antibody (GAPDH, 1:100000, Santa Cruz Biotechnology). Bound primary antibodies were detected using horseradish peroxidase-conjugated goat anti-mouse secondary antibodies (Cell Signaling Technology, Inc., Danvers, MA), and protein signals were visualized using the SuperSignal West Dura Extended Duration Substrate (Thermo Fisher Scientific). Bands corresponding to Na_V_1.5 were normalized to bands corresponding to TransR from the same sample. Na_V_1.5 phosphomutant protein expression (total or cell surface) is expressed relative to Na_V_1.5-WT protein expression (total or cell surface).

### Molecular simulations

Molecular simulations were performed with the CAMPARI software package (35), using the ABSINTH implicit solvation model (52) and parameters from the OPLS-AA forcefield. Markov Chain Metropolis Monte Carlo moves sampled the conformational space of each protein segment. To mimic a 5 mM NaCl concentration, neutralizing and excess Na^+^ and Cl^-^ ions were modeled explicitly with the protein segments in spherical droplets of (5 x number of residues) Å radius. Ten simulation runs were performed for each sequence construct, and the average of these ten runs was then plotted. The simulations denoted as EV (Excluded Volume) and FRC (Flory Random Coil) are reference models. In the EV limit, the only interactions considered are the pairwise repulsions. In the FRC limit, conformations are constructed by randomly sampling residue-specific backbone dihedral angles. Three EV and three FRC simulation runs were performed for each protein segment, and the averages were plotted.

### Statistical analyses

Results are expressed as means ± SEM. Data were first tested for normality using the D’Agostino and Pearson normality test. Depending on the results of normality tests, statistical analyses were then performed using the Mann-Whitney nonparametric test, the Kruskal-Wallis one-way ANOVA followed by the Dunn’s post-hoc test, or the one-way ANOVA followed by the Dunnett’s post-hoc test, as indicated in Figures and Tables. All these analyses, as well as plots and graphs were performed using Prism (GraphPad Software).

## Acknowledgements

This work was supported by the *Agence Nationale de la Recherche* [ANR-15-CE14-0006-01 and ANR-16-CE92-0013-01 to CM], the Deutsche Forschungsgemeinschaft (DFG) [Ma 1982/5-1 to LSM], and the National Institutes of Health [R01-HL148803 to CM, RVP and JRS; R01-HL034161 and R01-HL142520 to JMN; and 5R01NS056114 to RVP]. The mass spectrometric experiments were performed at the Washington University Proteomics Shared Resource (WU-PSR). The WU-PSR is supported in part by the WU Institute of Clinical and Translational Sciences (NCATS UL1 TR000448), the Mass Spectrometry Research Resource (NIGMS P41 GM103422) and the Siteman Comprehensive Cancer Center Support Grant (NCI P30 CA091842). ML was supported by a *Groupe de Réflexion sur la Recherche Cardiovasculaire-Société Française de Cardiologie* predoctoral fellowship [SFC/GRRC2018]. SB was supported by a Lefoulon-Delalande postdoctoral fellowship. The expert technical assistance of Petra Erdmann-Gilmore, Yiling Mi and Rose Connors is gratefully acknowledged. The content of the research reported is solely the responsibility of the authors and does not necessarily represent the official view of the funding agencies.

## Author Contributions

CM designed the study and wrote the paper. CM, DM, JMN, RRT and LSM designed, performed and/or analyzed the mass spectrometry experiments. ML, SB, AL, CC, PMC, BE, JMN and CM designed, performed and/or analyzed the functional analyses. EW, KMR, RVP and JRS designed, performed and/or analyzed the modeling analyses. SW and LSM provided the Sham/TAC mice and performed mouse echocardiography. All authors reviewed the results and approved the final version of the manuscript.

## Figure Legends

**Figure 1 - Table Supplement 1.**
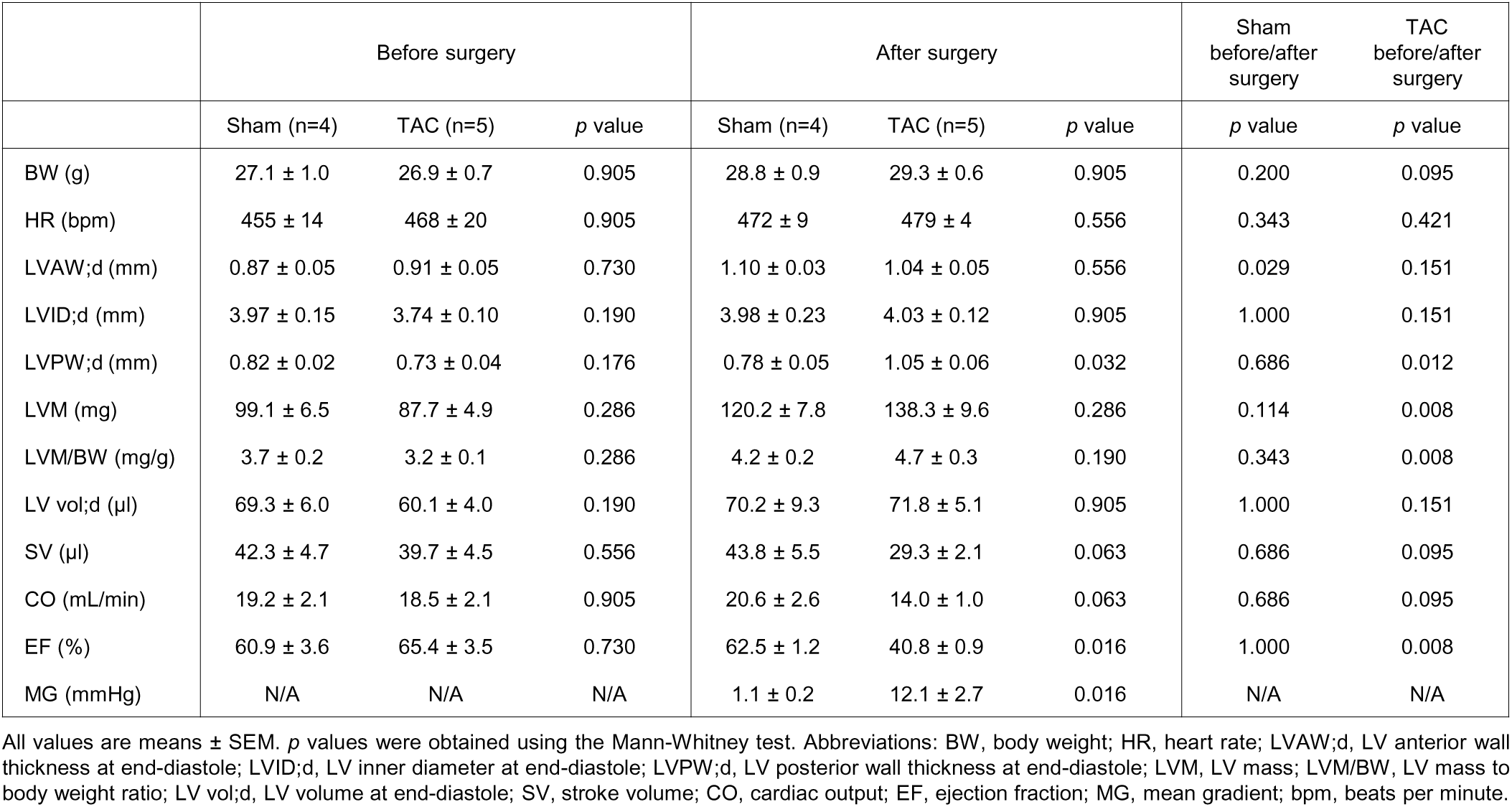
Echocardiographic parameters of Sham and TAC mice before and 5 weeks after surgery

**Figure 1 - Figure Supplement 1.**
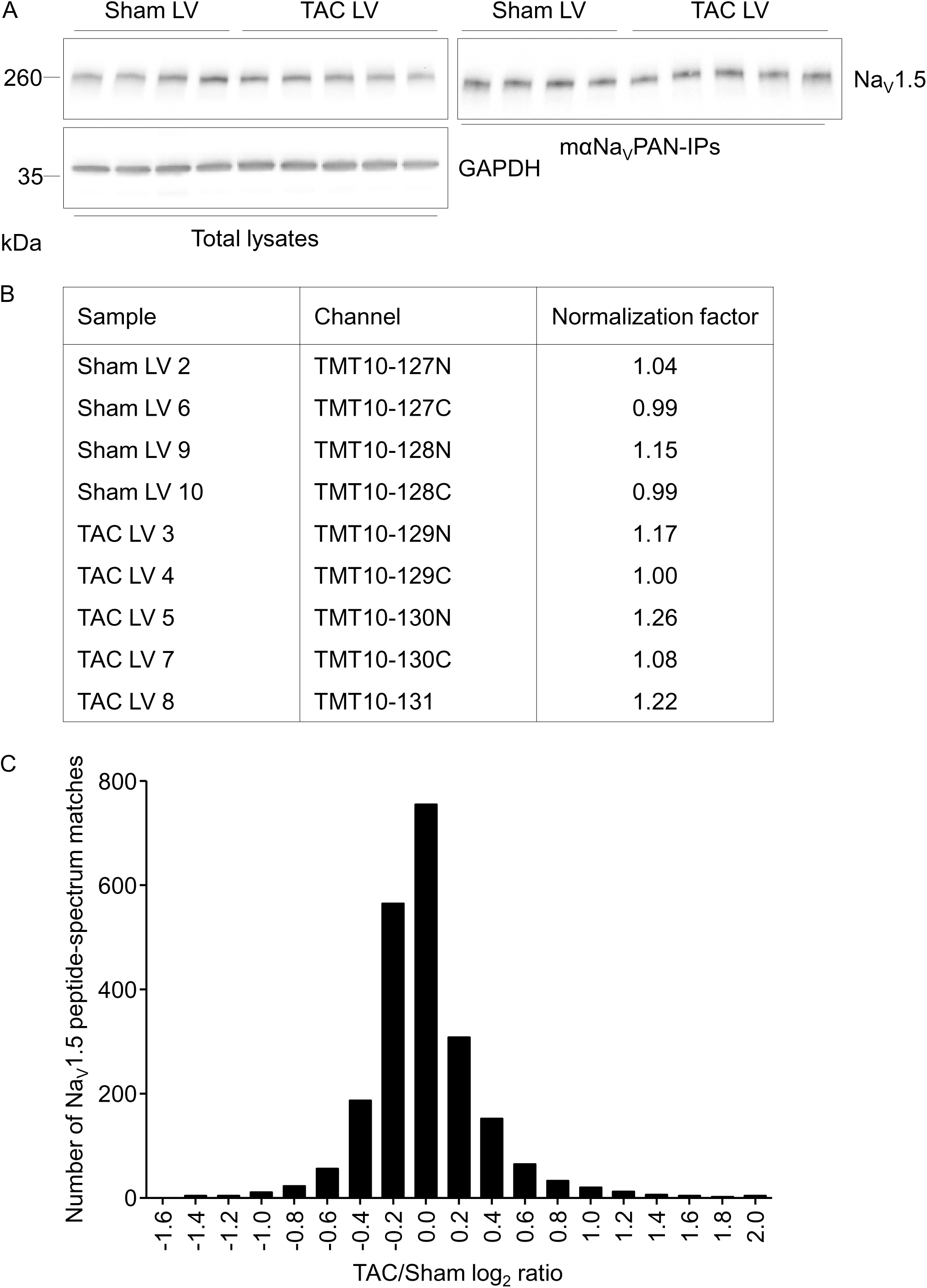
Immunoprecipitation yields and relative quantification of Na_V_1.5 peptide abundances from Sham and TAC mouse left ventricles. (**A**) Representative western blots of total lysates and mαNa_V_PAN-IPs from Sham and TAC left ventricles probed with the anti-Na_V_1.5 rabbit polyclonal (RbαNa_V_1.5) and anti-GAPDH mouse monoclonal antibodies. (**B**) Normalization factors used in MS1 and MS2 analyses to correct for technical variabilities in Na_V_1.5 protein abundance in mαNa_V_PAN-IPs from Sham and TAC left ventricles. (**C**) Distribution of TAC/Sham log_2_ normalized ratios of Na_V_1.5 peptide-spectrum matches. Both biochemical (**A**) and mass spectrometry (**B, C**) analyses of Na_V_1.5 immunoprecipitation yields and peptide relative abundance demonstrate low technical variability.

**Figure 1 - Figure Supplement 2.**
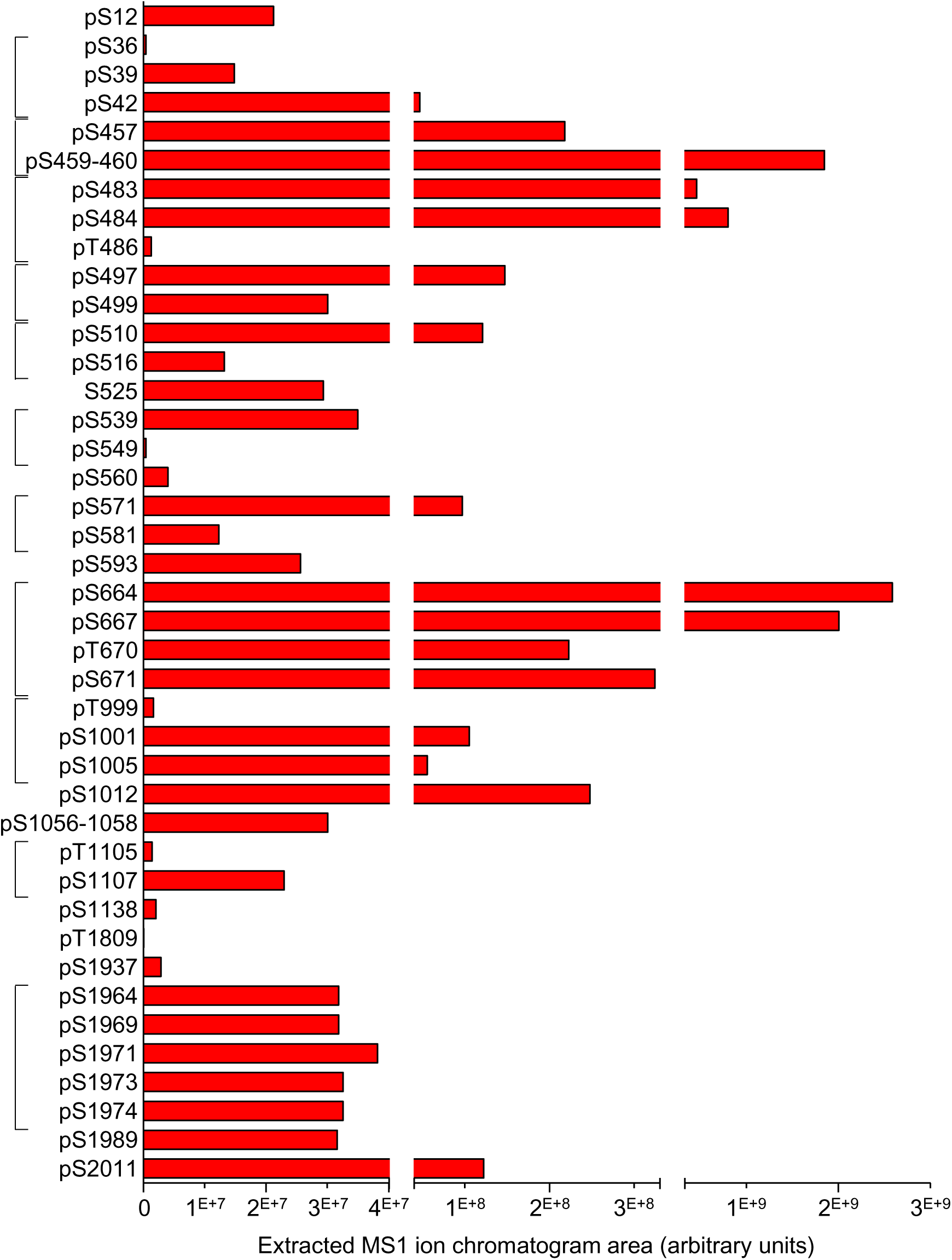
Relative abundances of phosphorylated Na_V_1.5 peptides at indicated phosphorylation site(s) in mαNa_V_PAN-IPs from Sham and TAC mouse left ventricles. Values correspond to the areas of extracted MS1 ion chromatograms corresponding to MS2 spectra assigning phosphorylated peptides at indicated phosphorylation site(s). The brackets indicate the subgroups of phosphorylation sites analyzed in Figure 1B. Independent quantification of S459 and S460 phosphorylated peptides was not possible because localization of the phosphorylation site in most of the phosphorylated peptides could not be discriminated.

**Figure 3 - Table Supplement 1.**
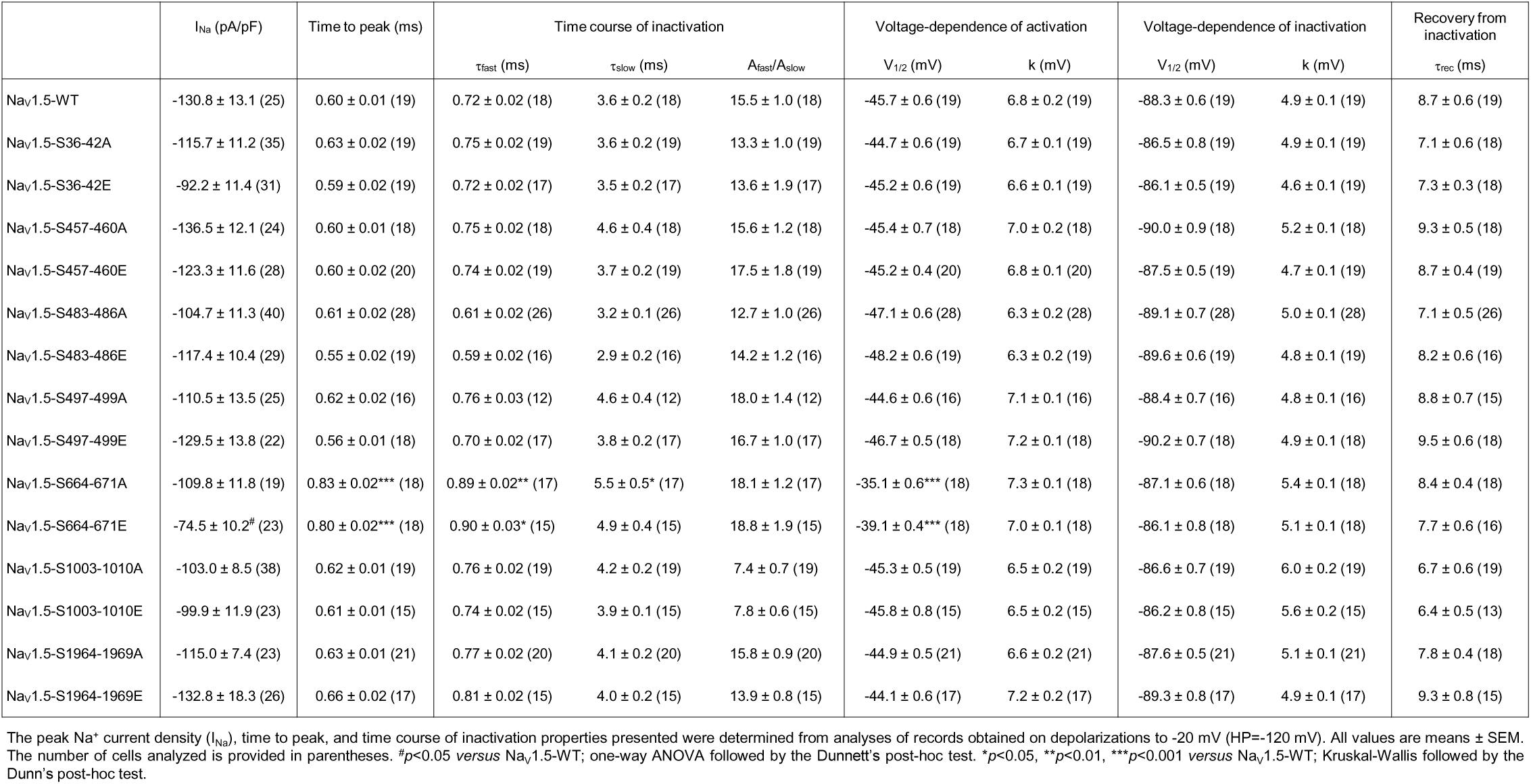
Current densities and properties of NaV1.5 channels mutant for the phosphorylation clusters in transiently transfected HEK293 cells

**Figure 3 - Figure Supplement 1.**
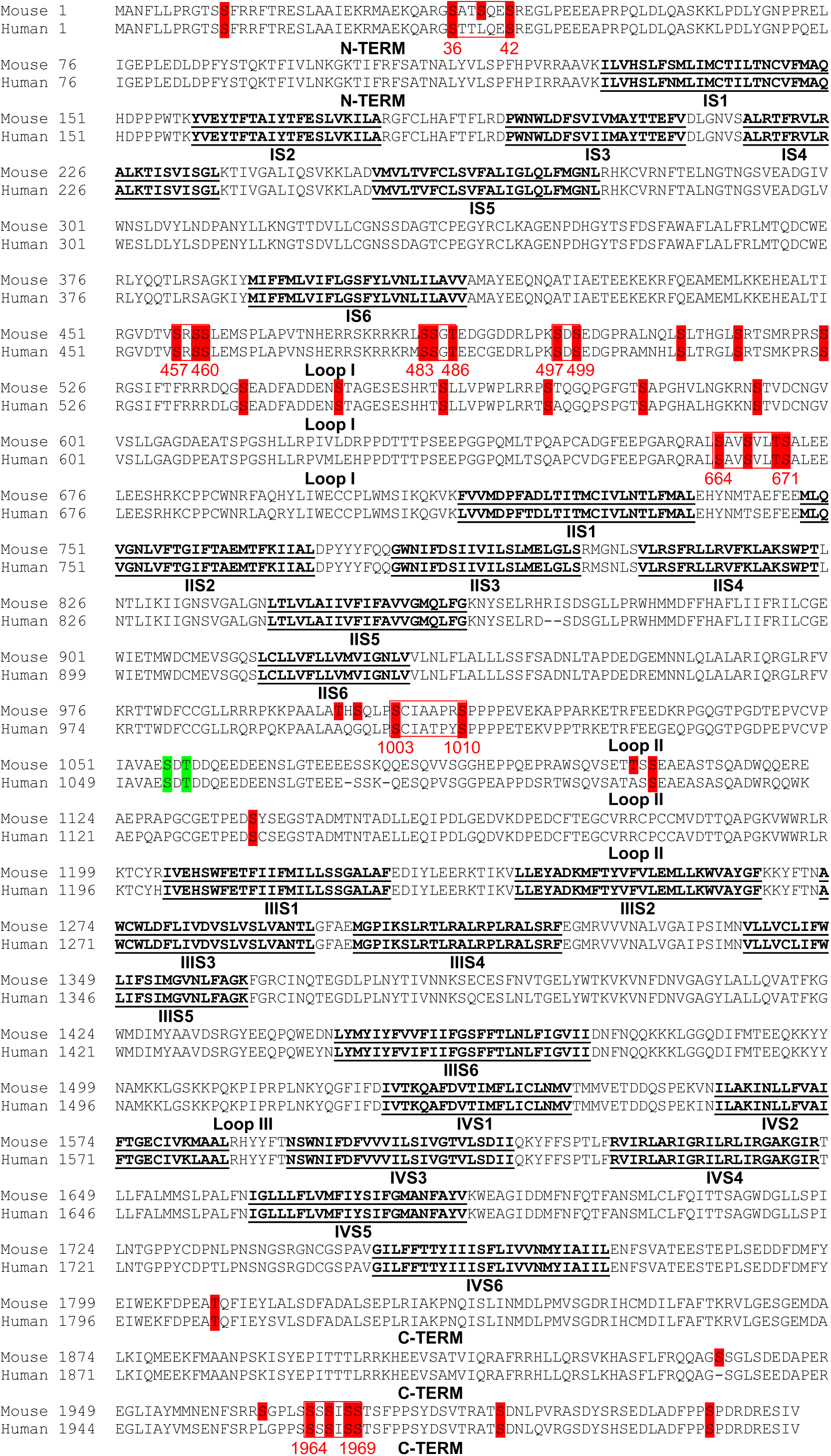
Conservation of phosphorylation sites in mouse and human Na_V_1.5. The mouse (Reference sequence NP_001240789.1) and human (NP_000326.2) Na_V_1.5 sequences are aligned, and phosphorylation sites identified on the mouse sequence and conserved in human are highlighted in red. Two phosphorylation site locations are possible at amino acids S1056-T1058 (in green). Transmembrane segments (S1-S6) in each domain (I-IV) are in bold and underlined in black; loops I, II and III correspond to interdomains I-II, II-III and III-IV, respectively. The seven phosphorylation clusters analyzed electrophysiologically are boxed in red.

**Figure 3 - Figure Supplement 2.**
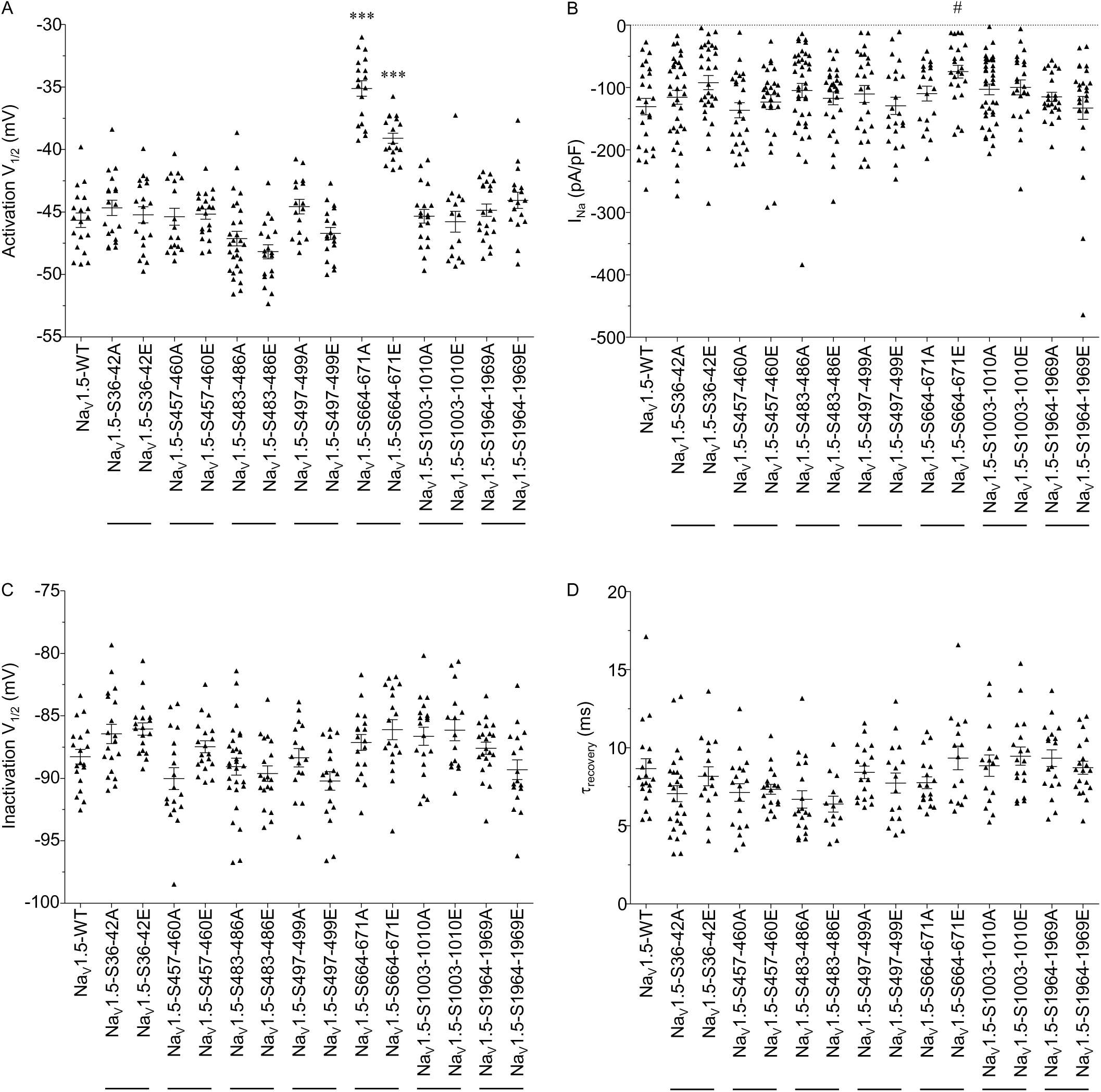
Distributions and mean ± SEM membrane potentials for half-activation (**A**) and half-inactivation (**C**), peak Na^+^ current (I_Na_) densities (**B**), and time constants of recovery from inactivation (**D**) of WT and mutant Na_V_1.5 channels. Currents were recorded as described in the legend to Figure 3. The I_Na_ densities presented were determined from analyses of records obtained on depolarizations to -20 mV (HP=-120 mV). ^#^*p*<0.05 *versus* Na_V_1.5-WT; one-way ANOVA followed by the Dunnett’s post-hoc test. \*\*\**p*<0.001 *versus* Na_V_1.5-WT; Kruskal-Wallis followed by the Dunn’s post-hoc test. Current densities, time- and voltage-dependent properties, as well as statistical comparisons across groups, are provided in **Figure 3 - Table Supplement 1**.

**Figure 4 - Table Supplement 1.**
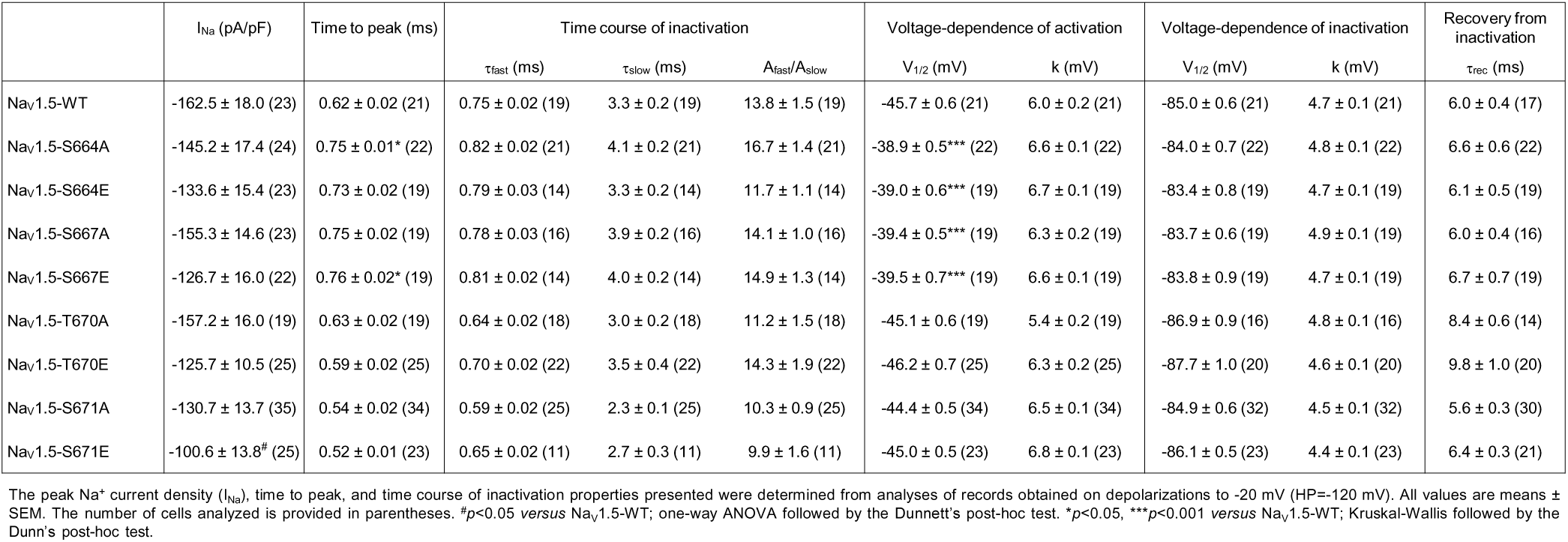
Current densities and properties of NaV1.5 channels mutant for S664, S667, T670 and S671 in transiently transfected HEK293 cells

**Figure 4 - Figure Supplement 1.**
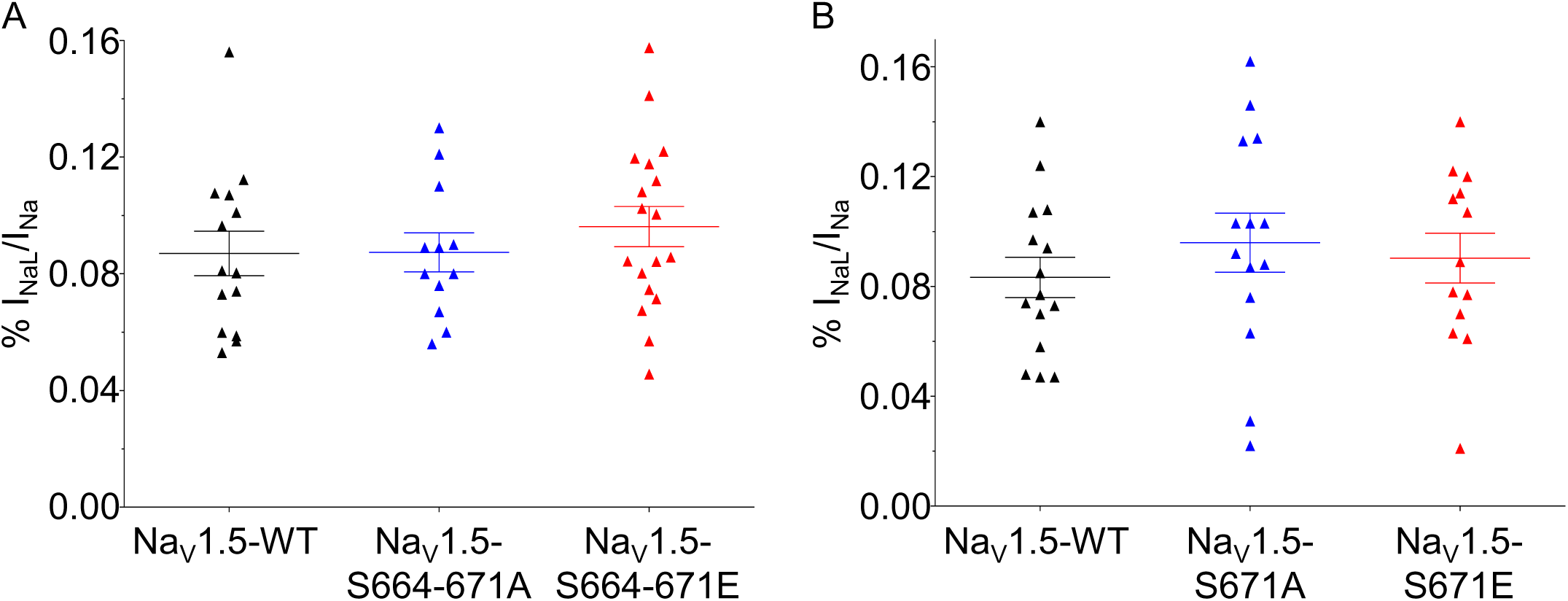
Distributions and mean ± SEM TTX-sensitive late Na^+^ current (I_NaL_) densities of quadruple S664-671 (**A**) and simple S671 (**B**) Na_V_1.5 phosphomutants. TTX-sensitive I_NaL_ were evoked during prolonged depolarizations (350 ms at -20 mV, HP=-120 mV) forty-eight hours after transfection of HEK-293 cells with WT (black), phosphosilent (blue) and phosphomimetic (red) Na_V_1.5 channels and Na_V_β1. No significant differences between mutant and WT channels were observed.

**Table 2 - Figure Supplement 1.**
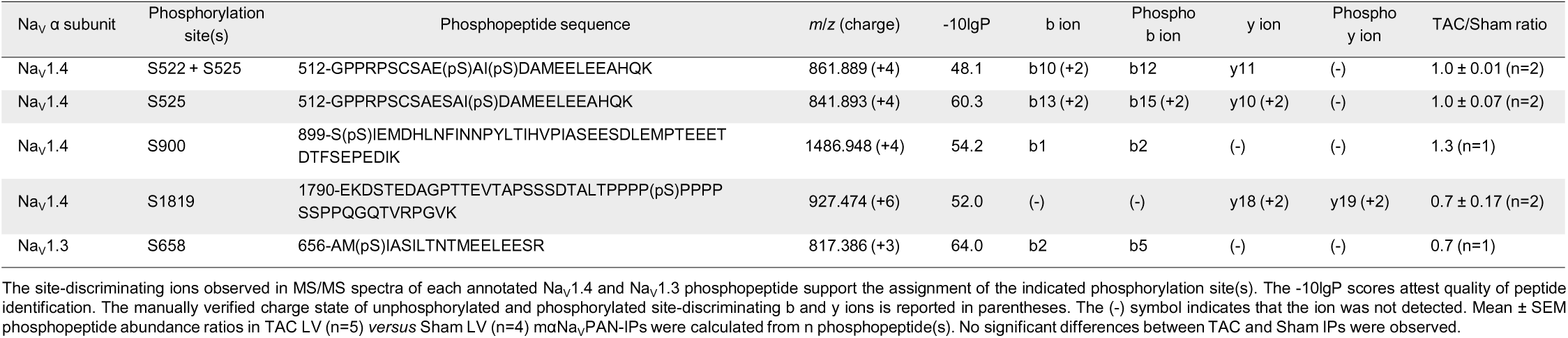
Representative MS/MS spectra of singly or doubly phosphorylated Na_V_1.5, Na_V_1.4 and Na_V_1.3 tryptic peptides (listed in Table 2 **& Table 2 - Table Supplement 1**). The presence of the b- (highlighted in blue) and y- (in red) ion series describing the amino acid sequences, and of the unphosphorylated and phosphorylated site-discriminating ions unambiguously supported the assignments of the indicated phosphorylation site(s). The PEAKS -10lgP peptide scores and the mass errors of parent ions (in ppm) for each phosphopeptide are indicated at the top of each page. The charge state confirmations of site-discriminating ions are presented in Table 2 **& Table 2 - Table Supplement 1**.

**Table 2 - Table Supplement 1.**
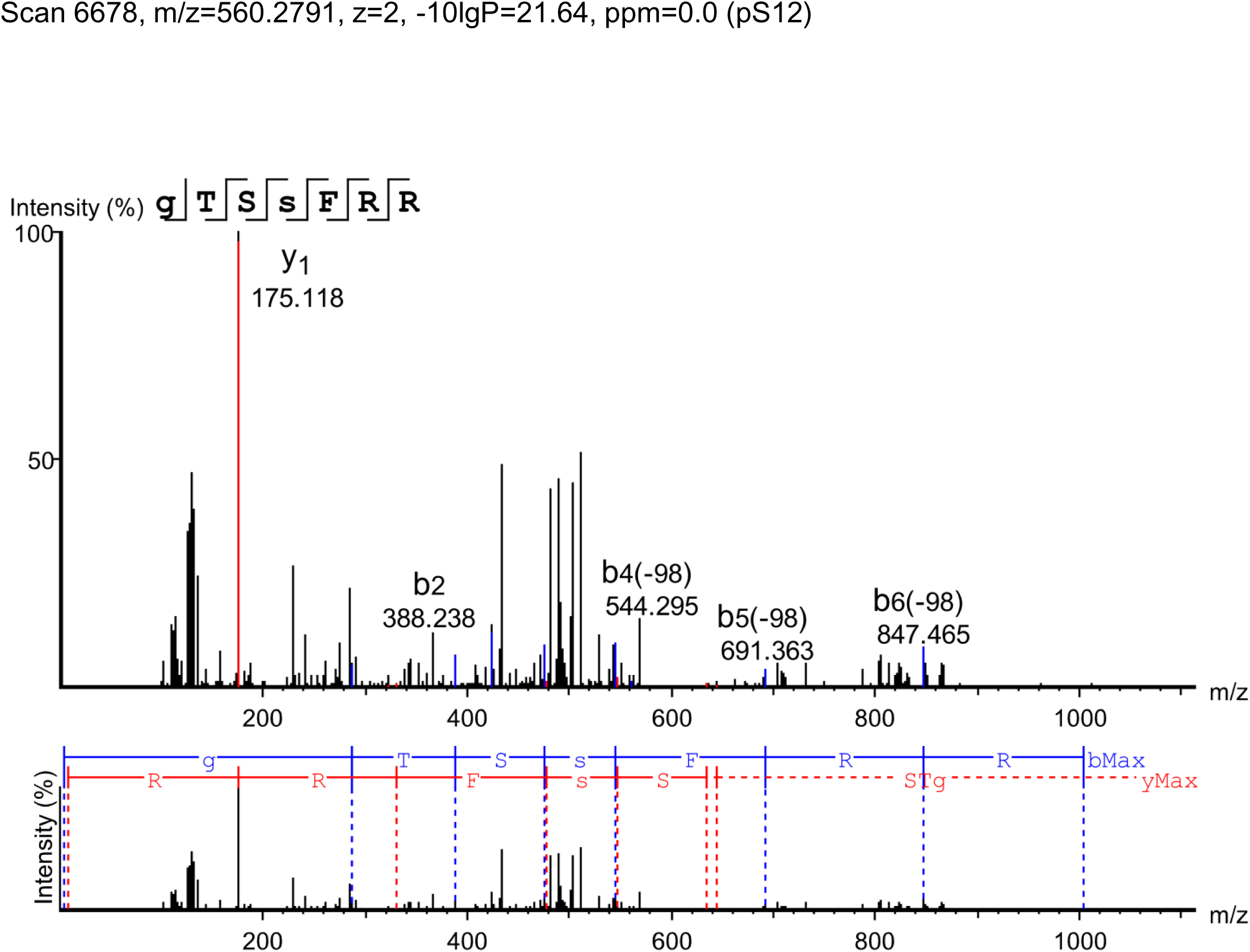
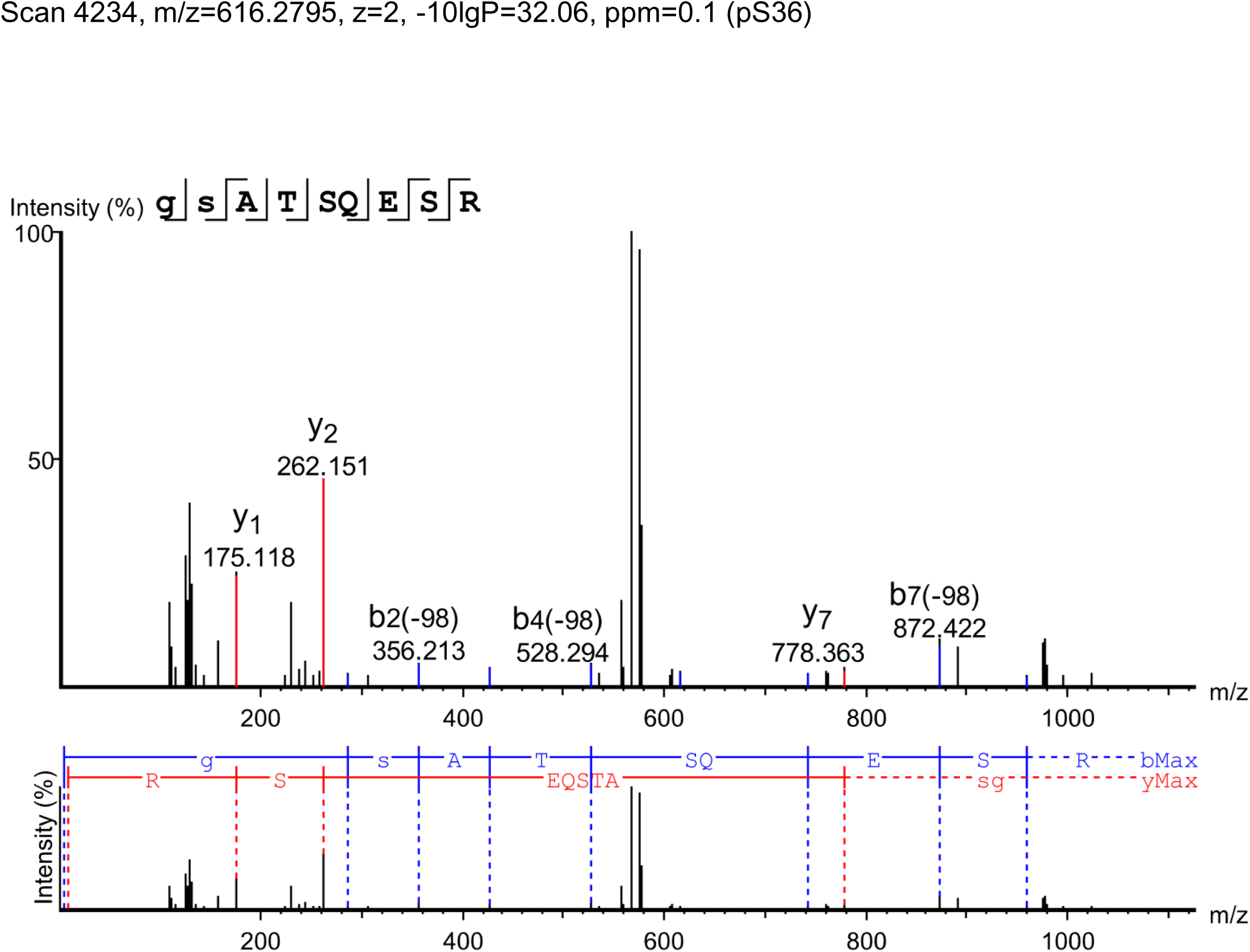
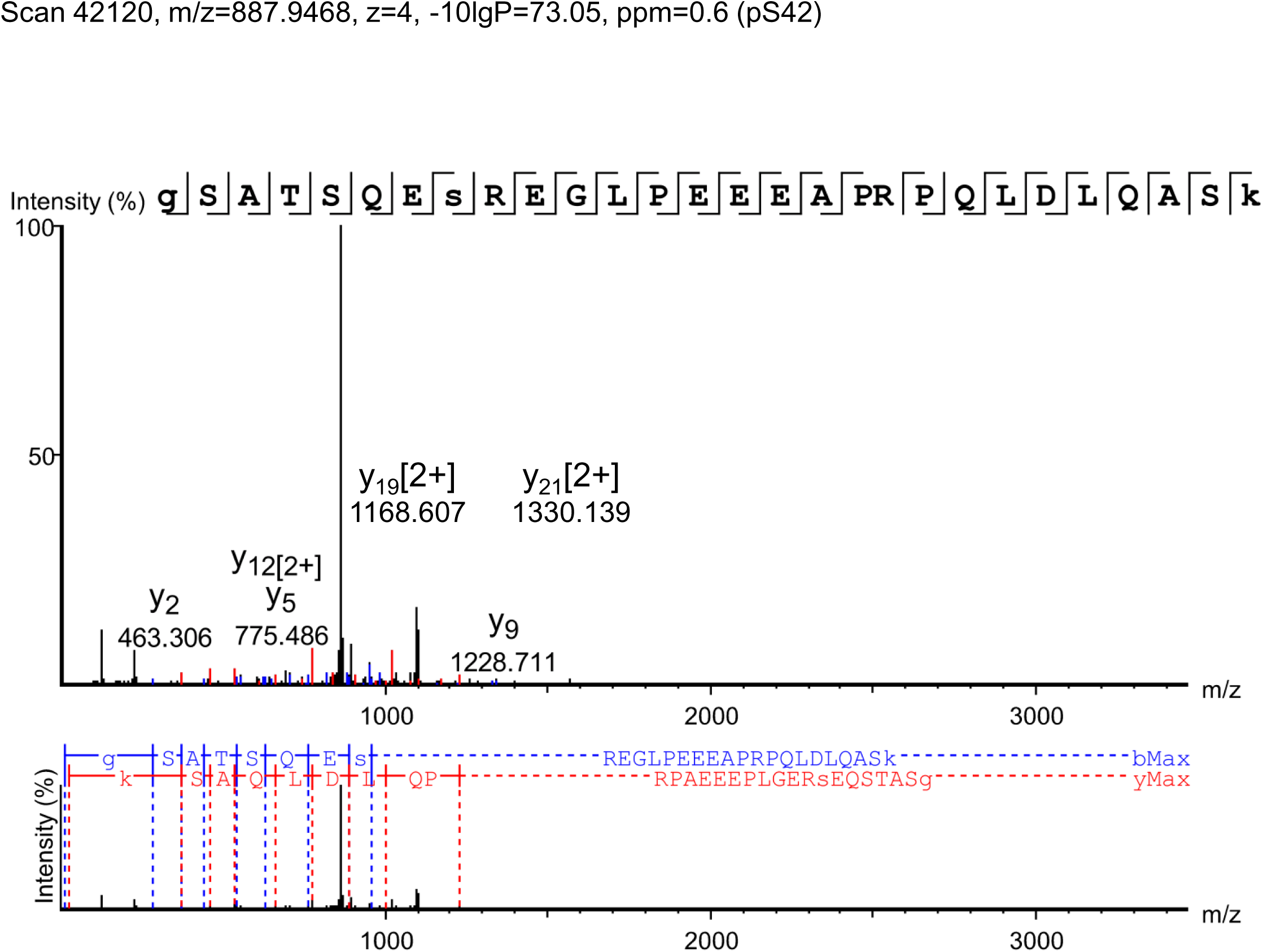
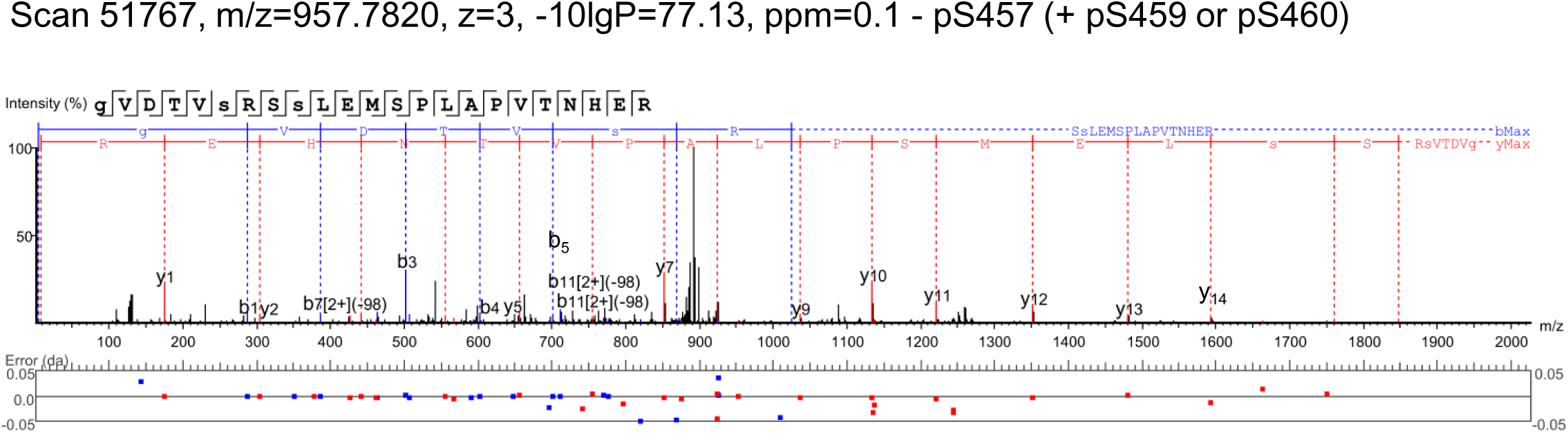
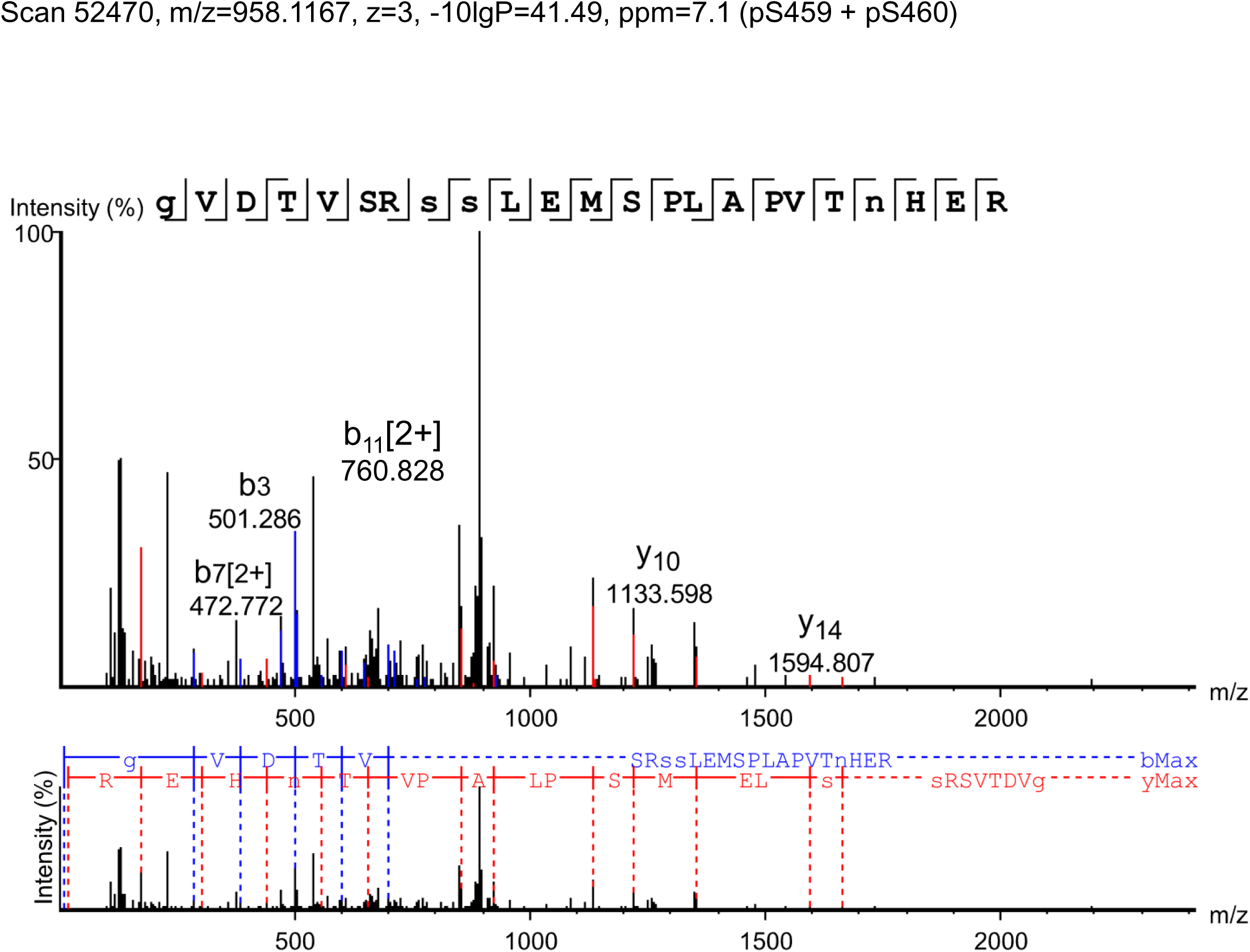
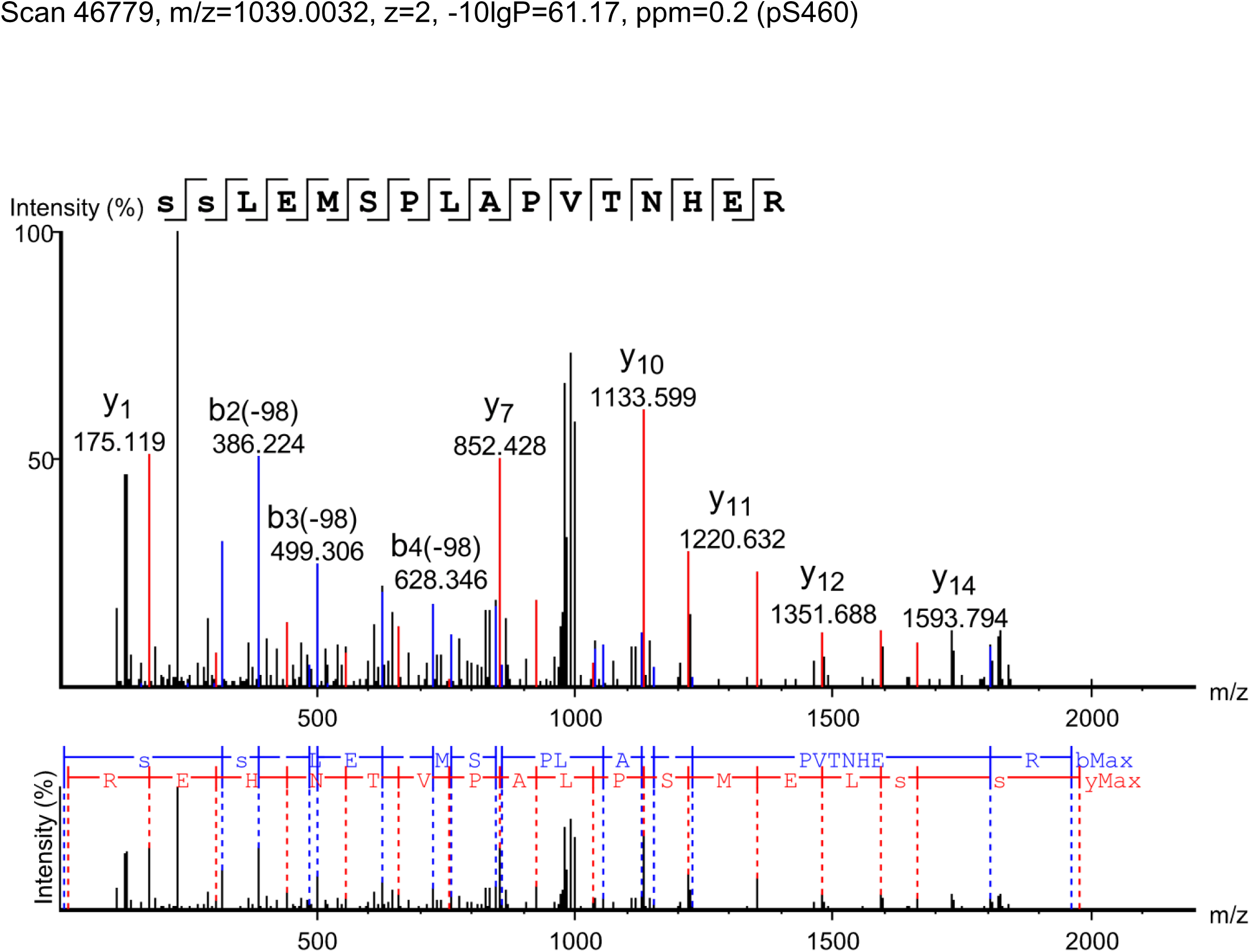
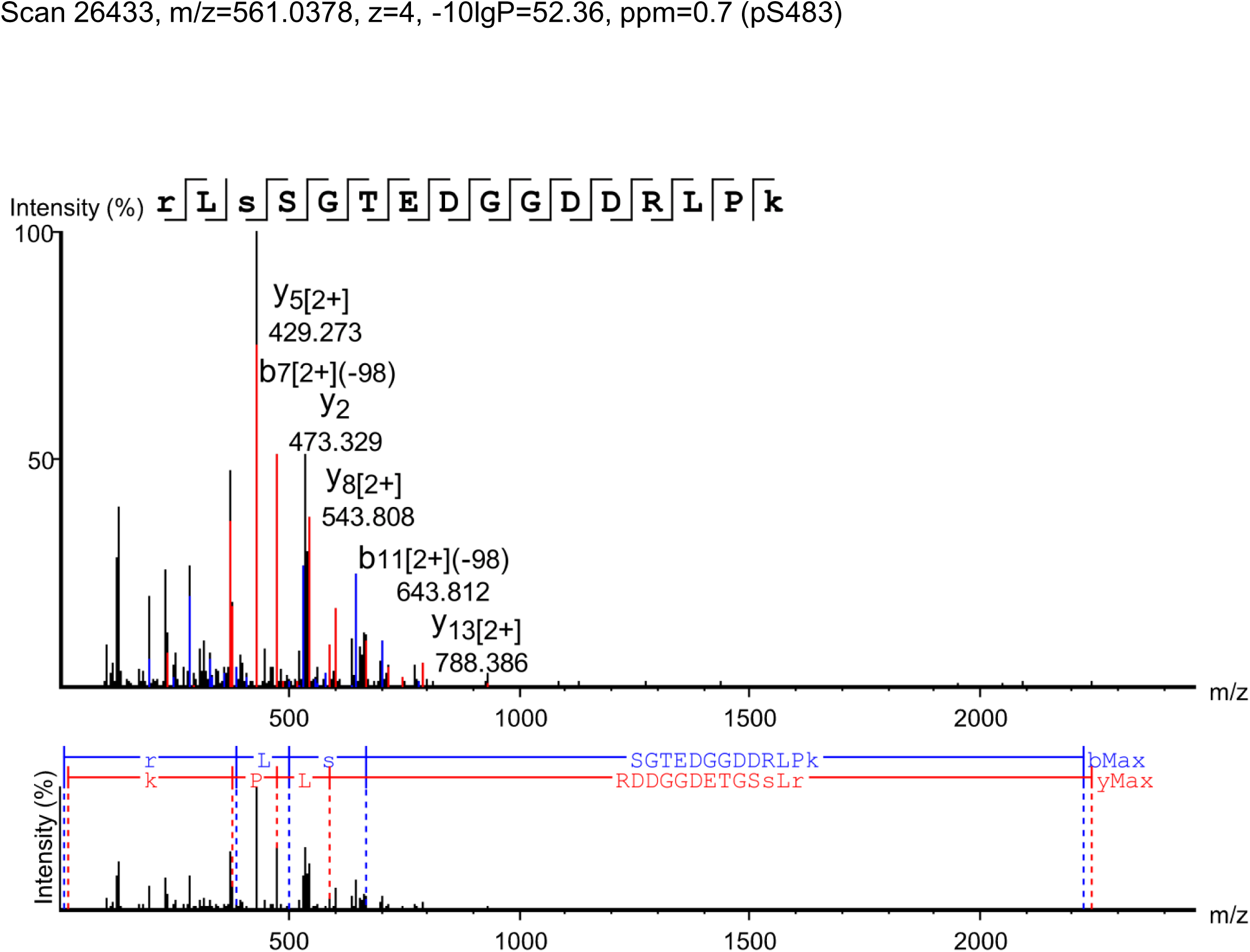
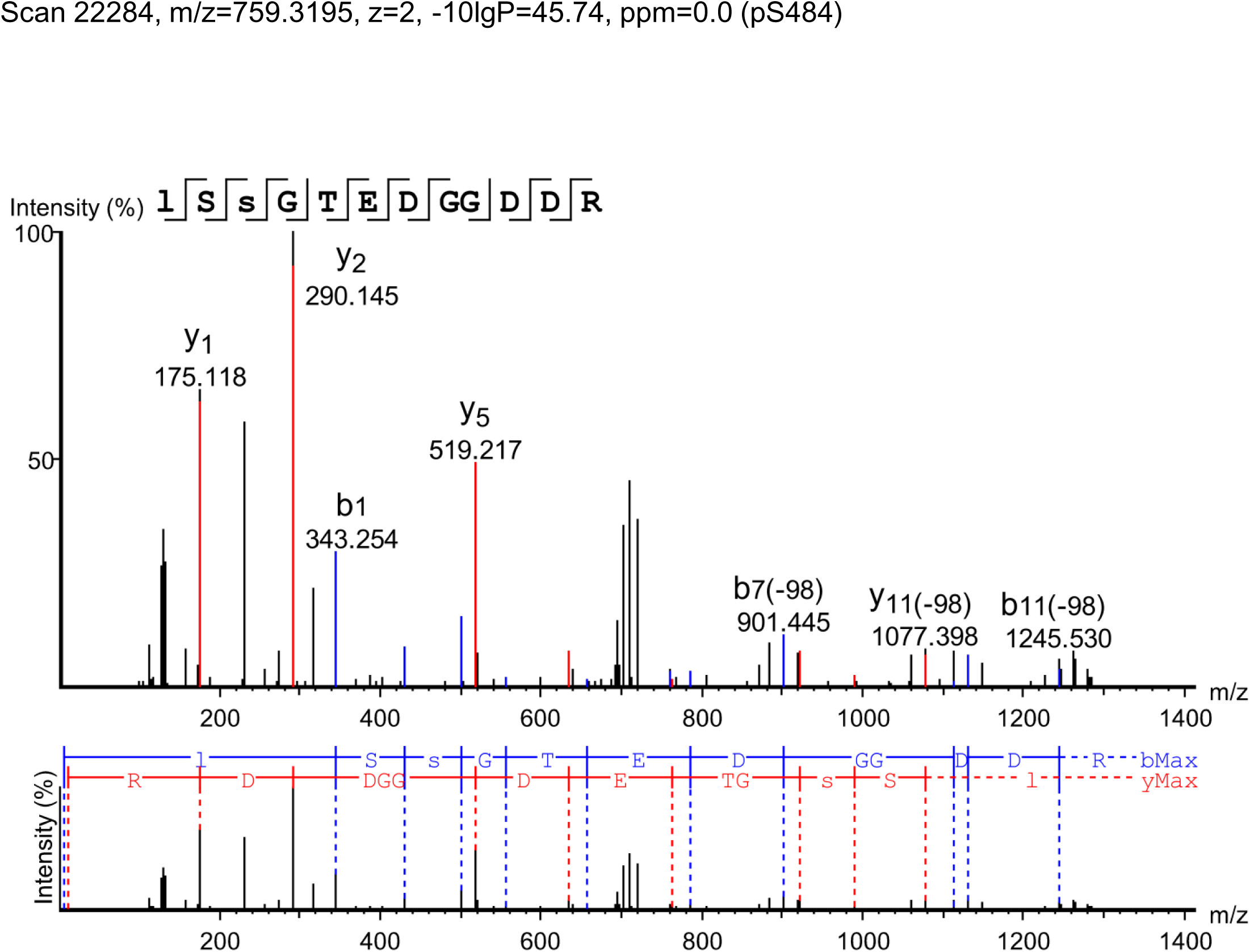
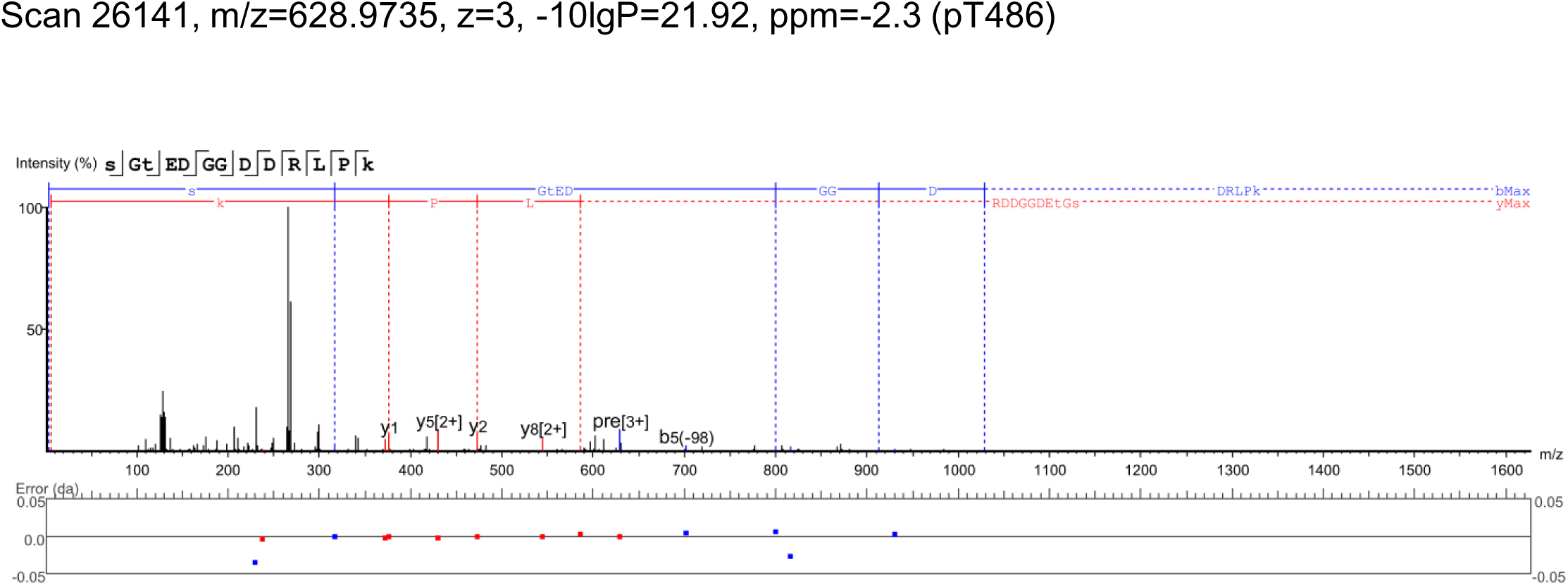
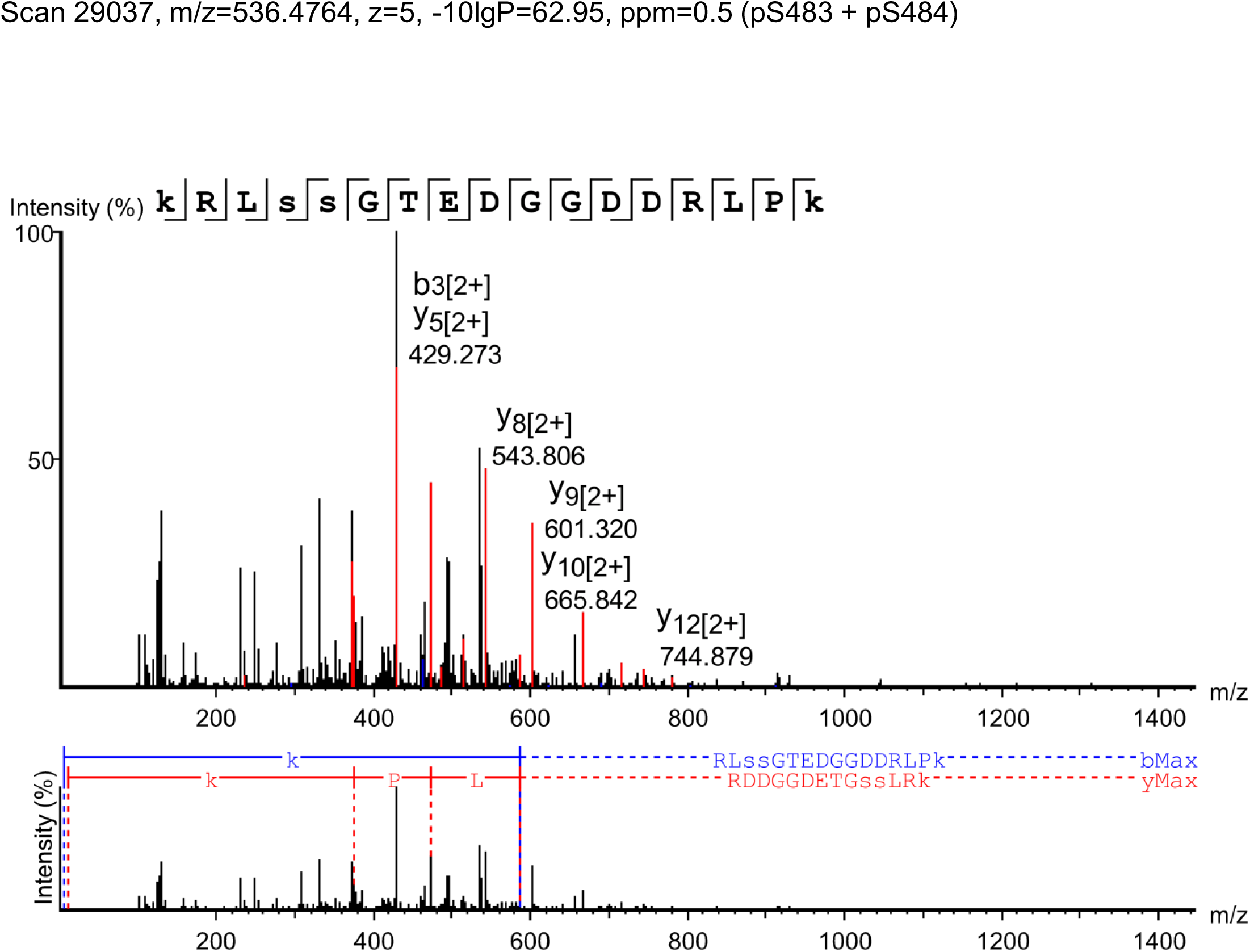
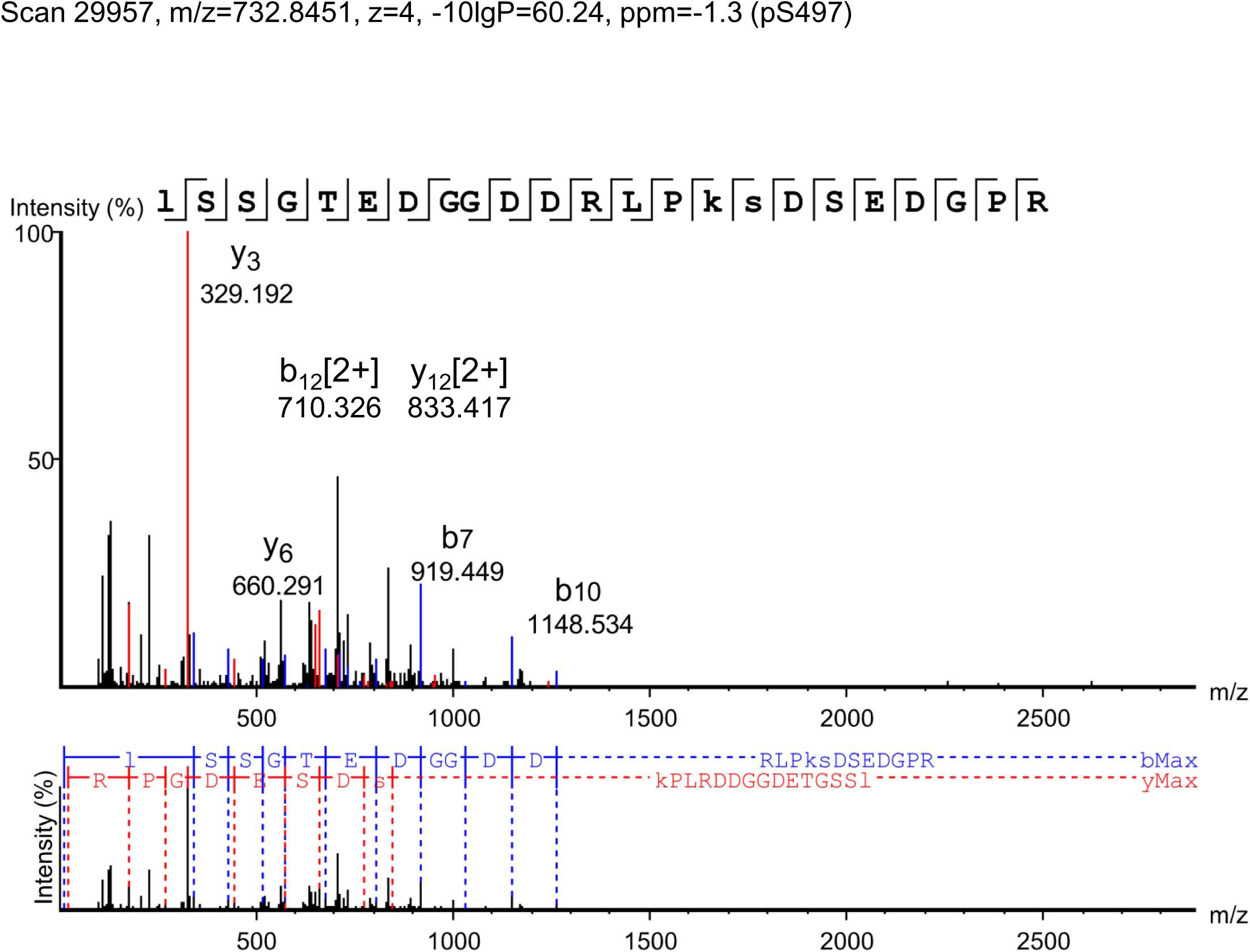
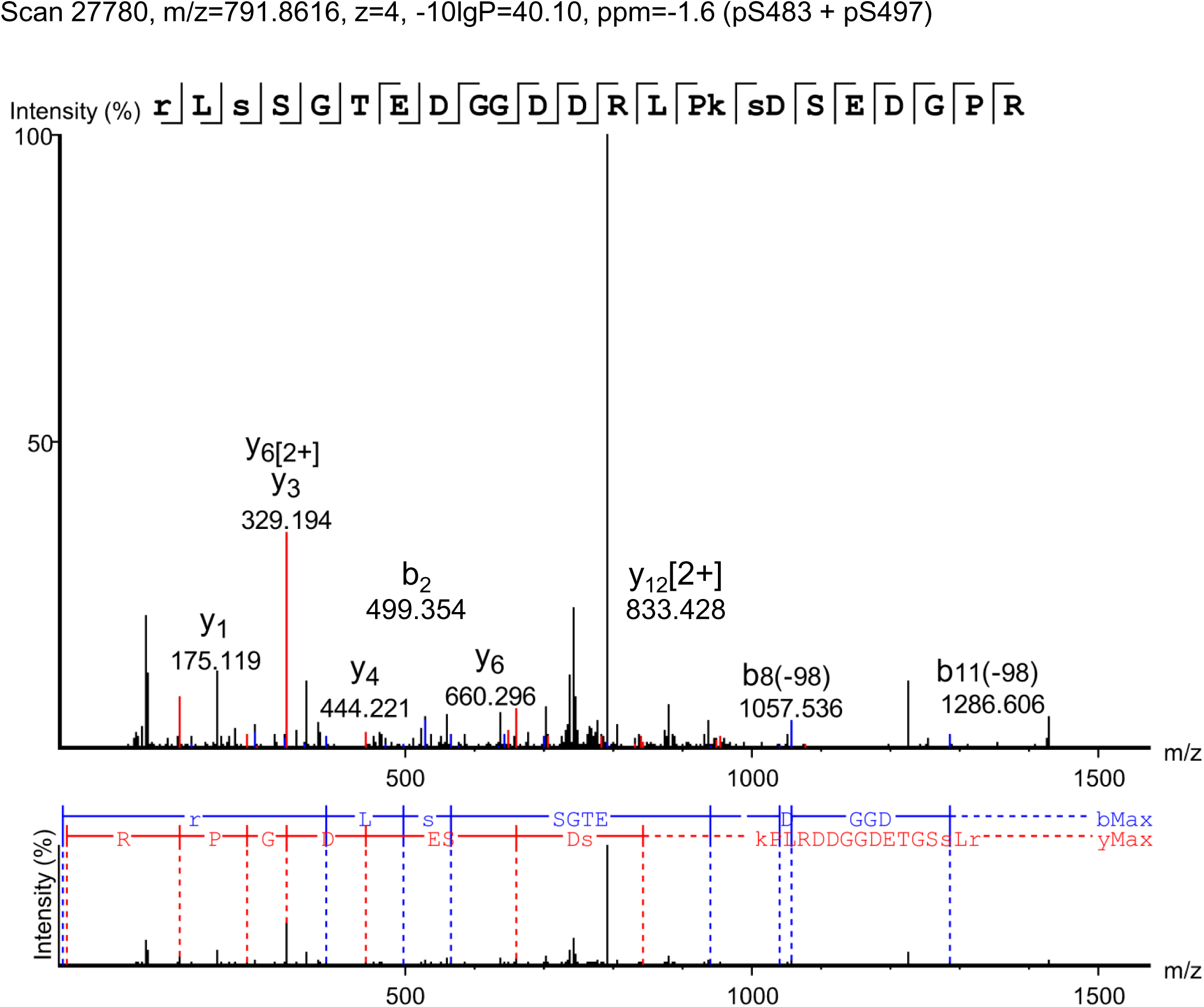
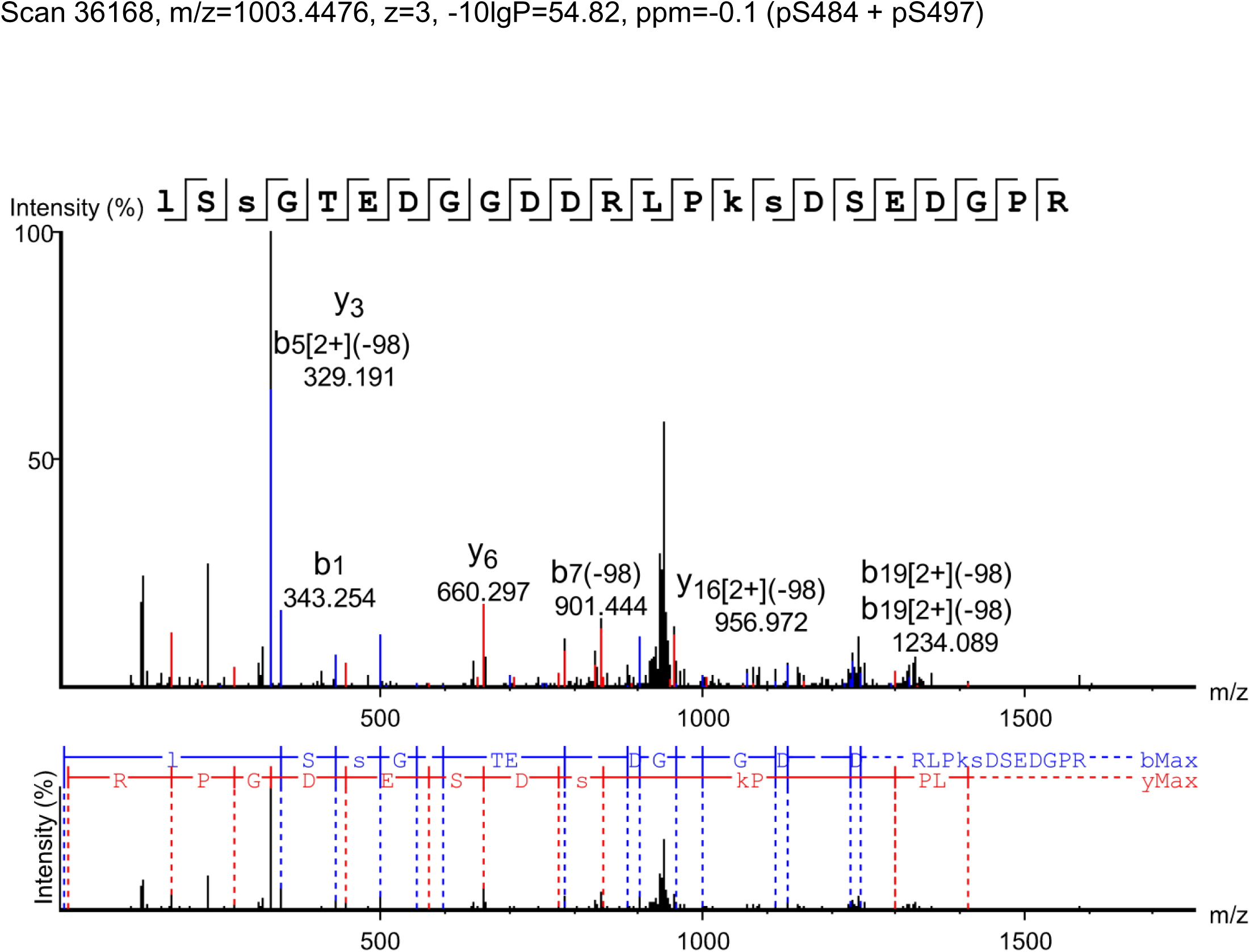
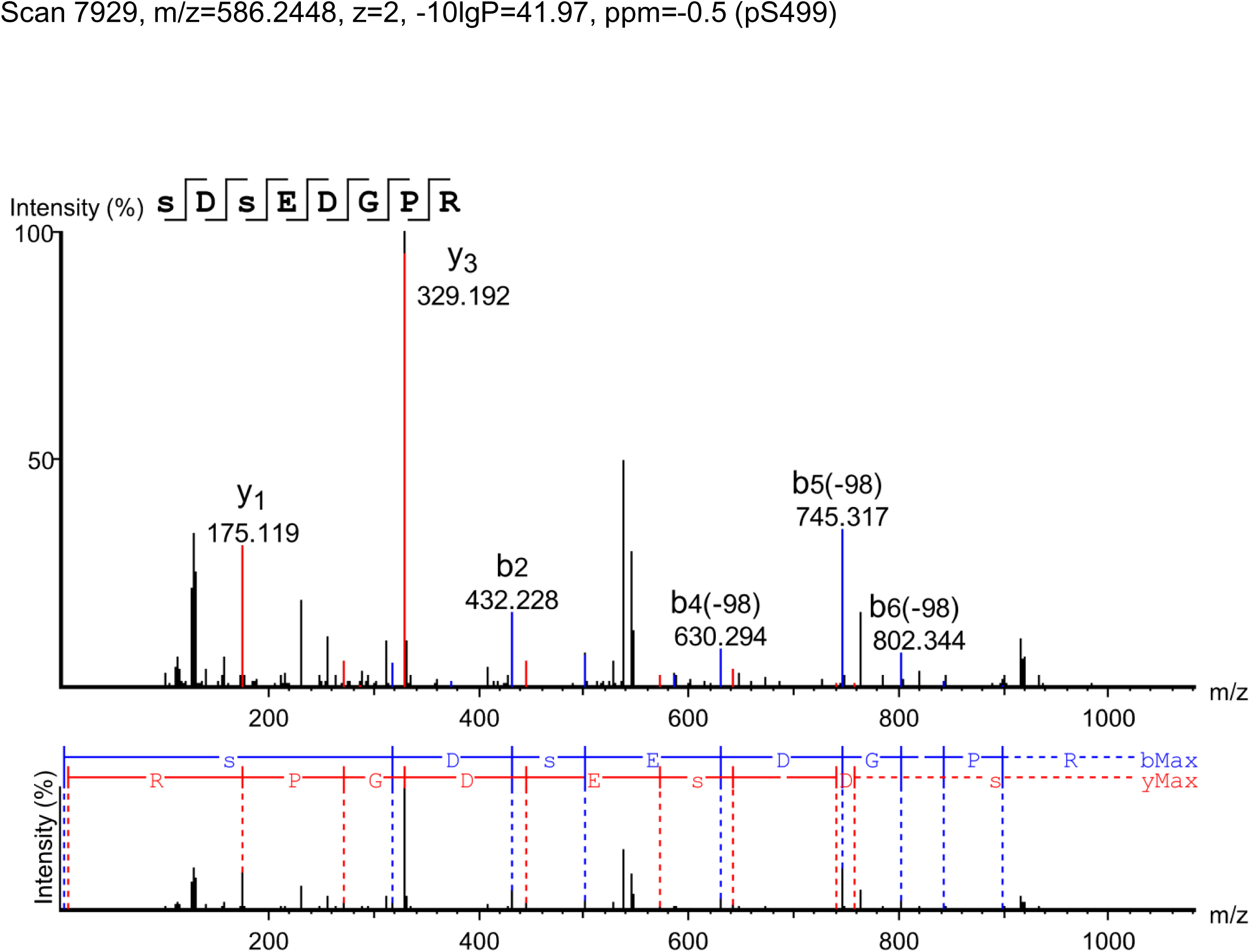
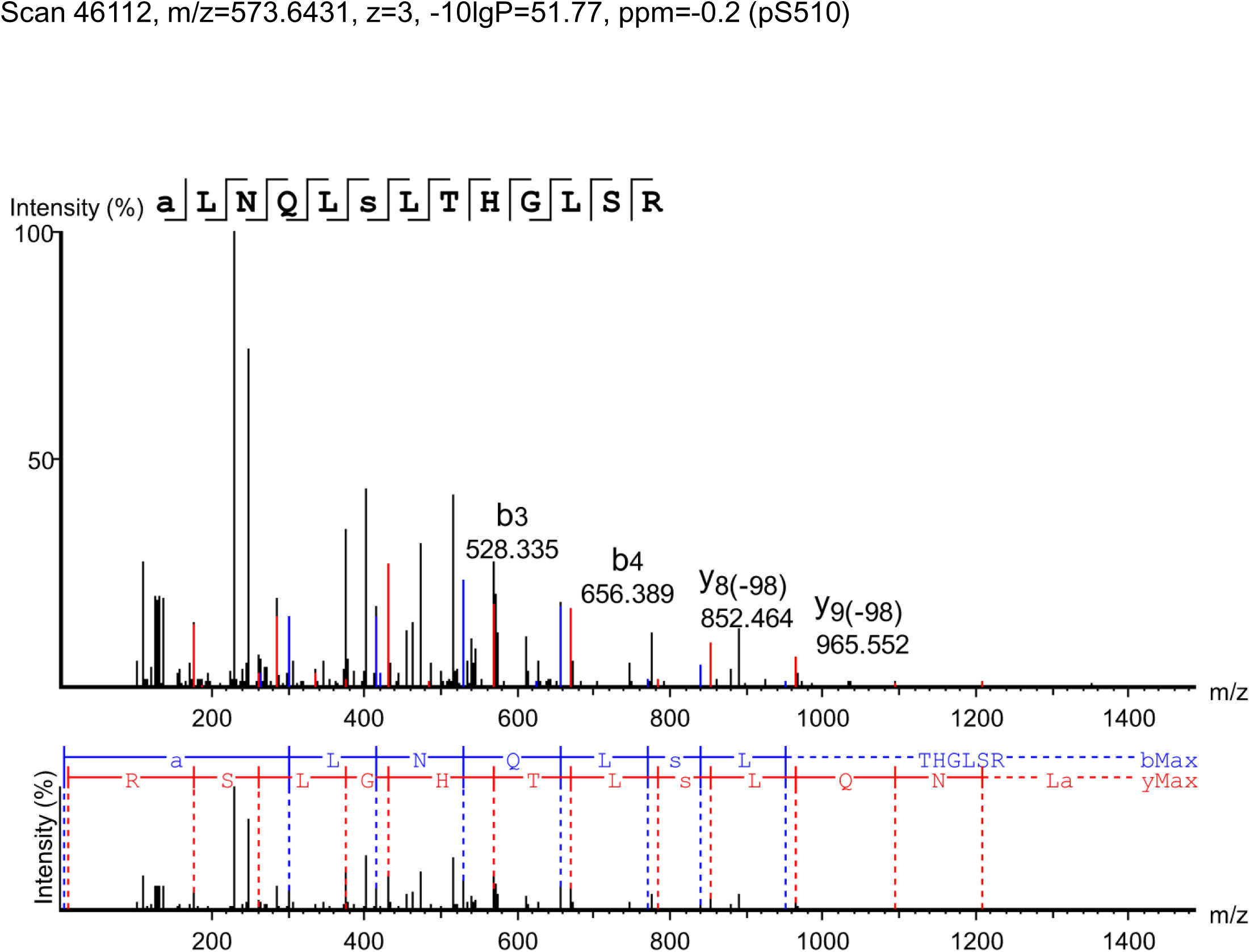
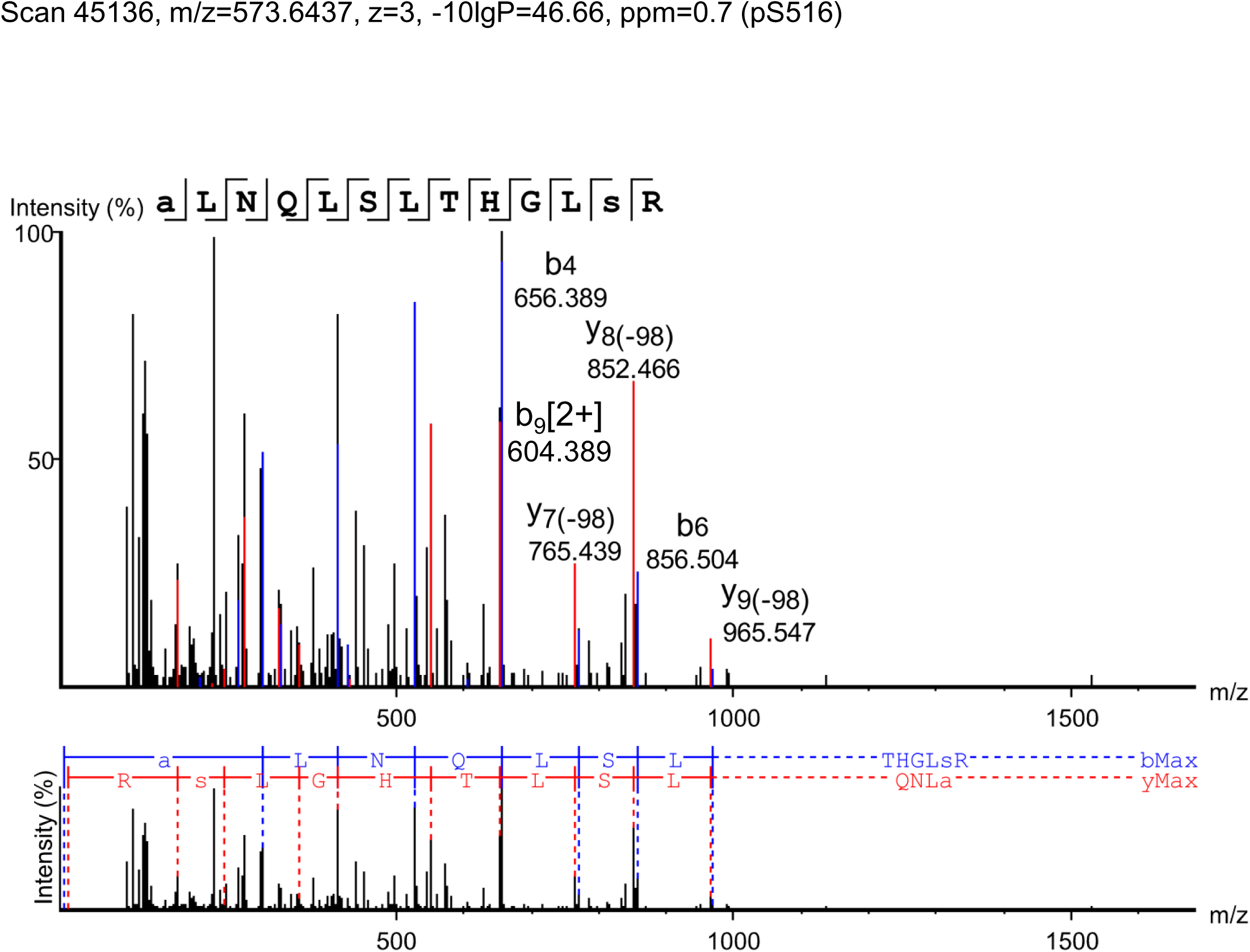
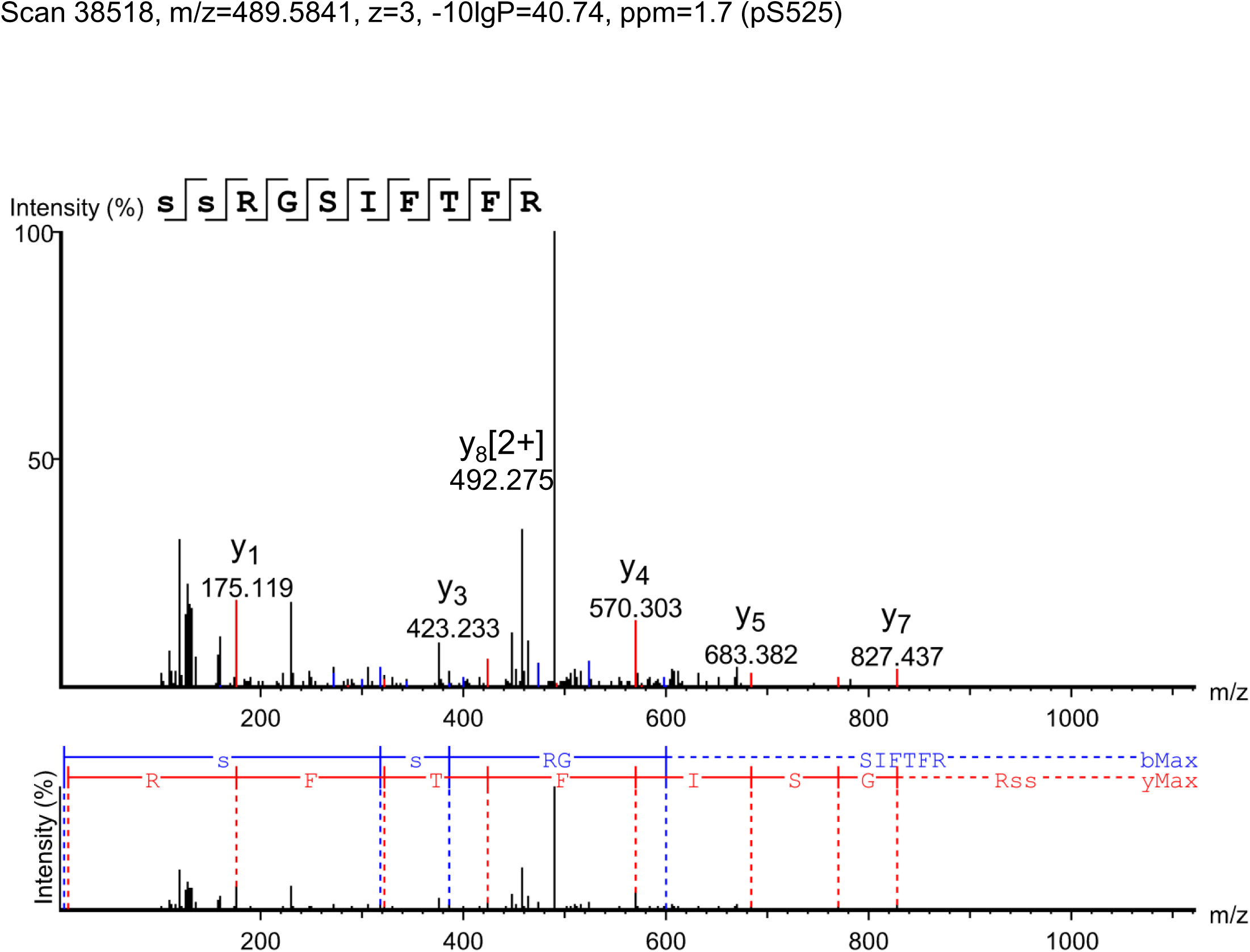
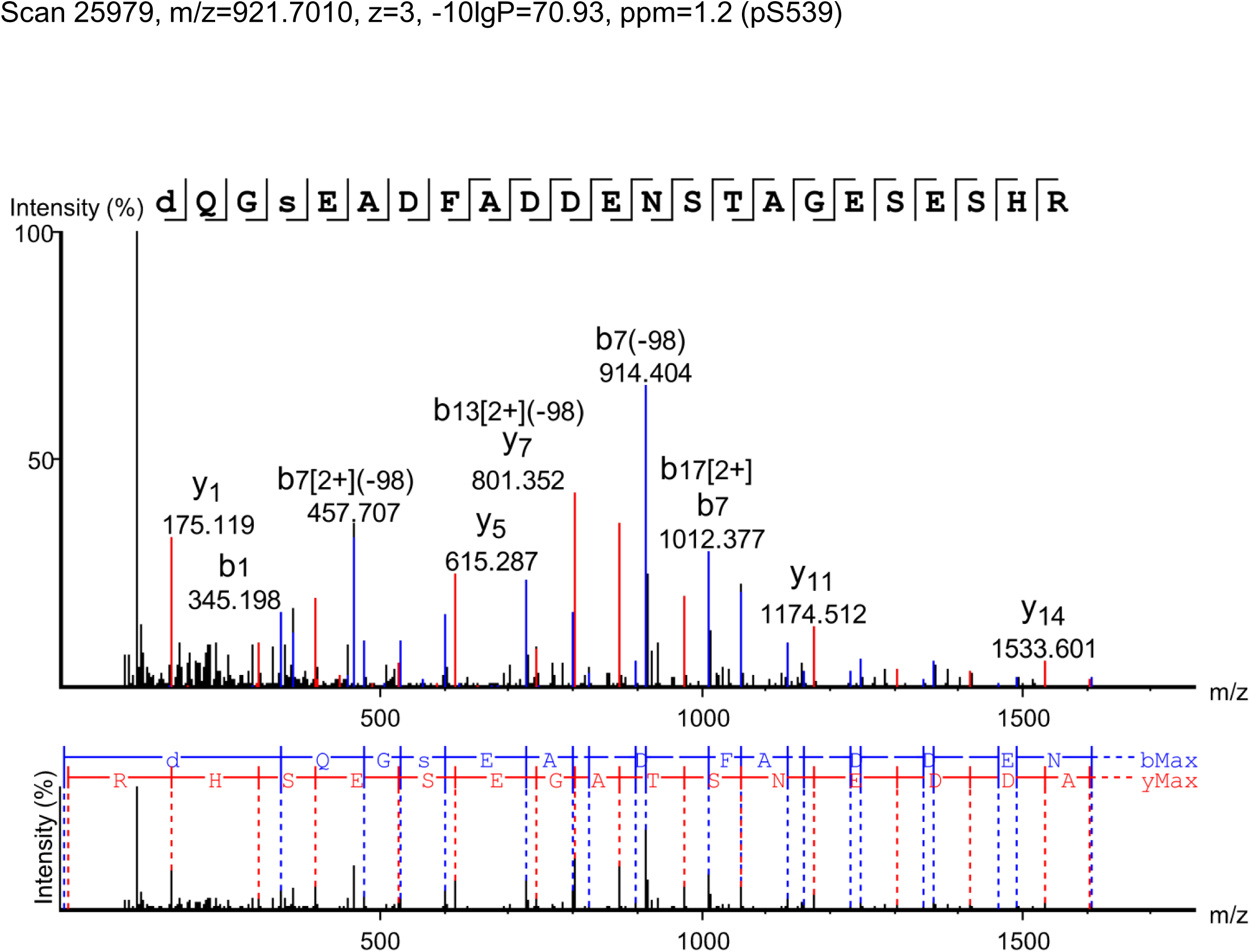
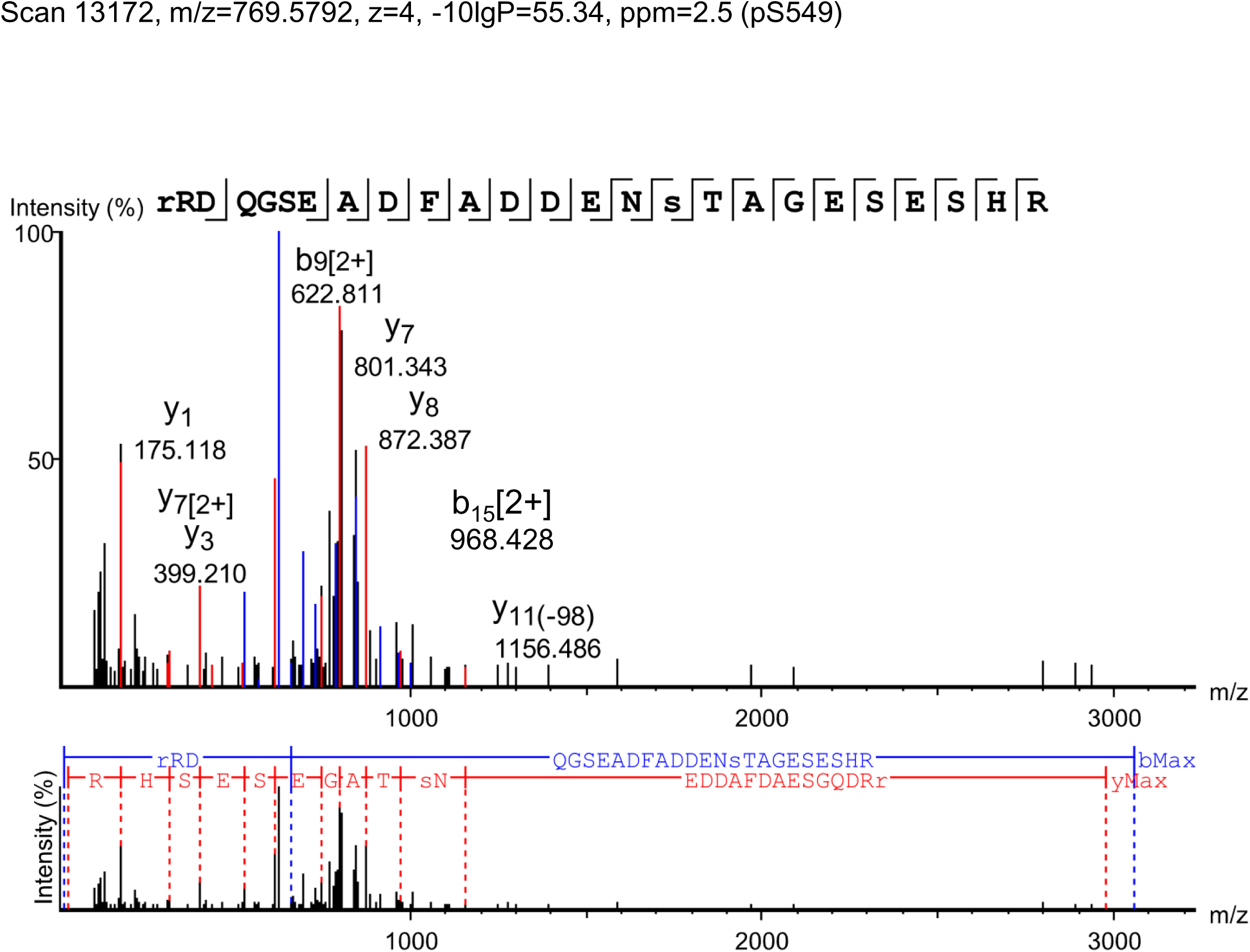
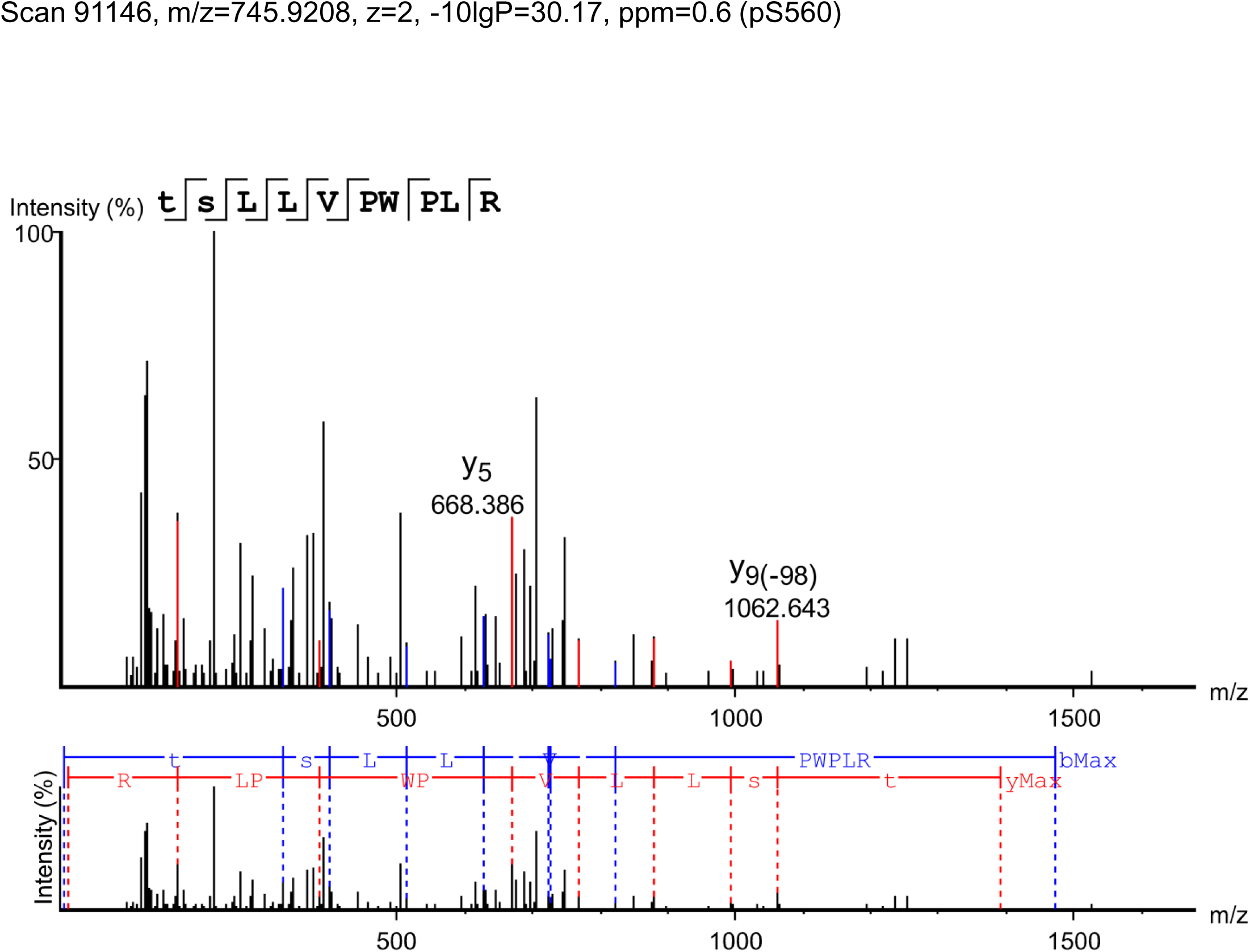
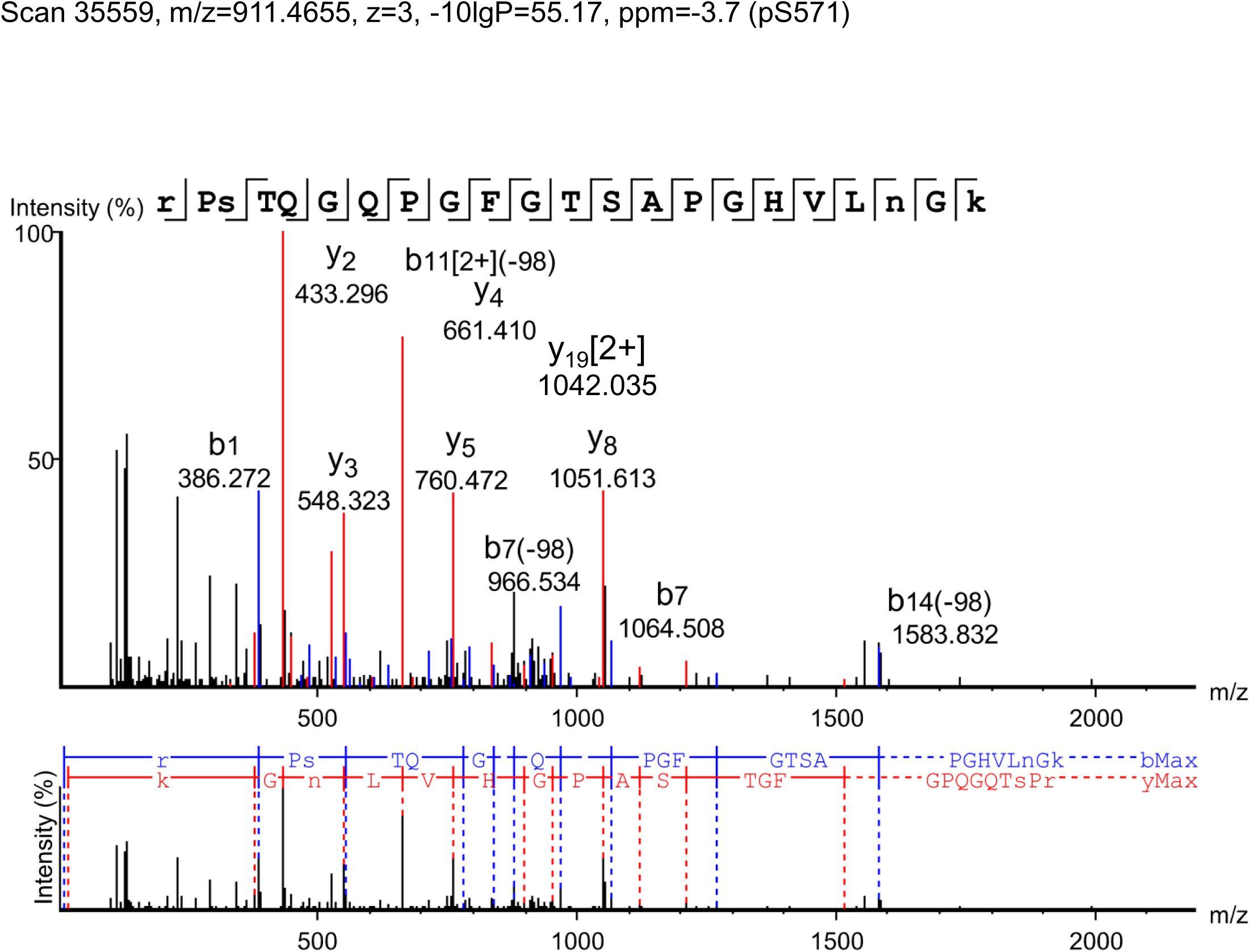
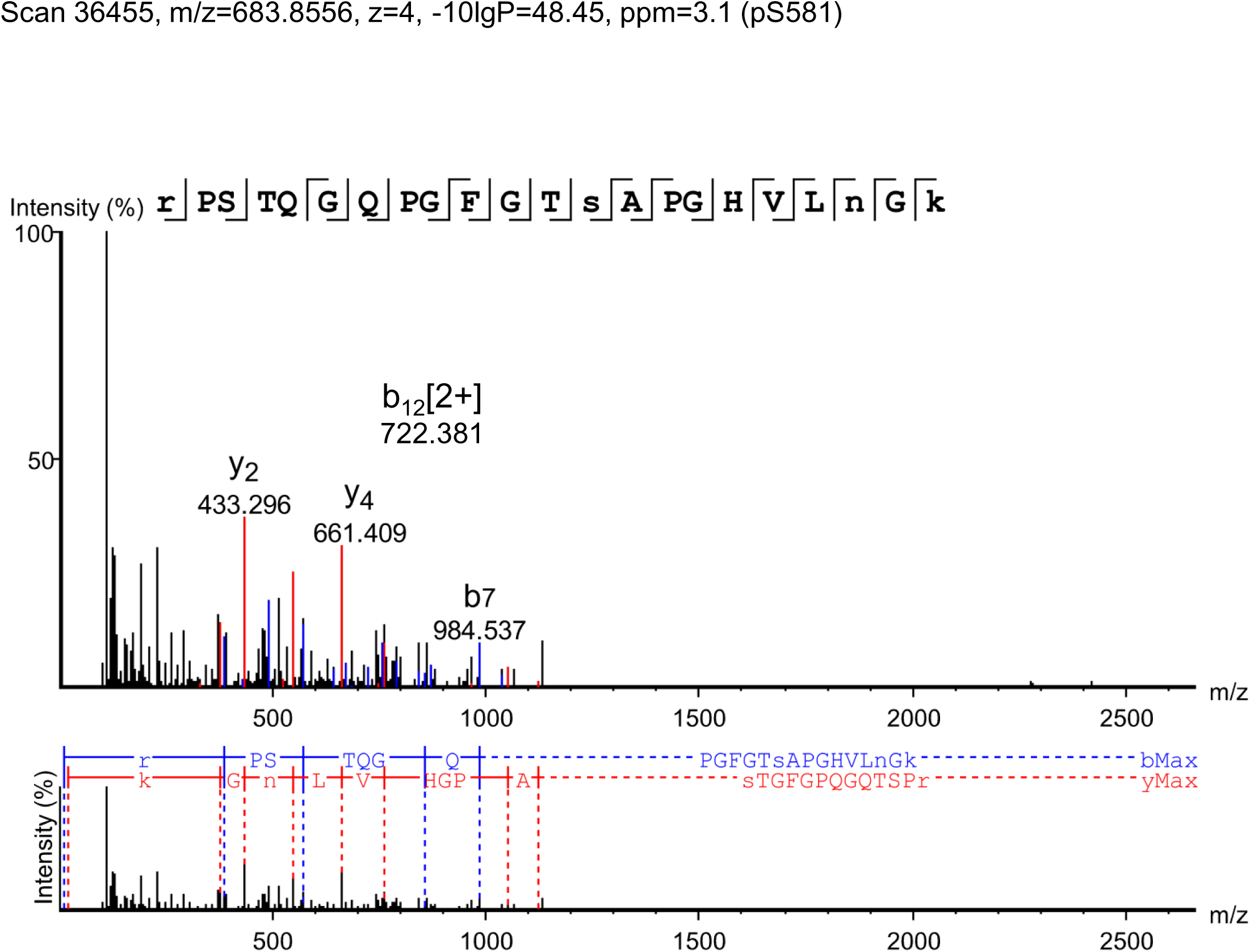
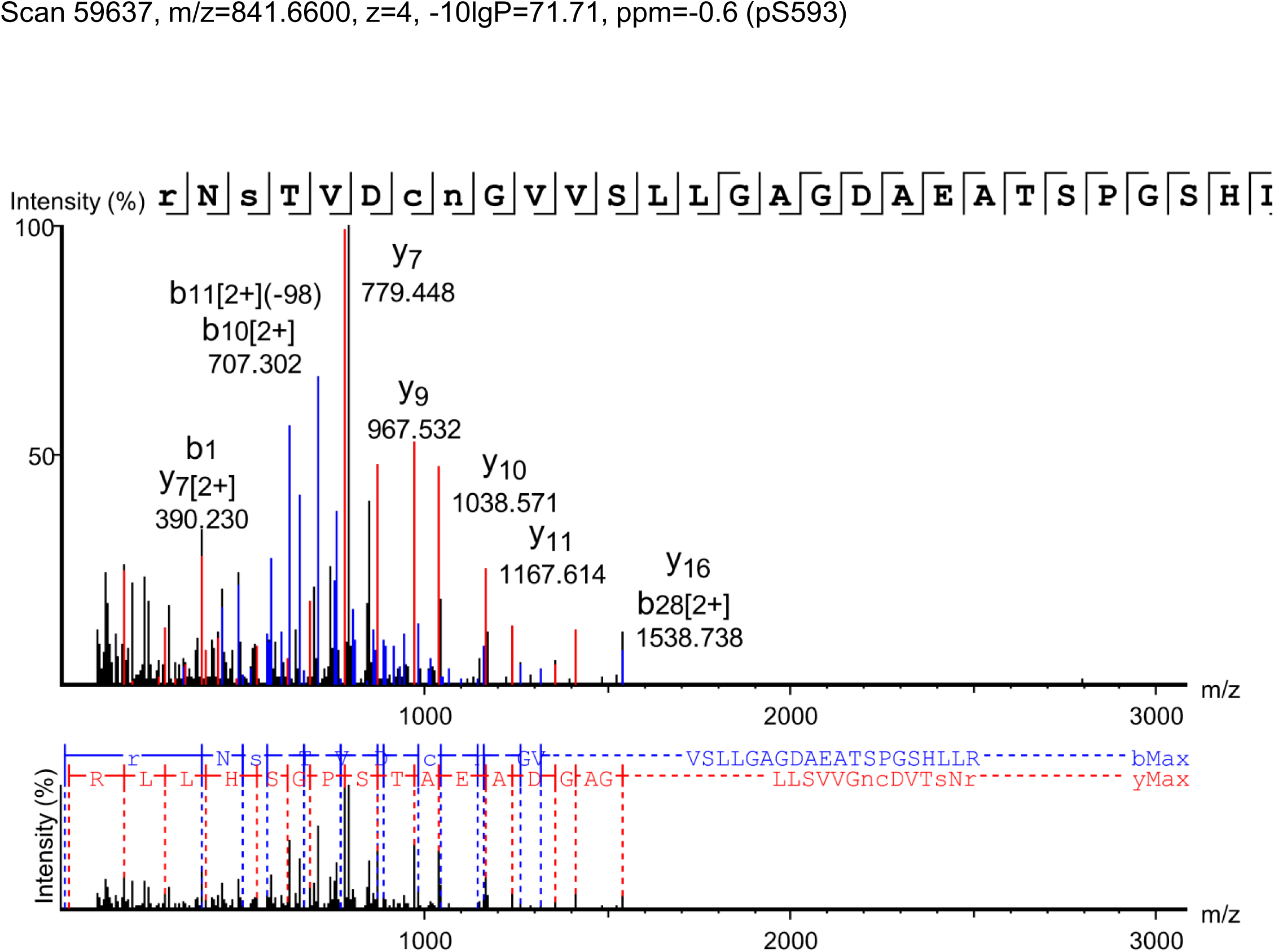
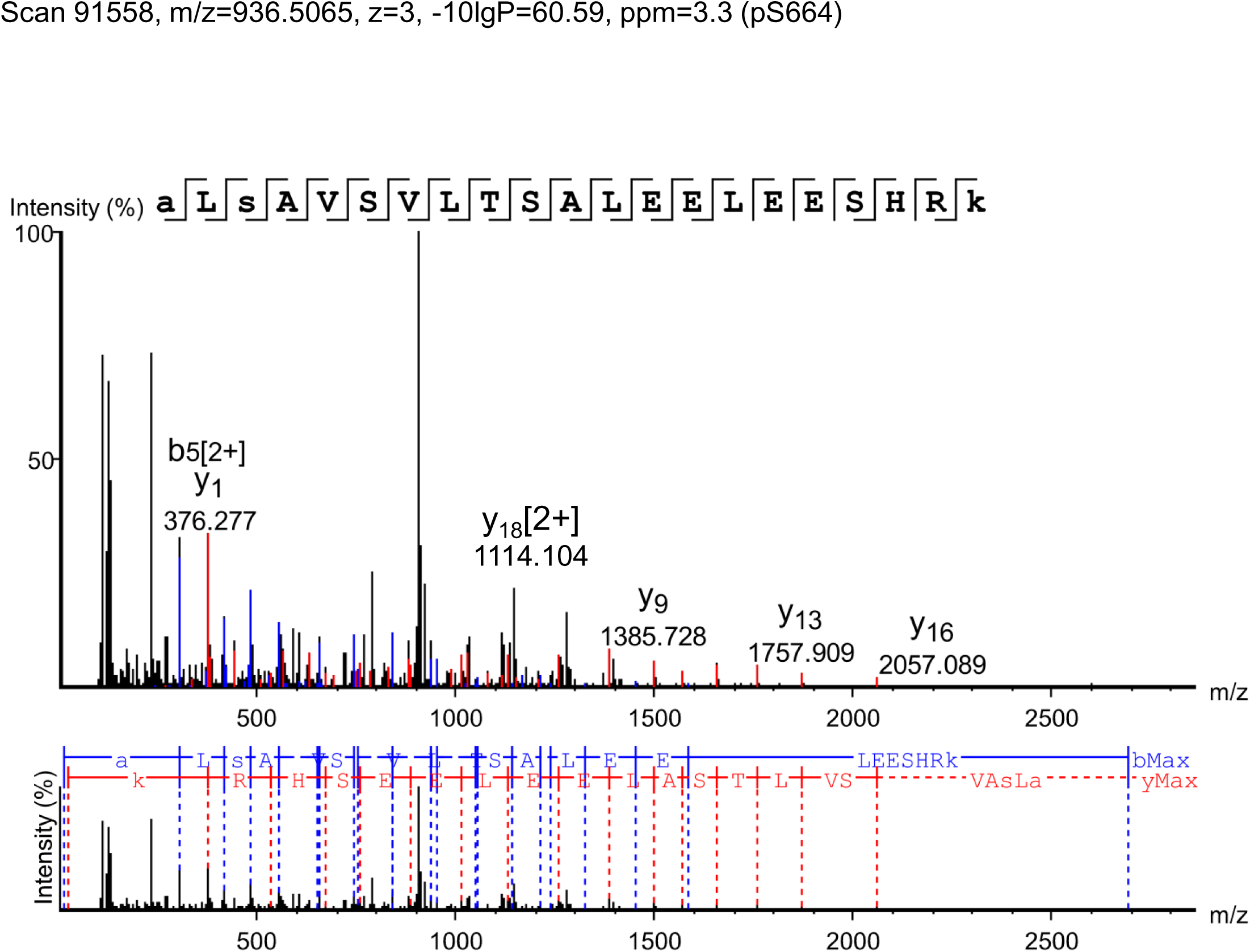
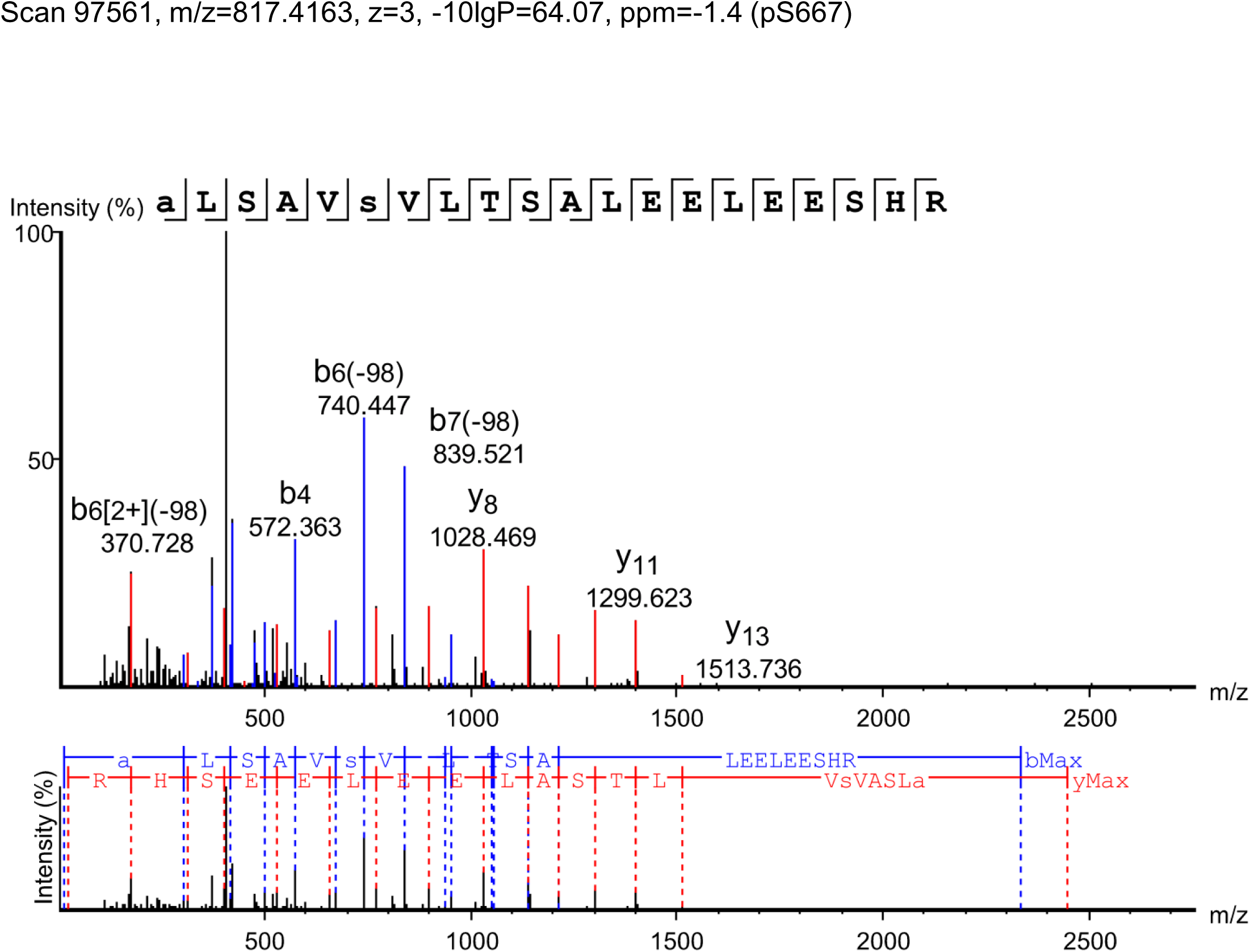
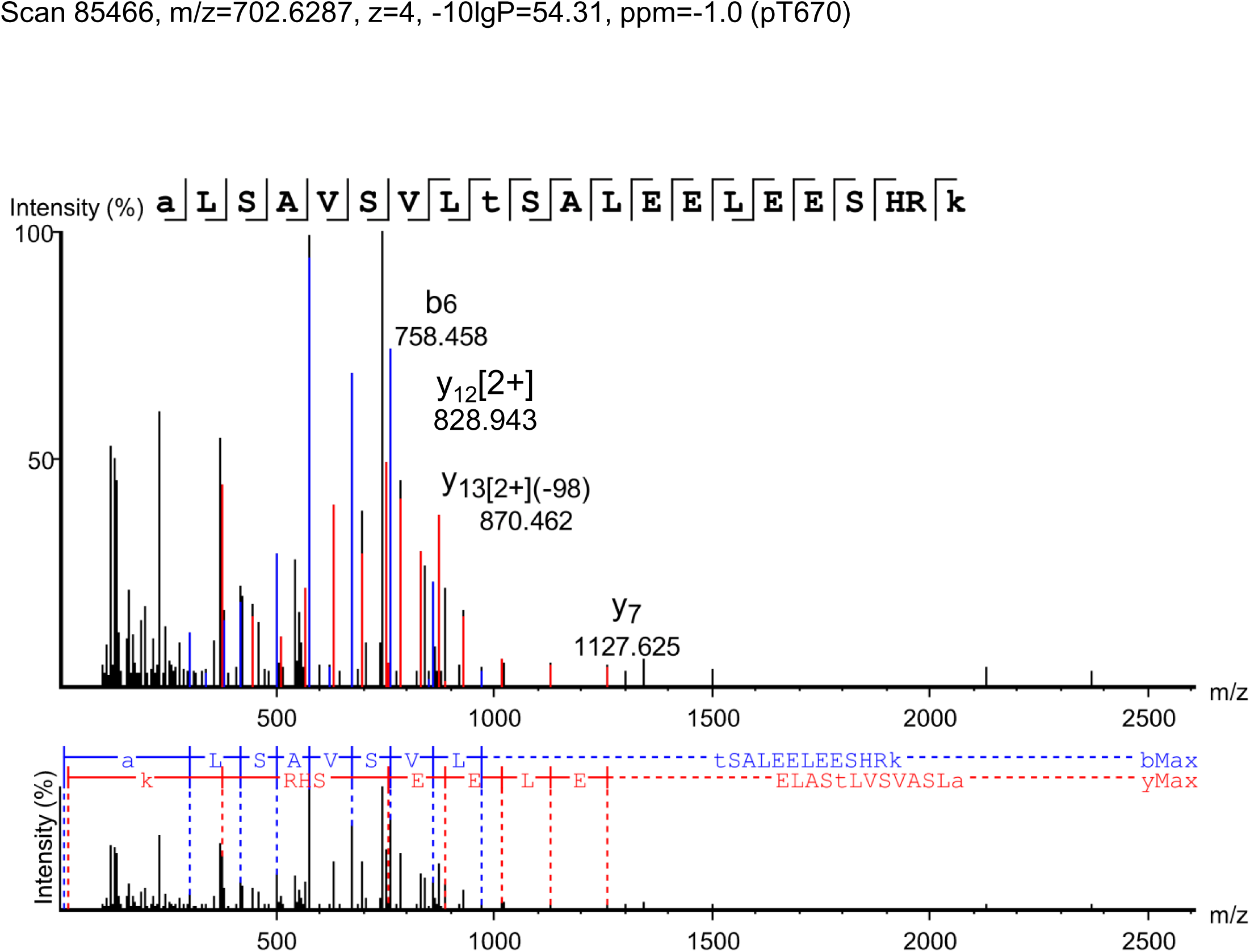
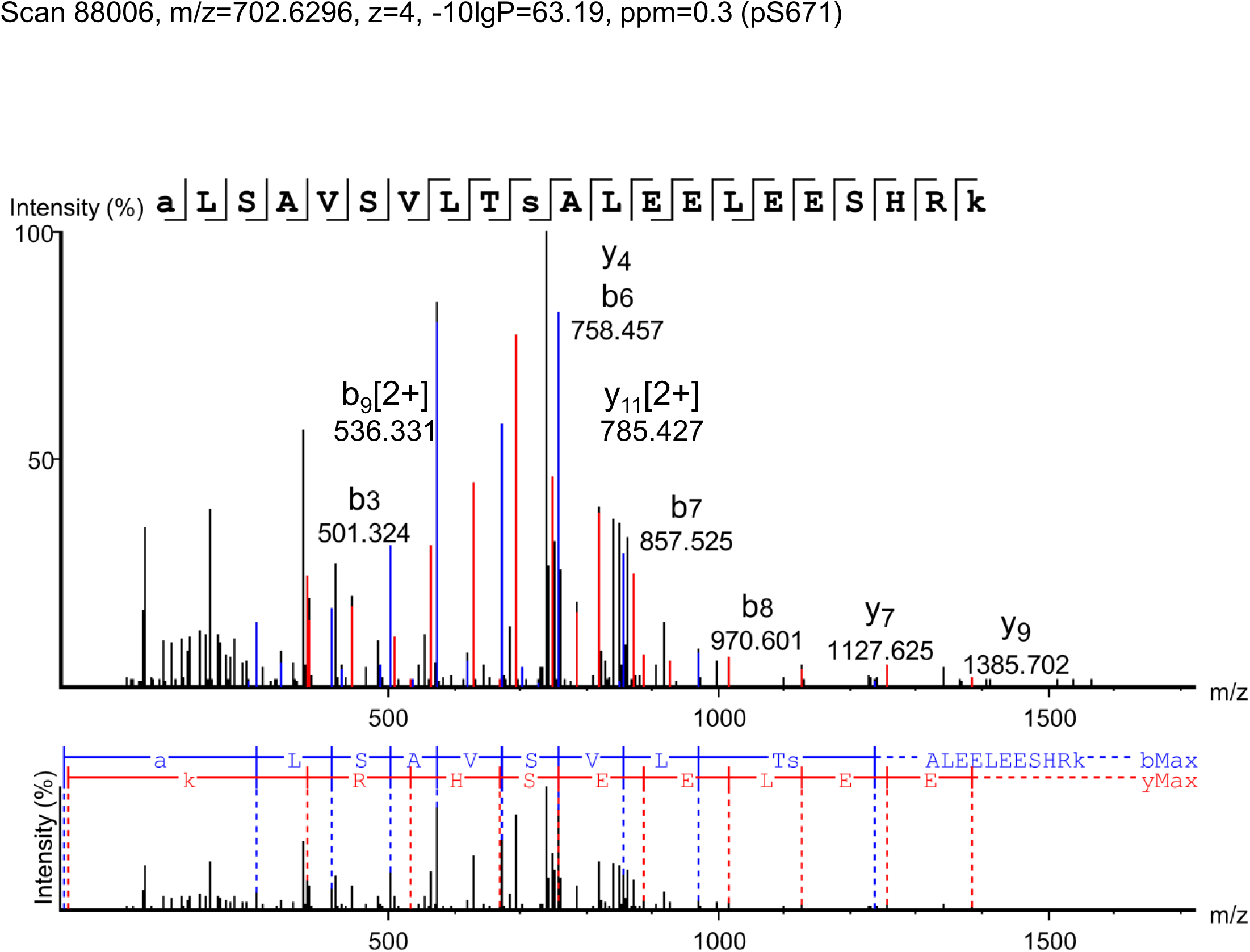
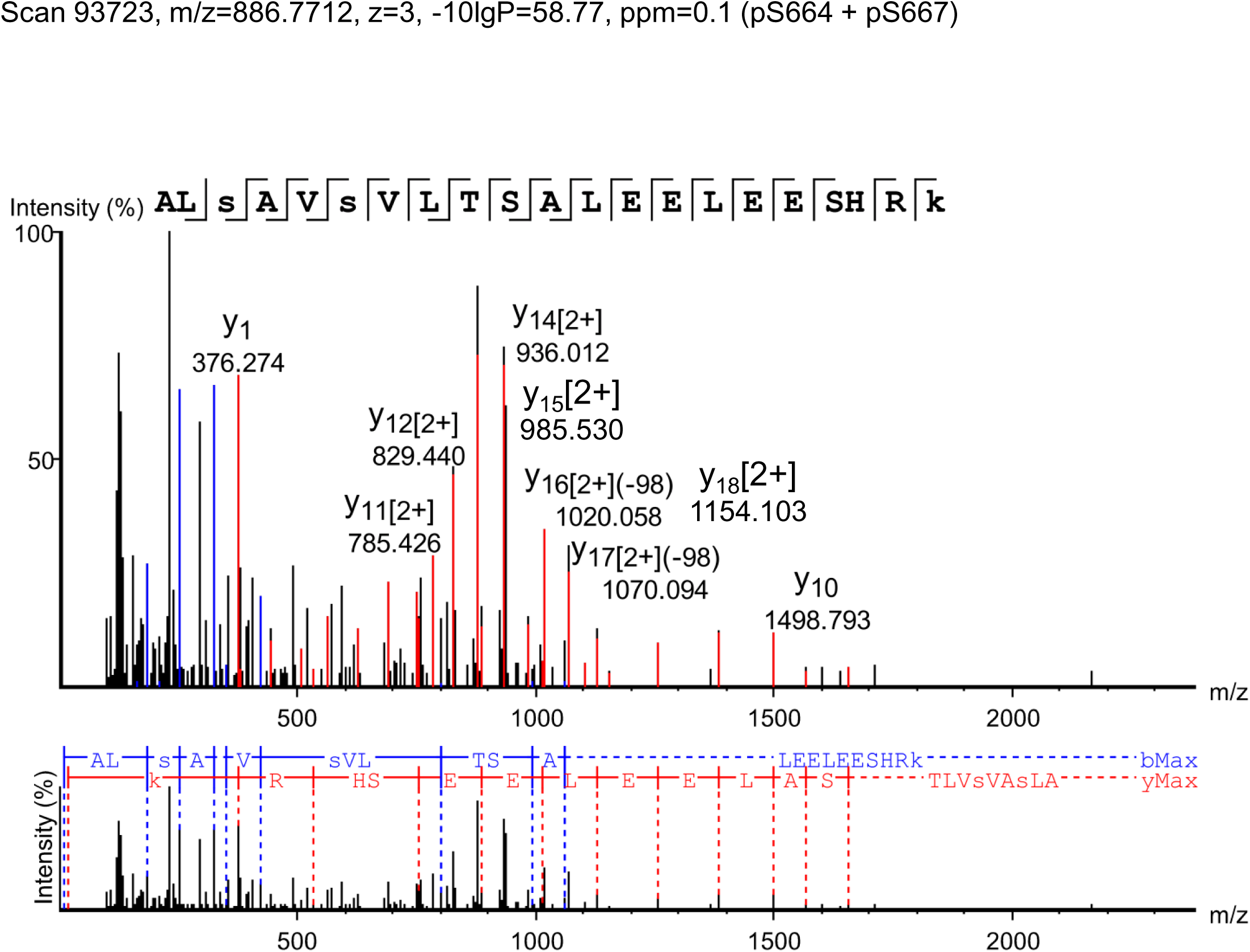
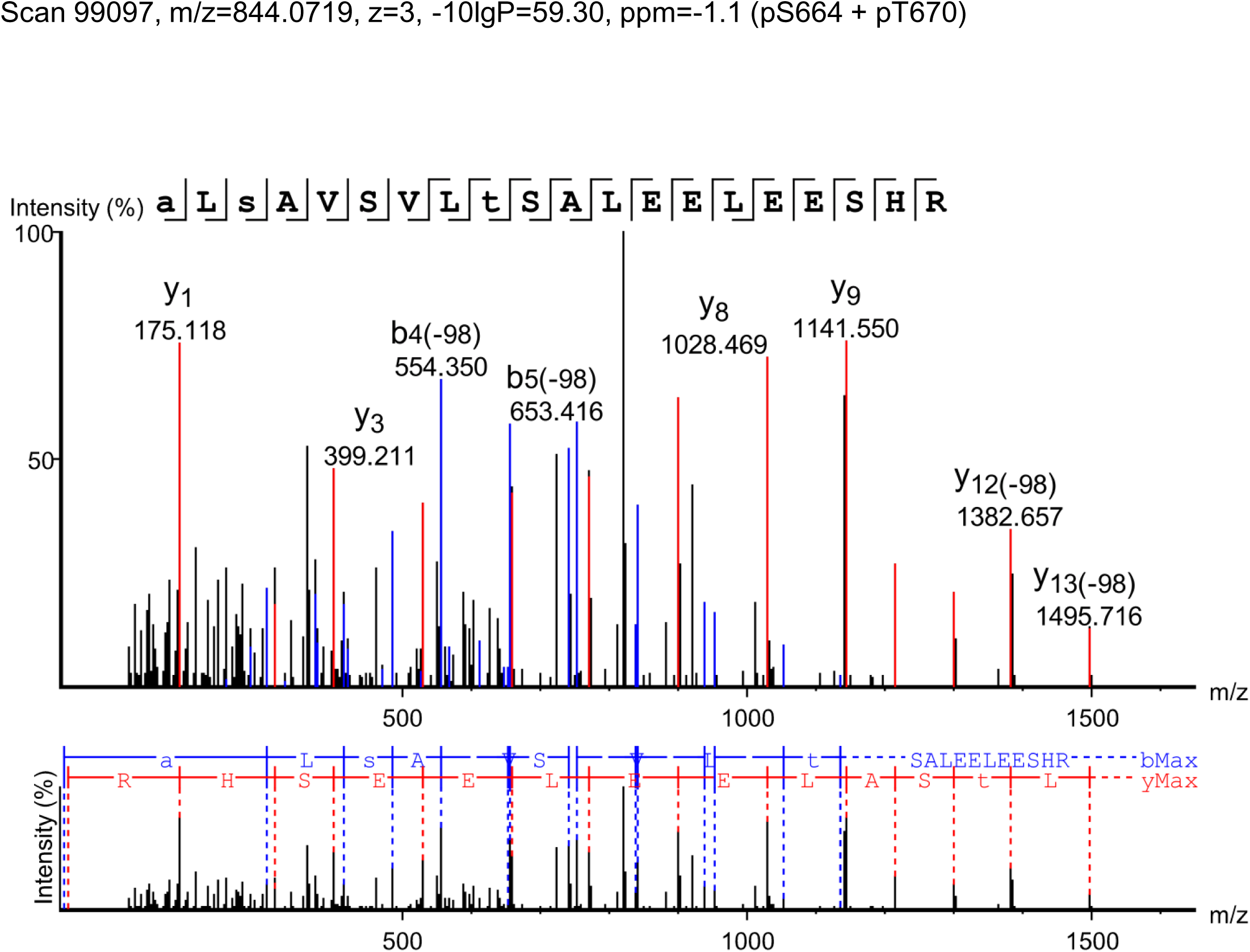
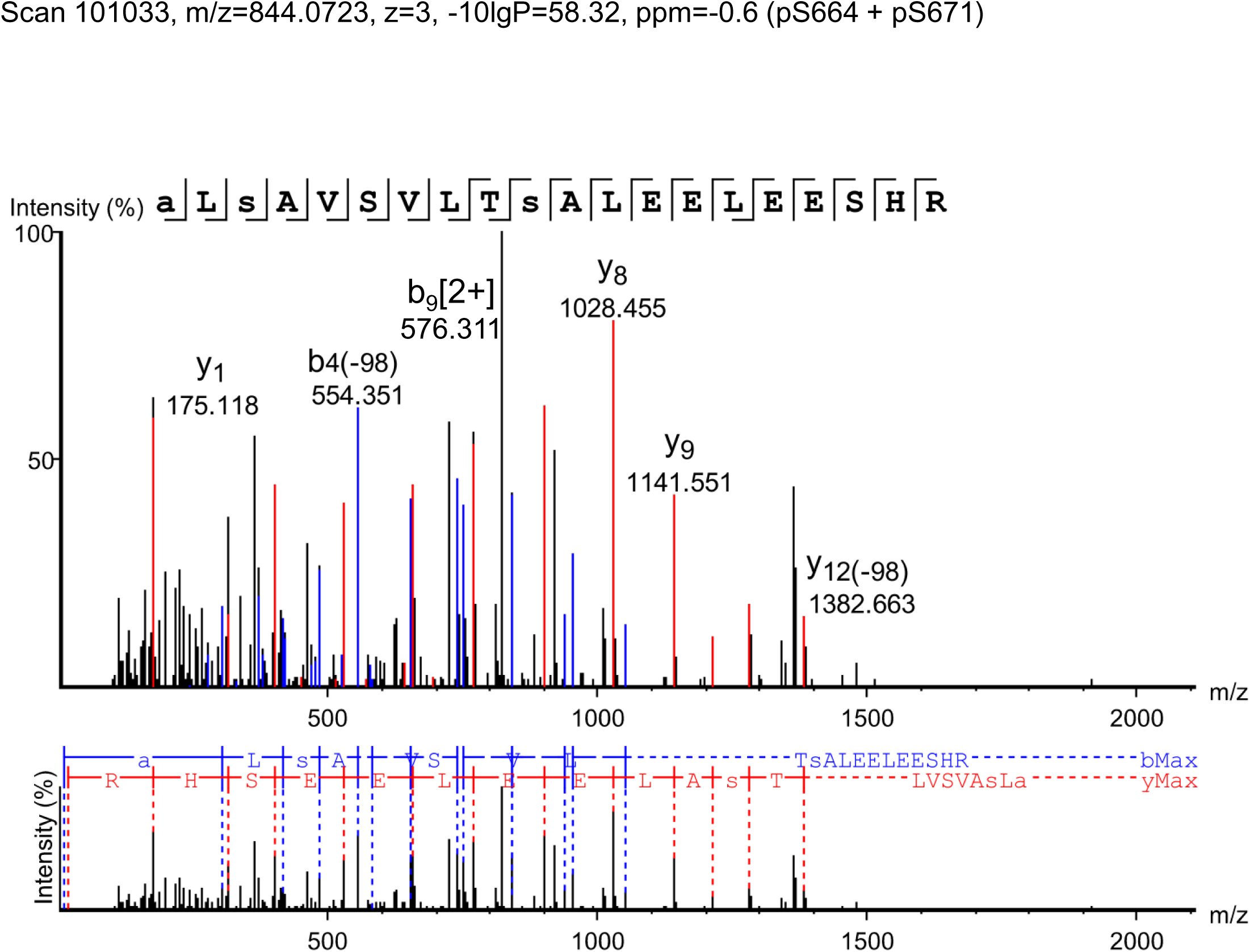
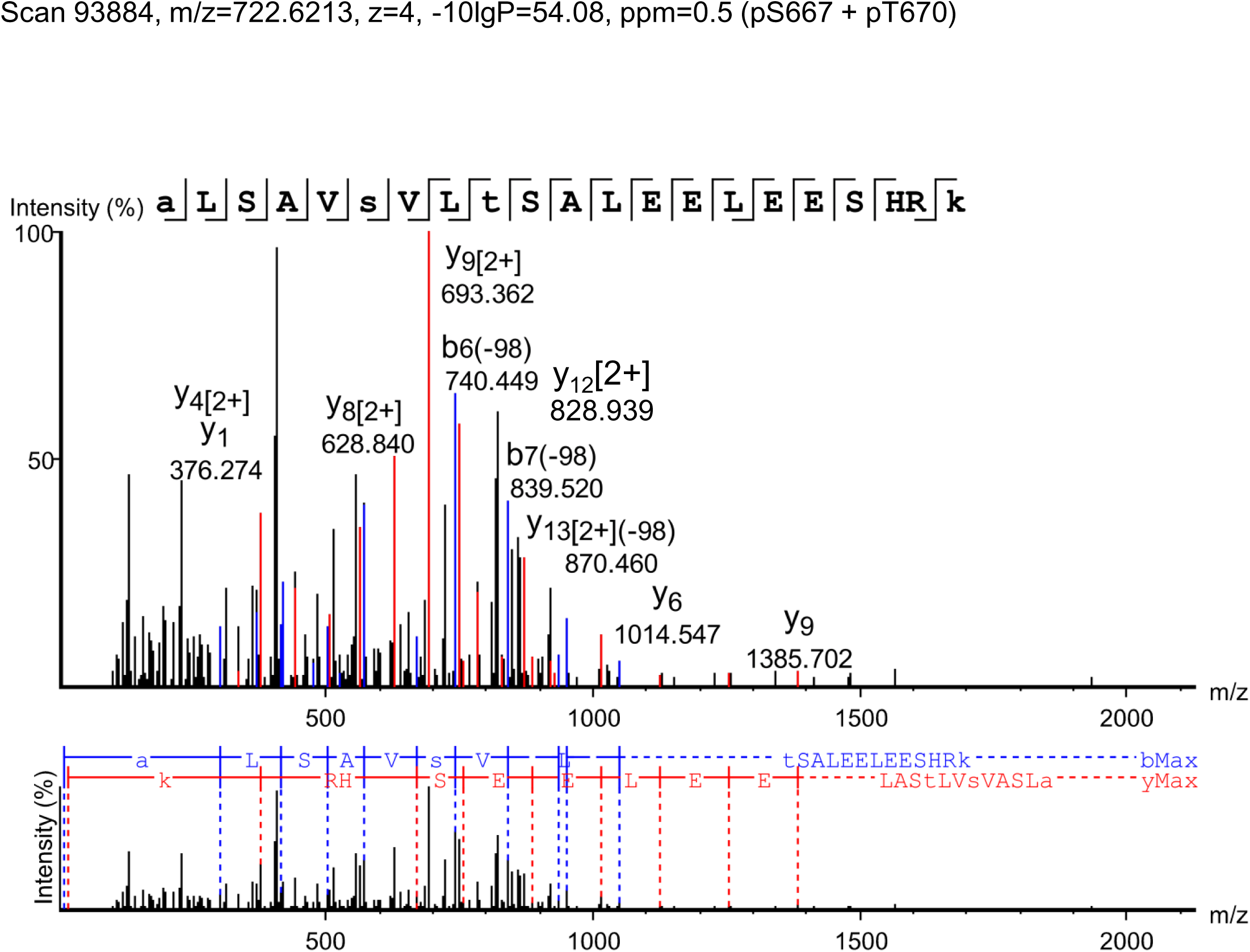
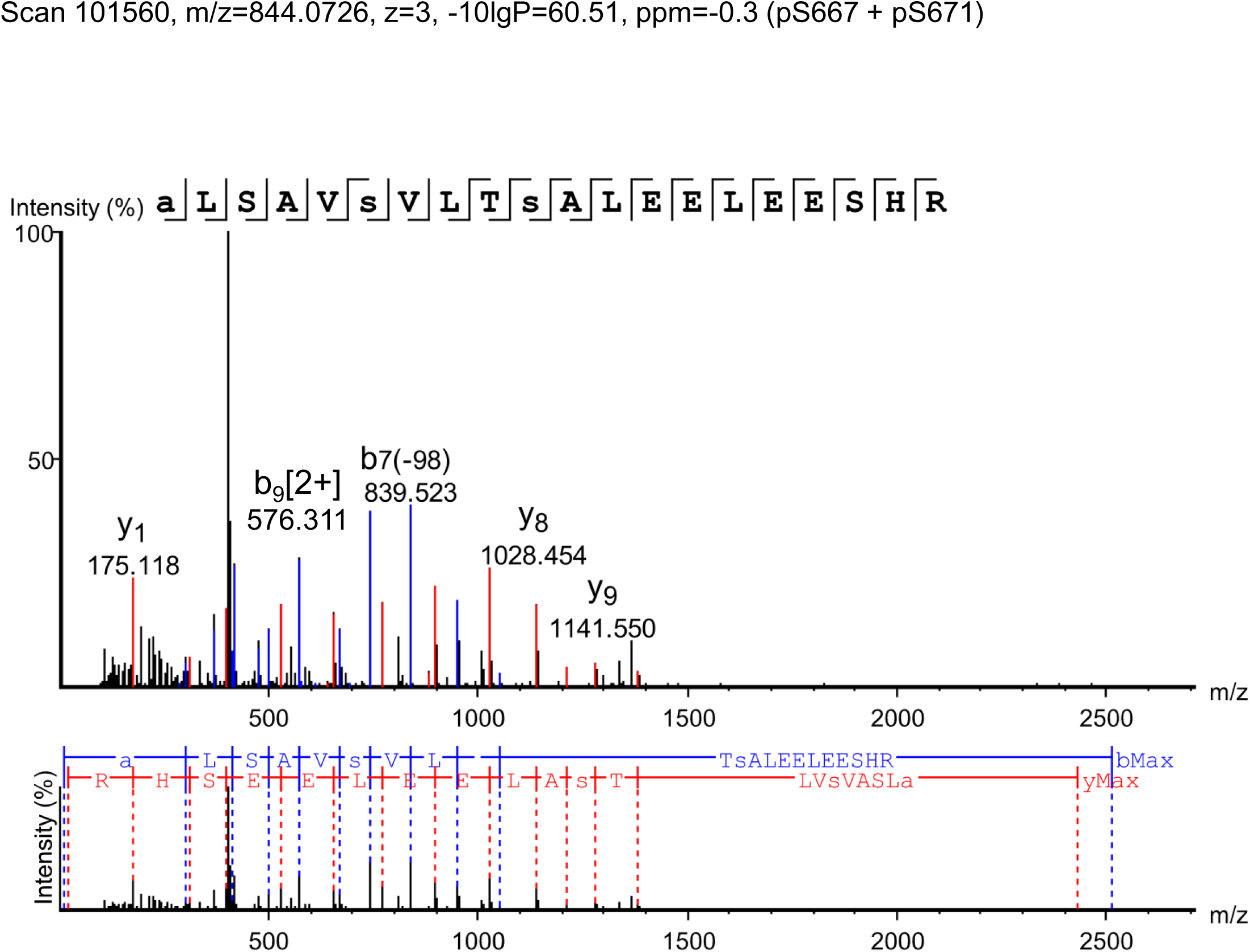
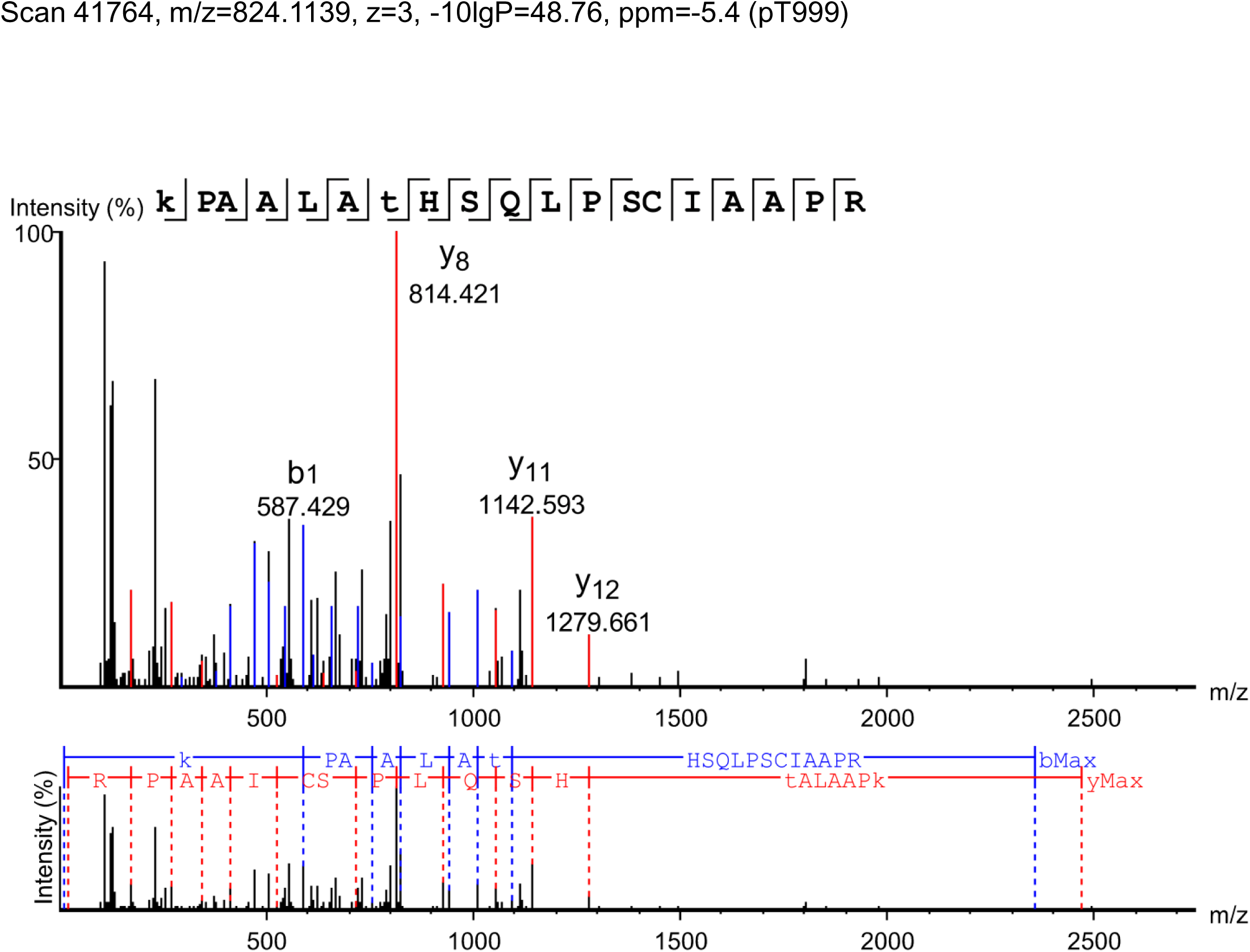
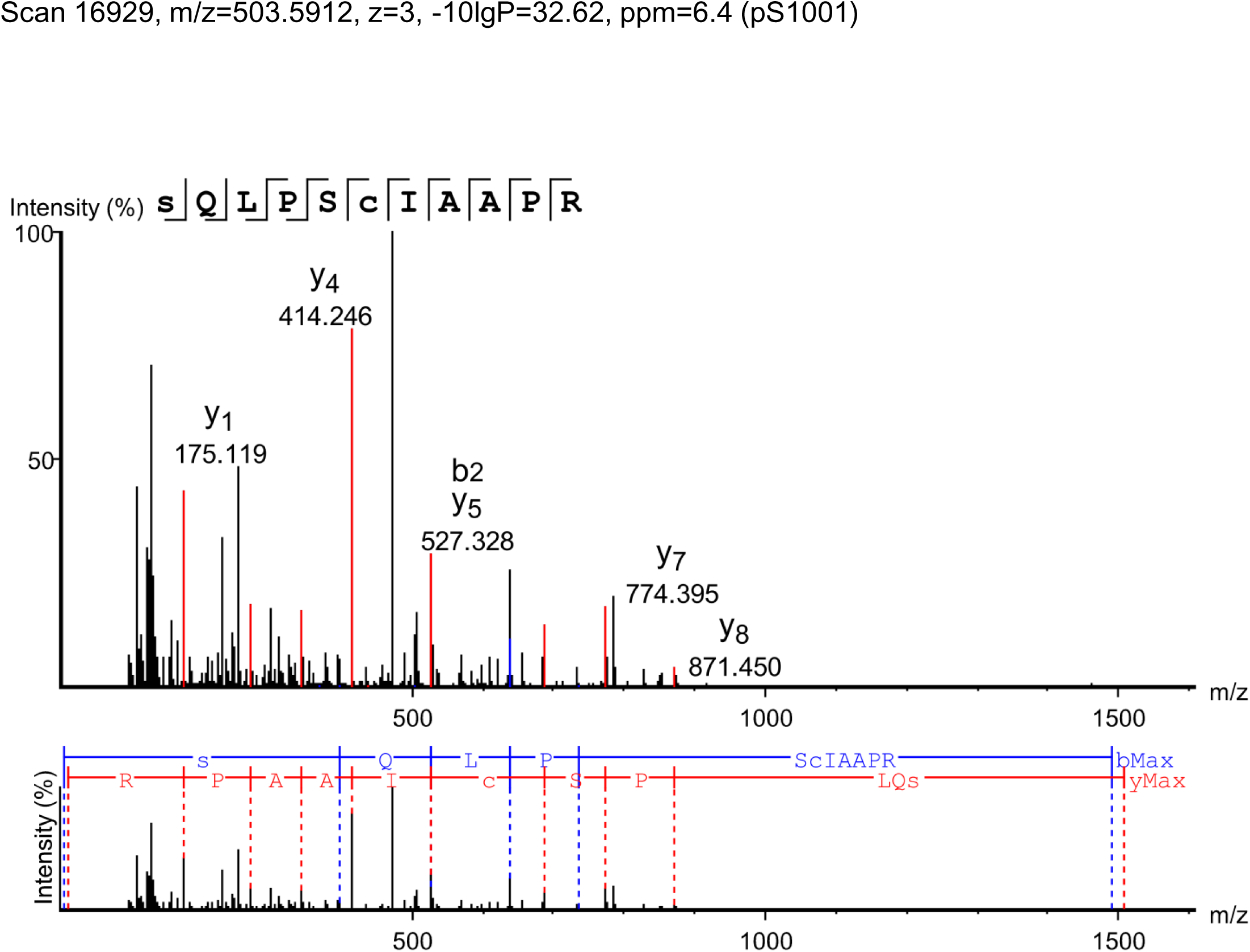
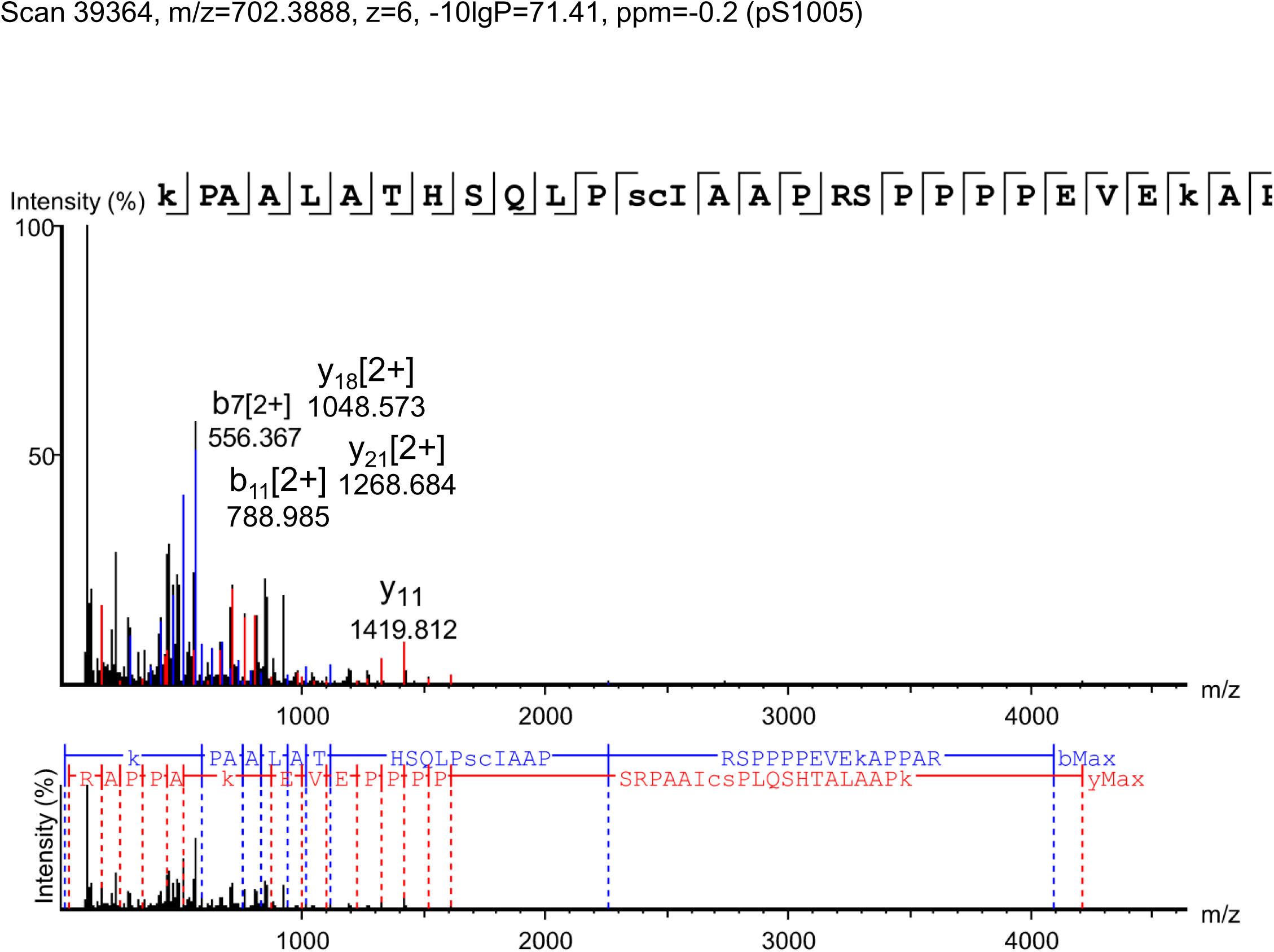
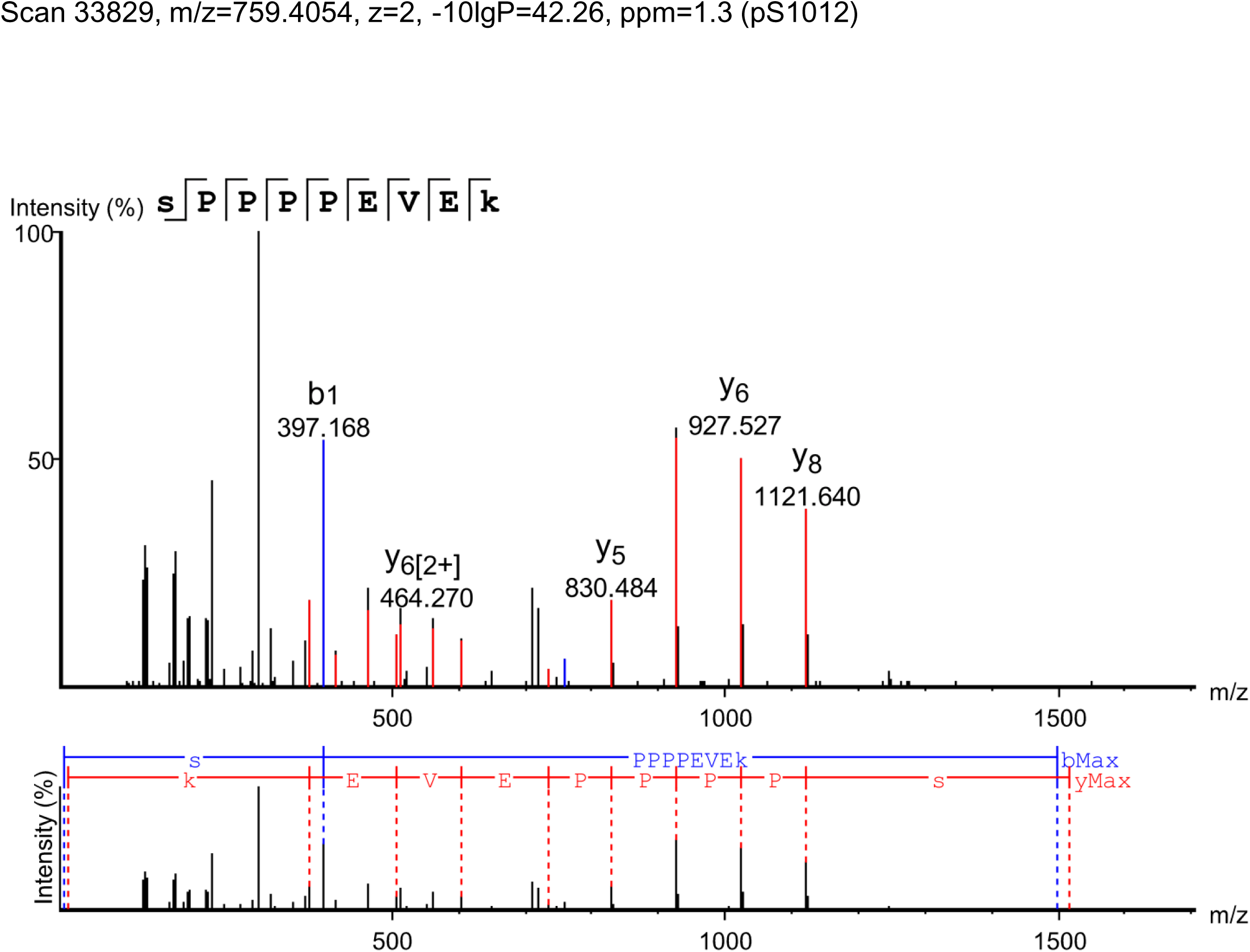
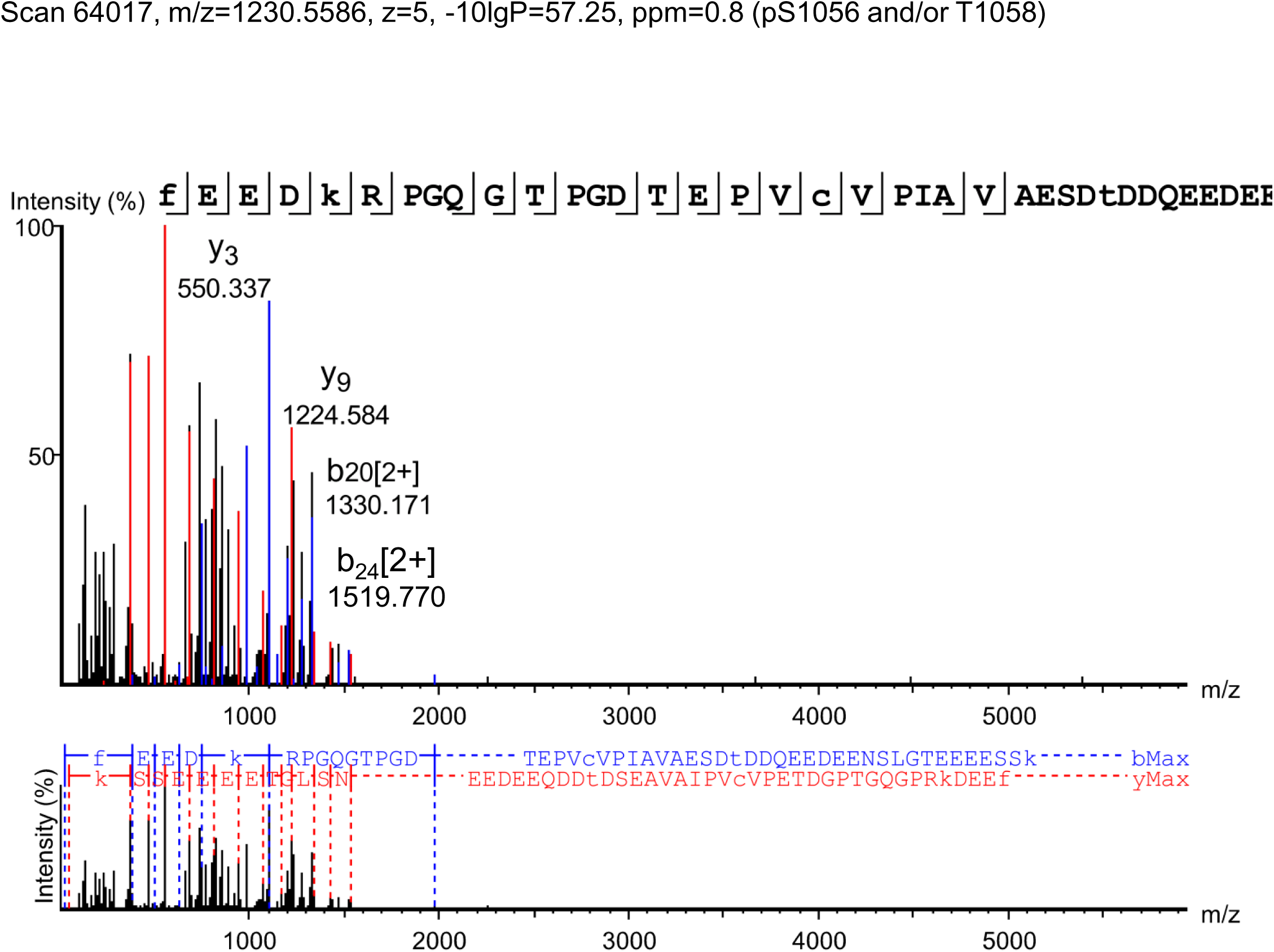
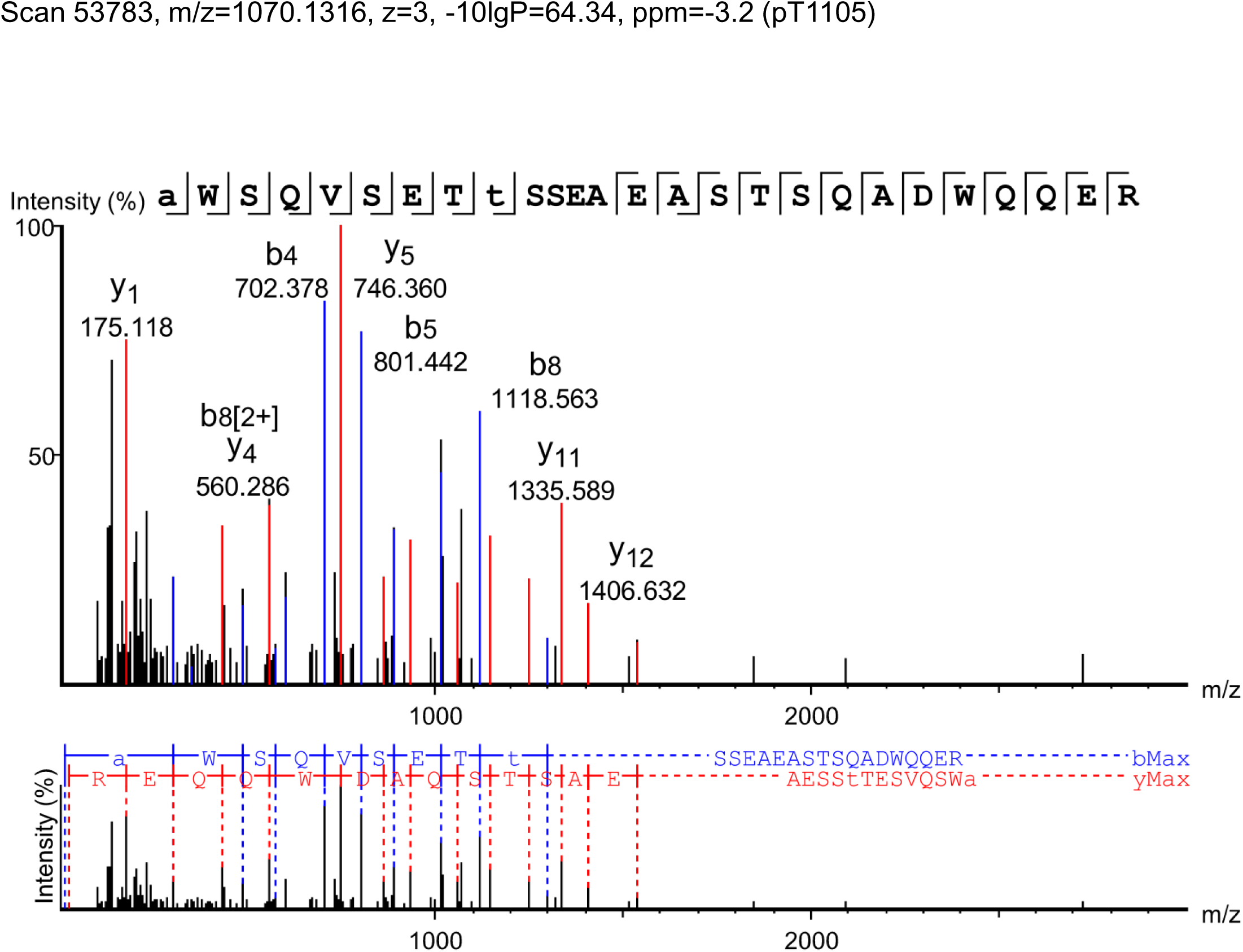
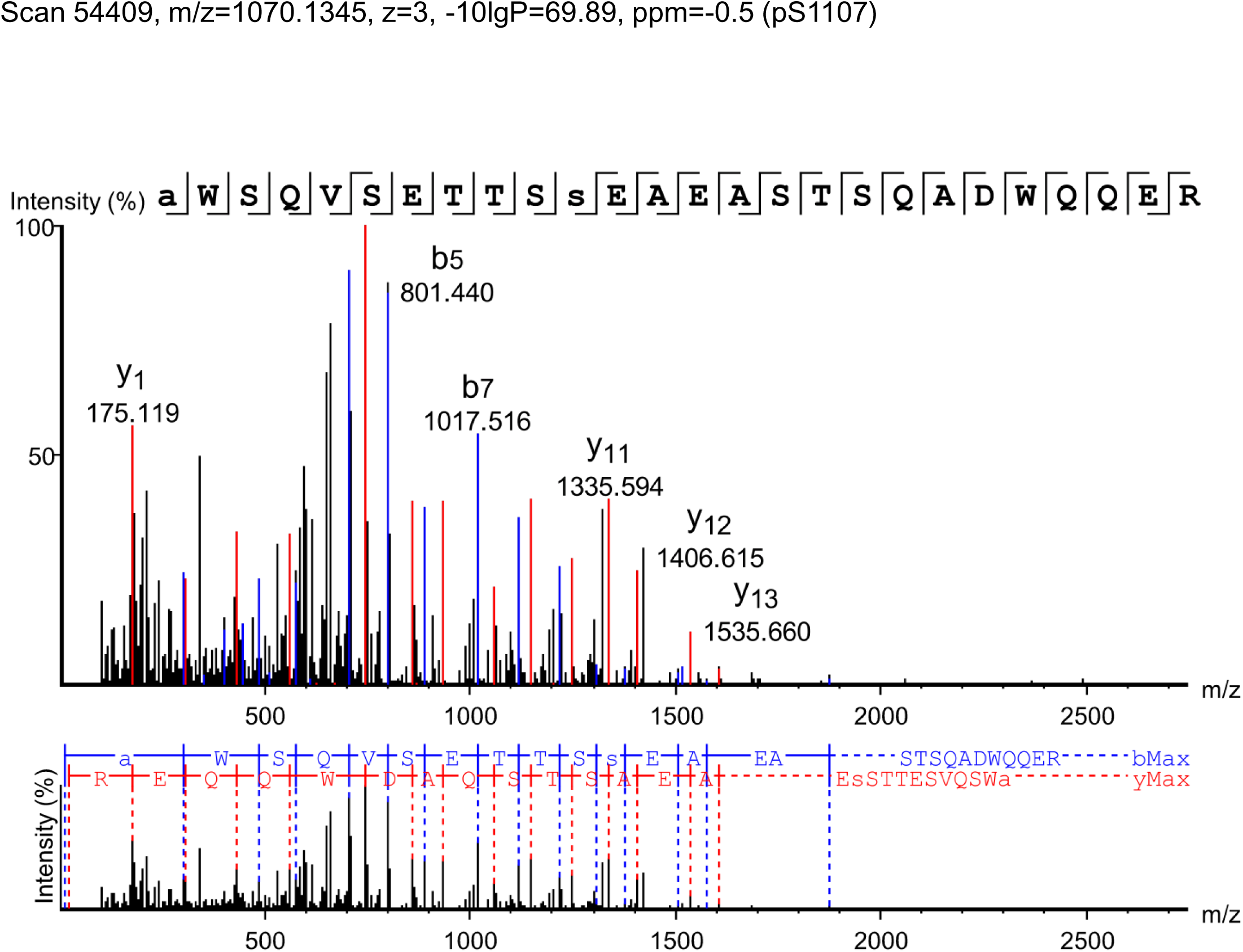
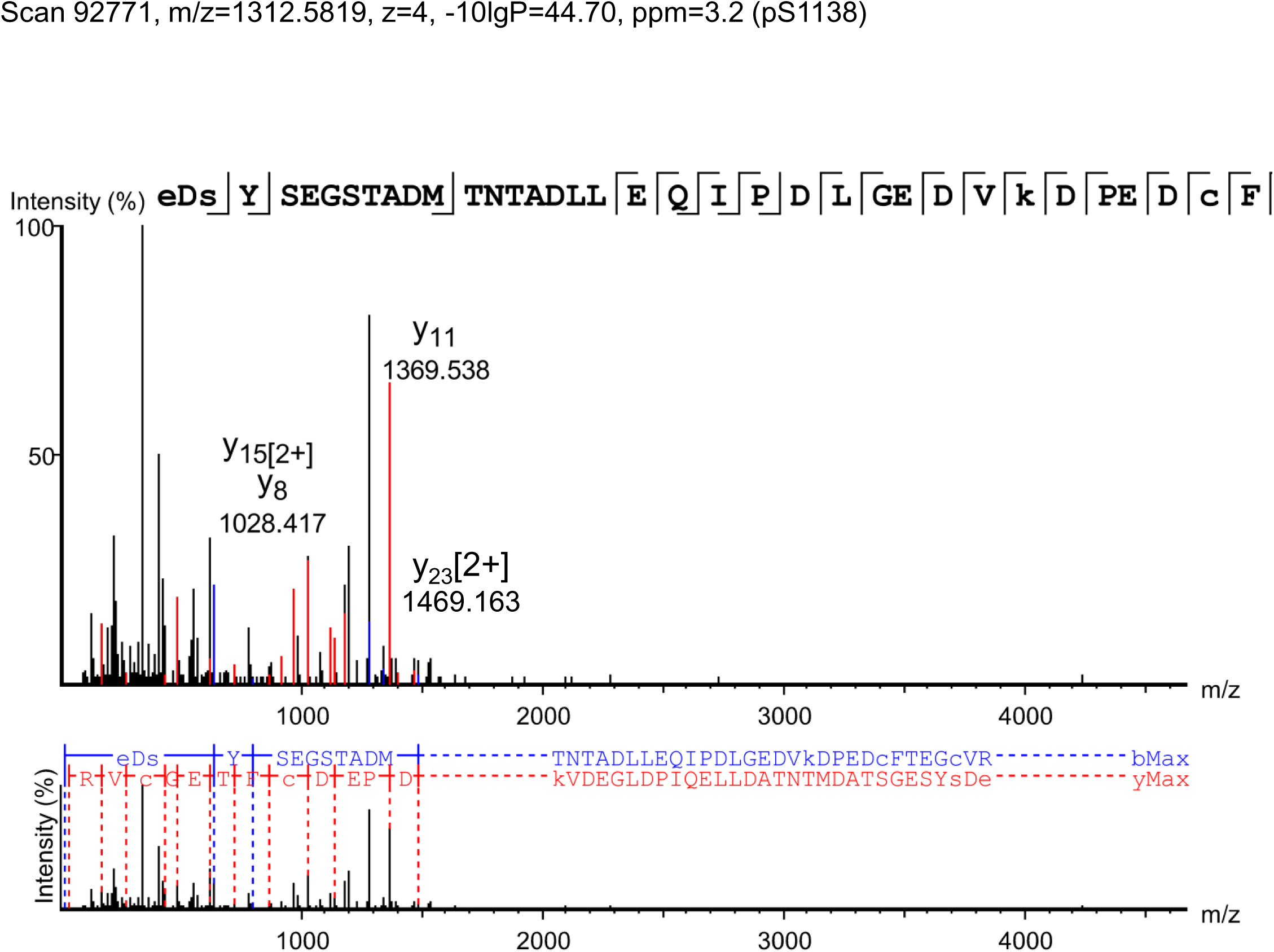
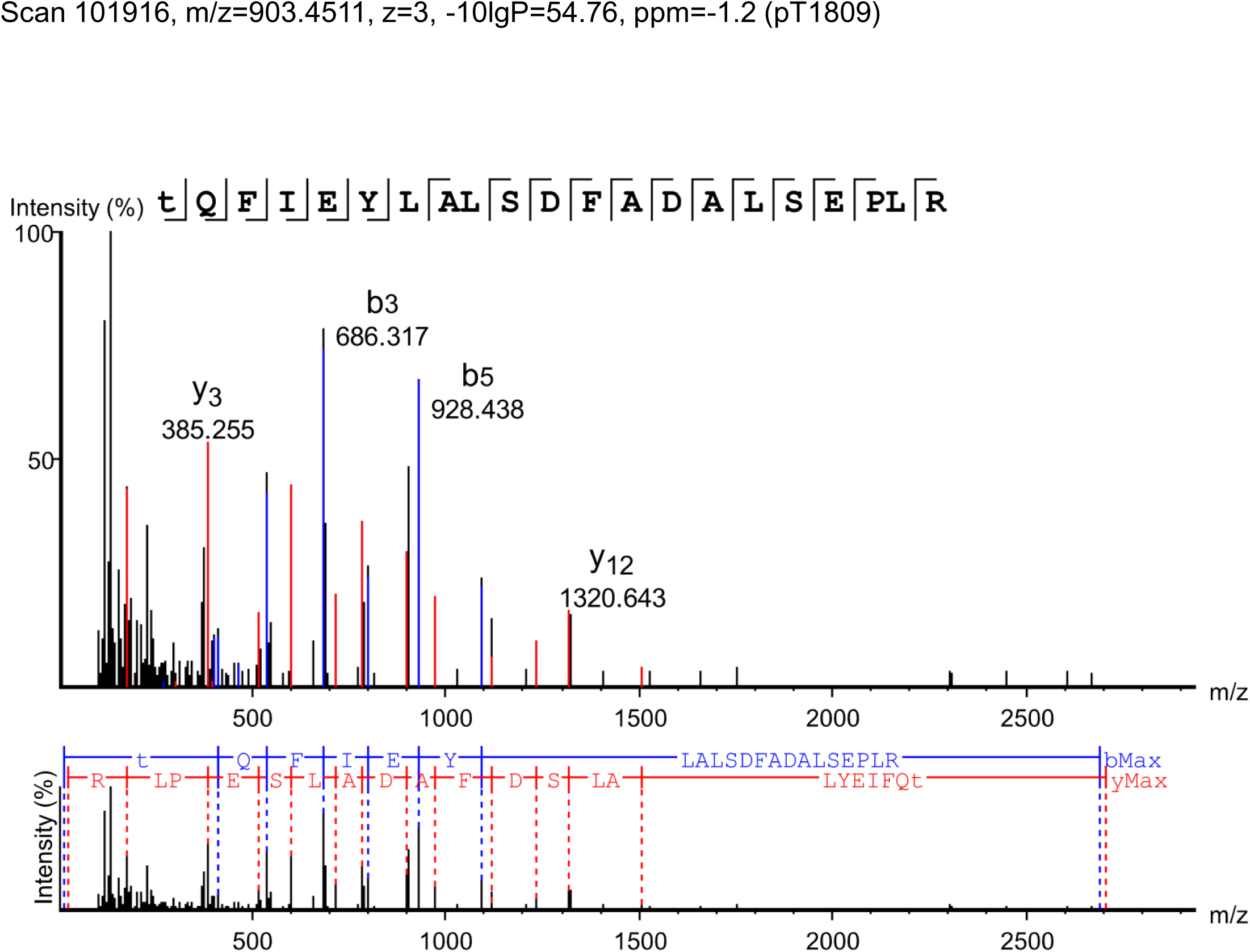
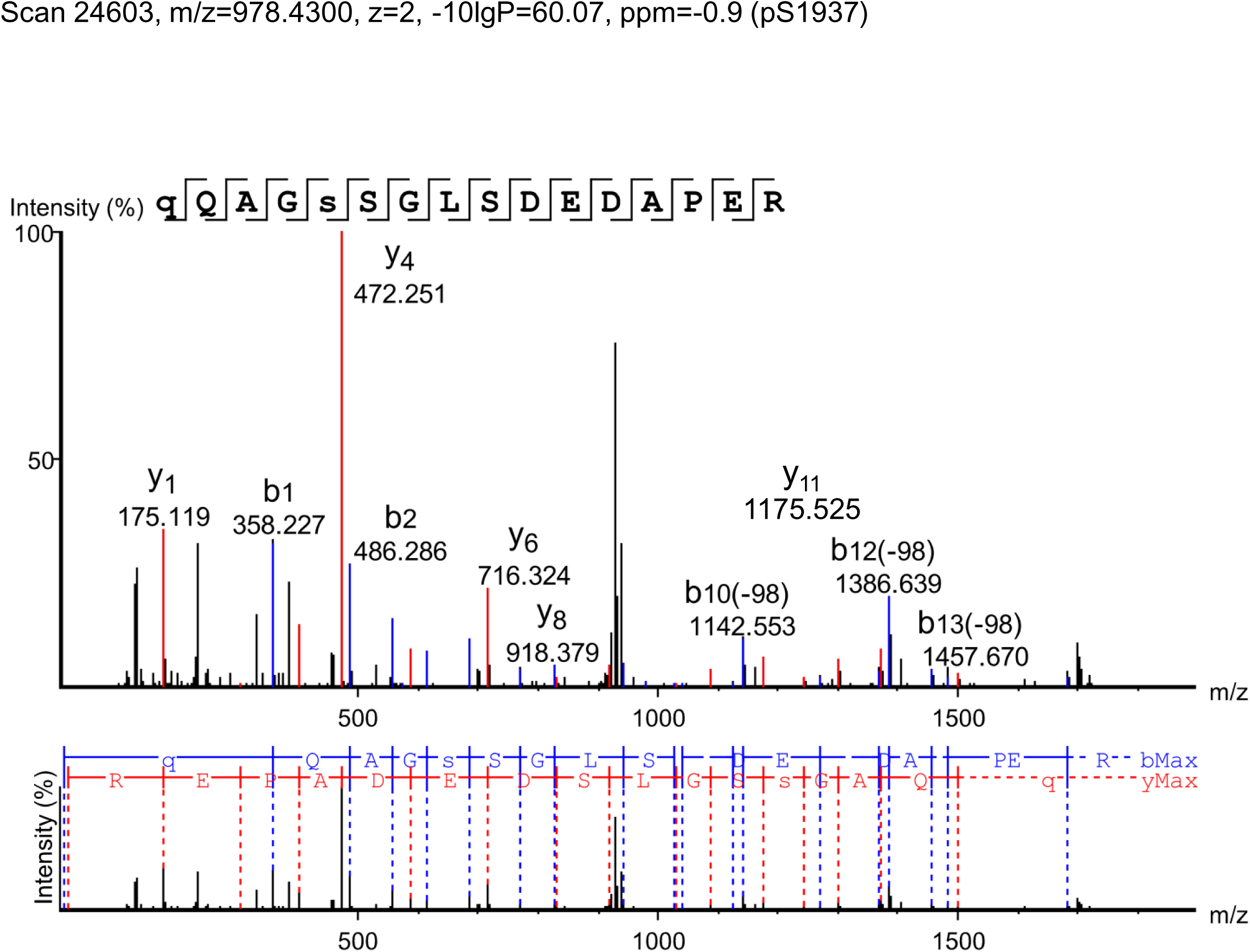
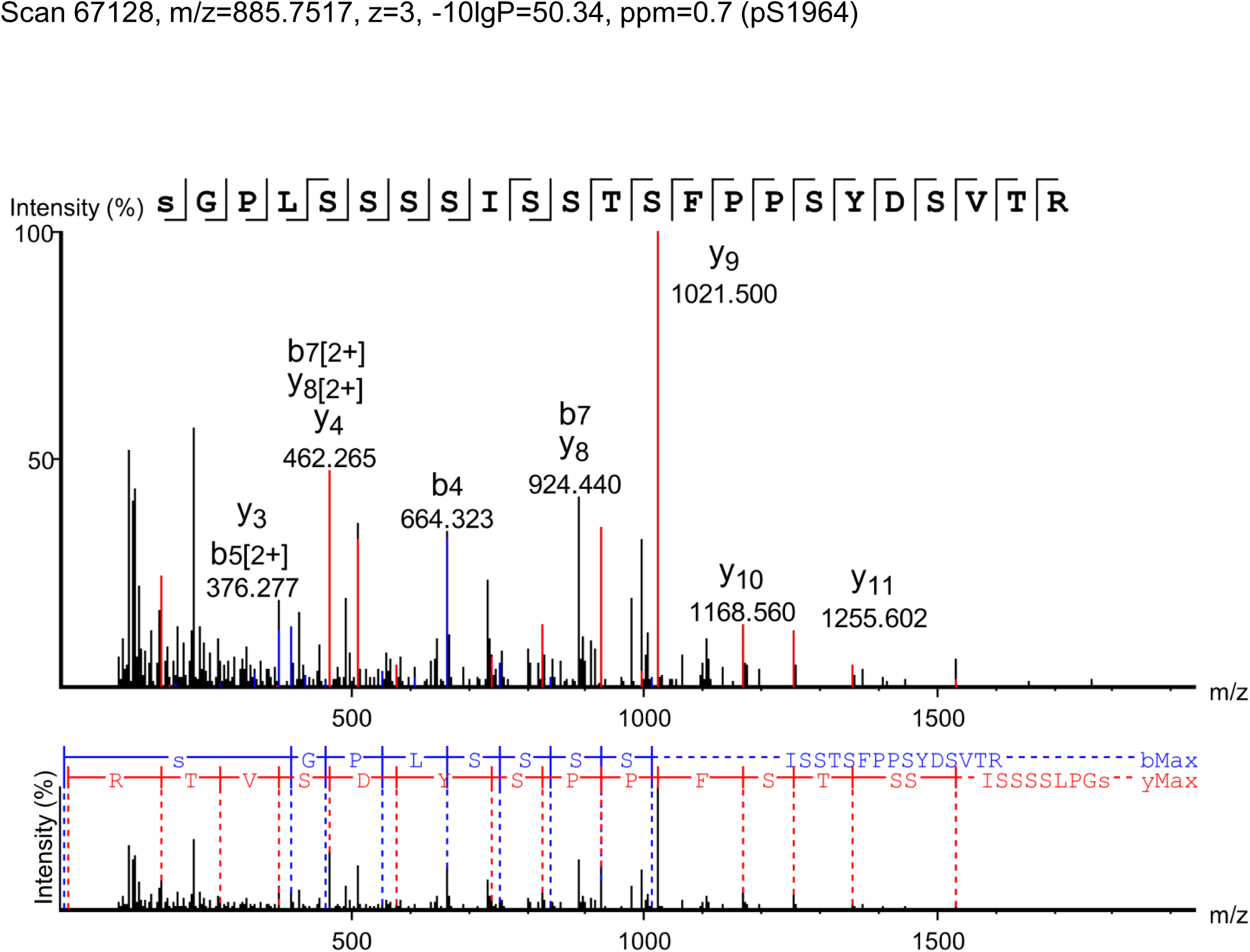
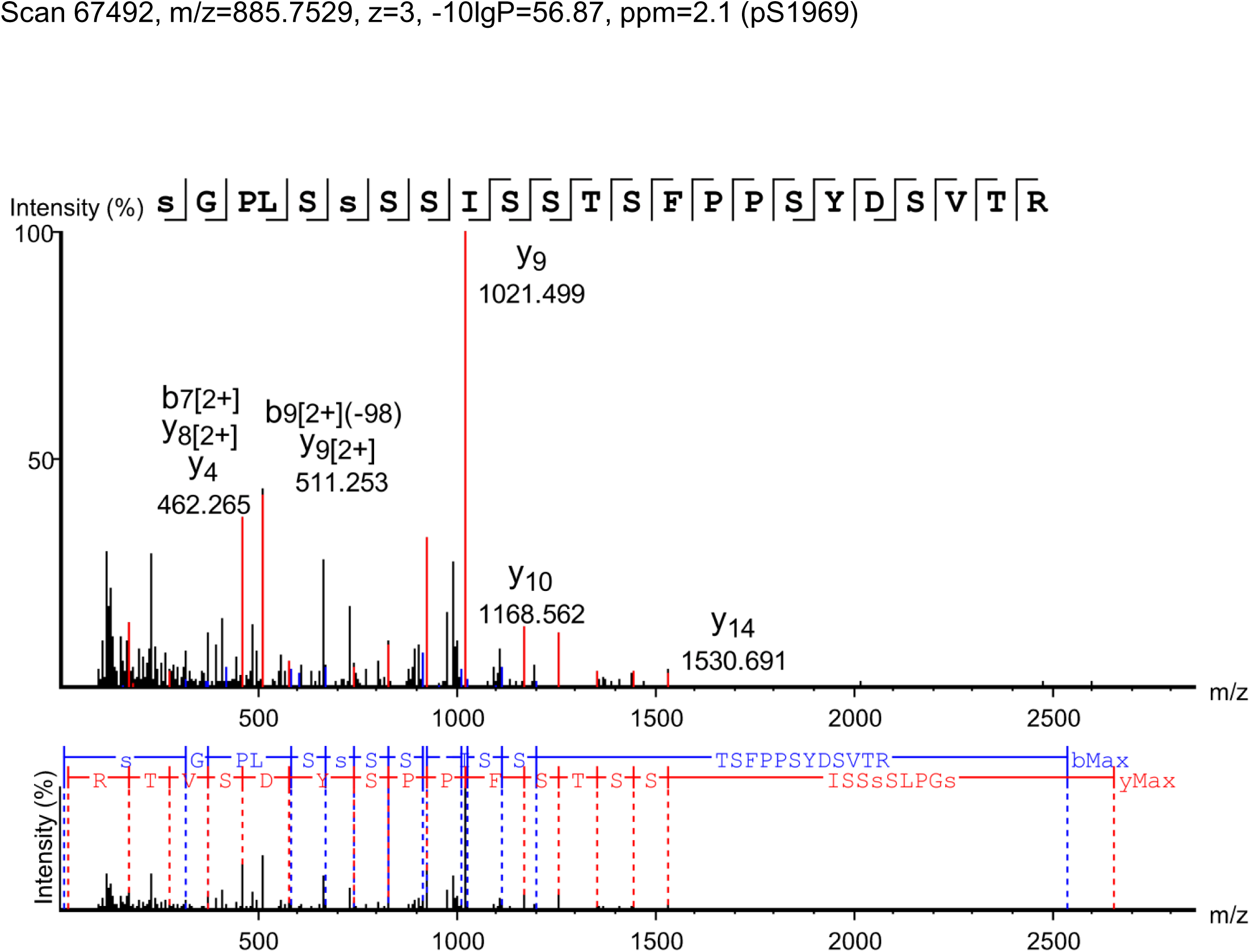
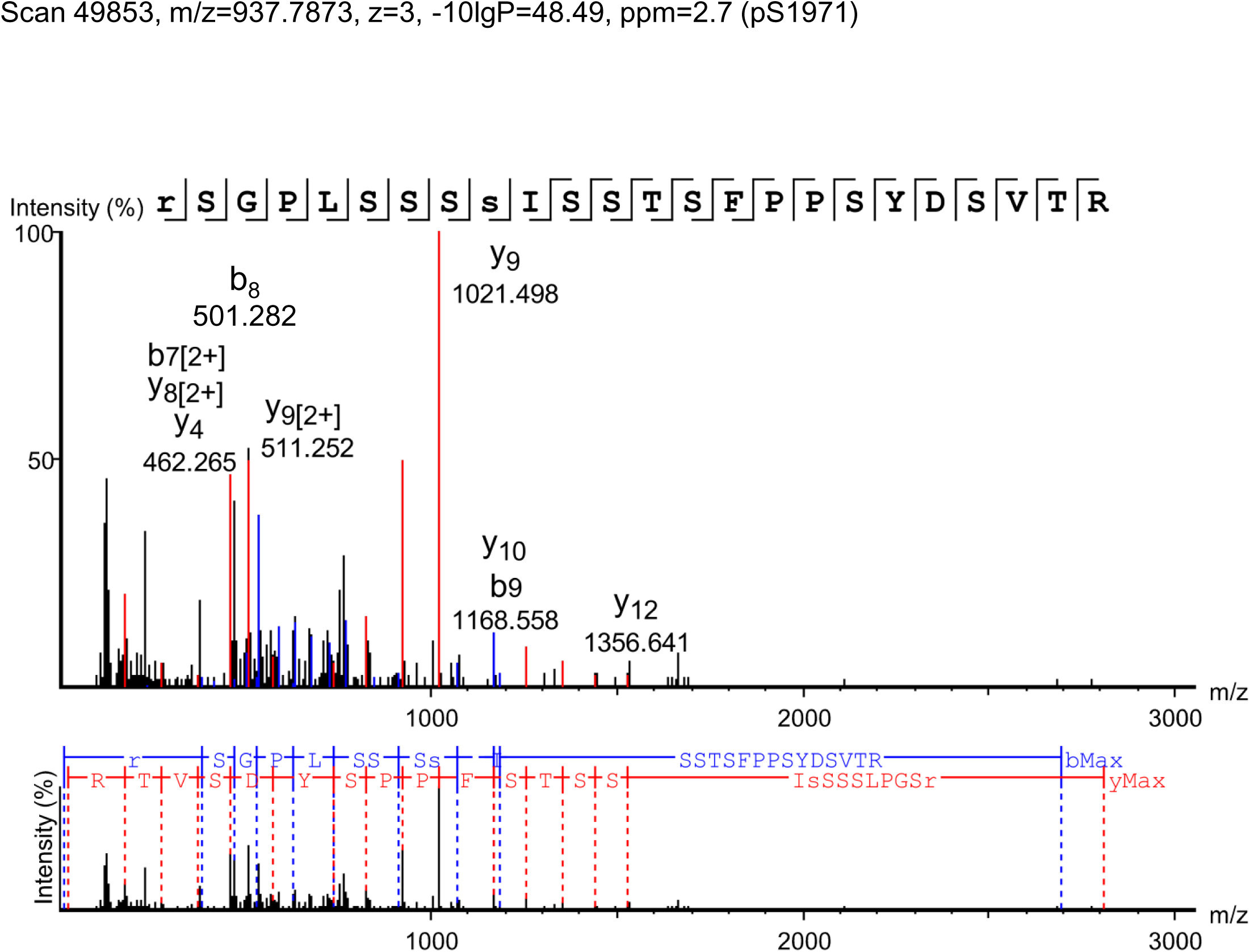
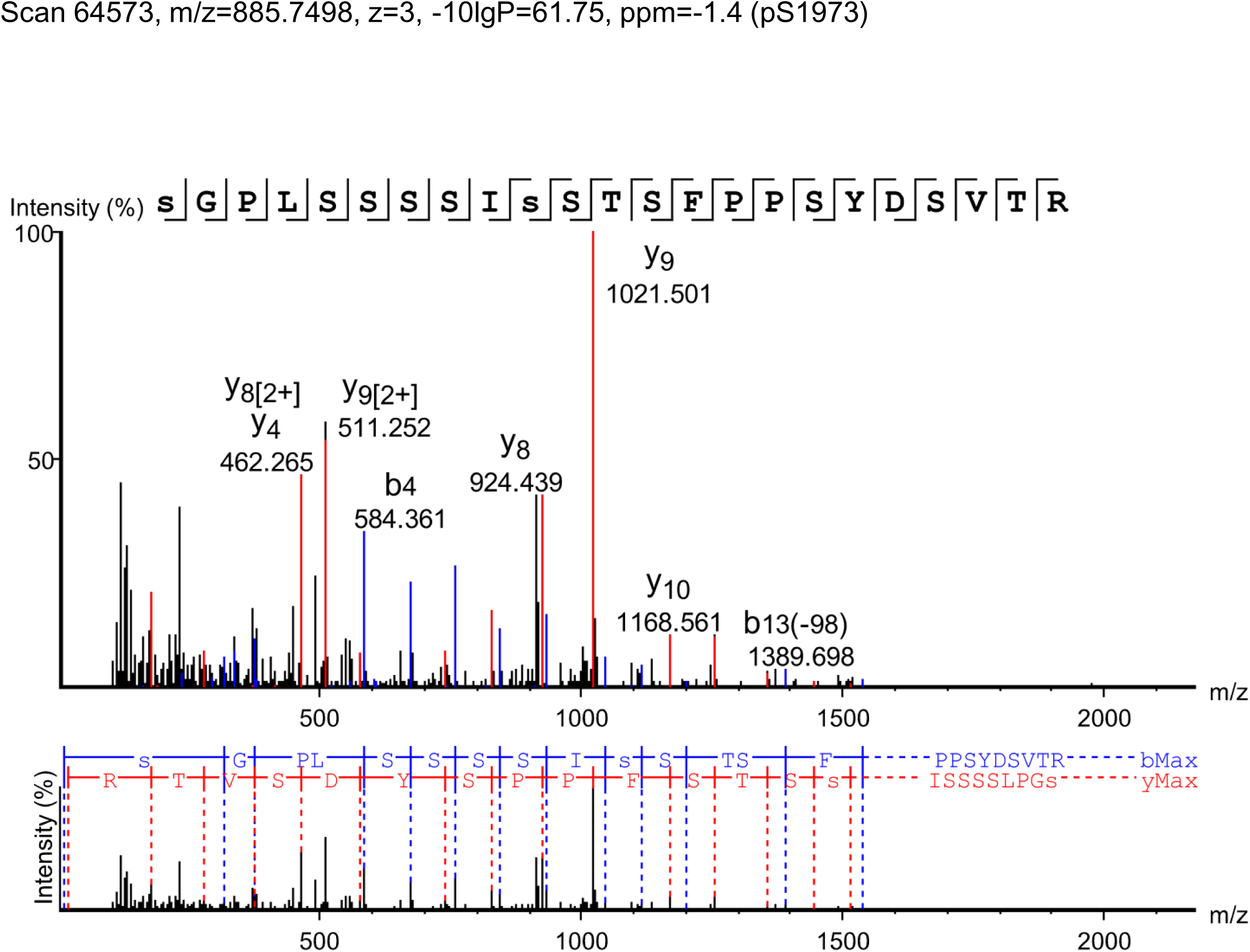
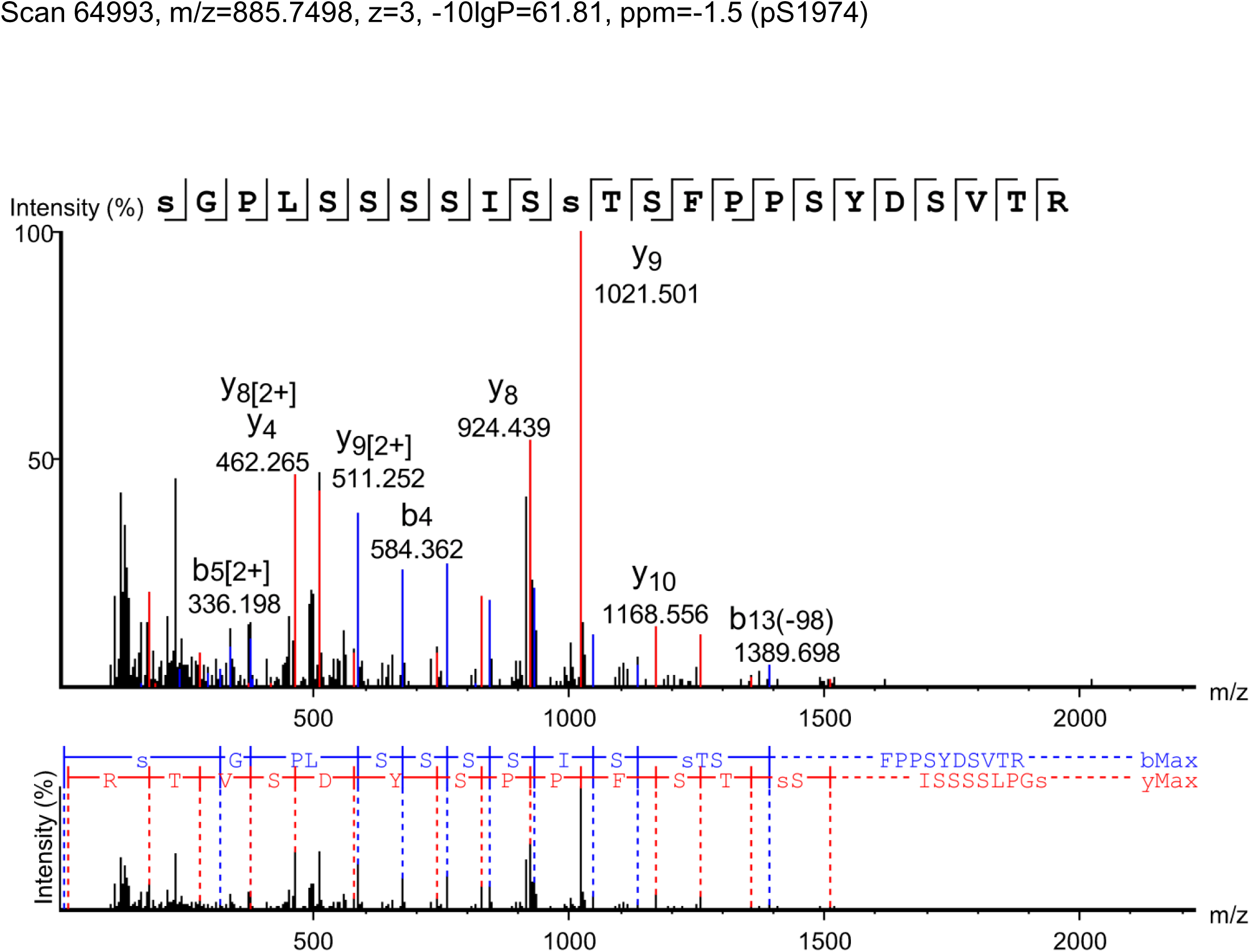
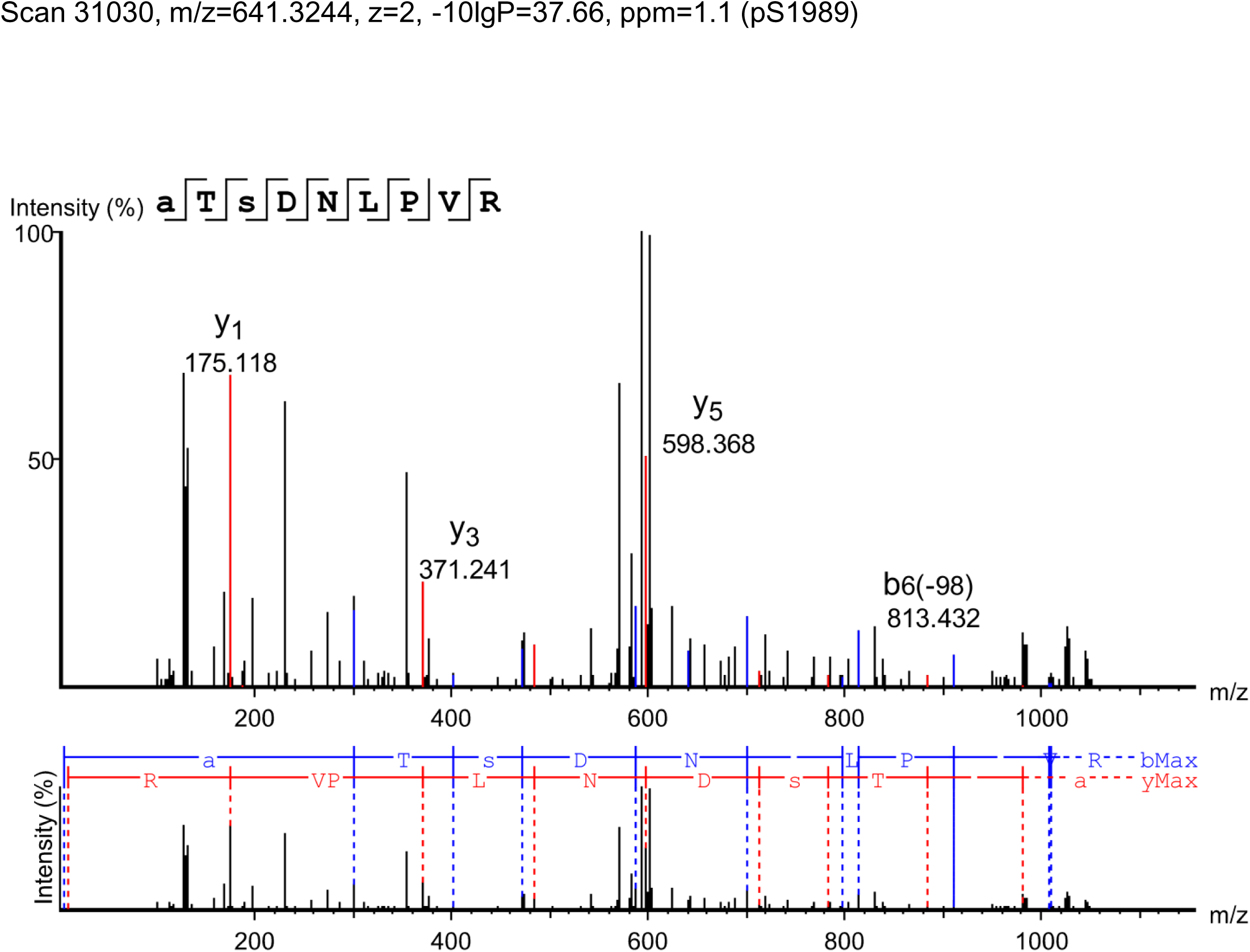
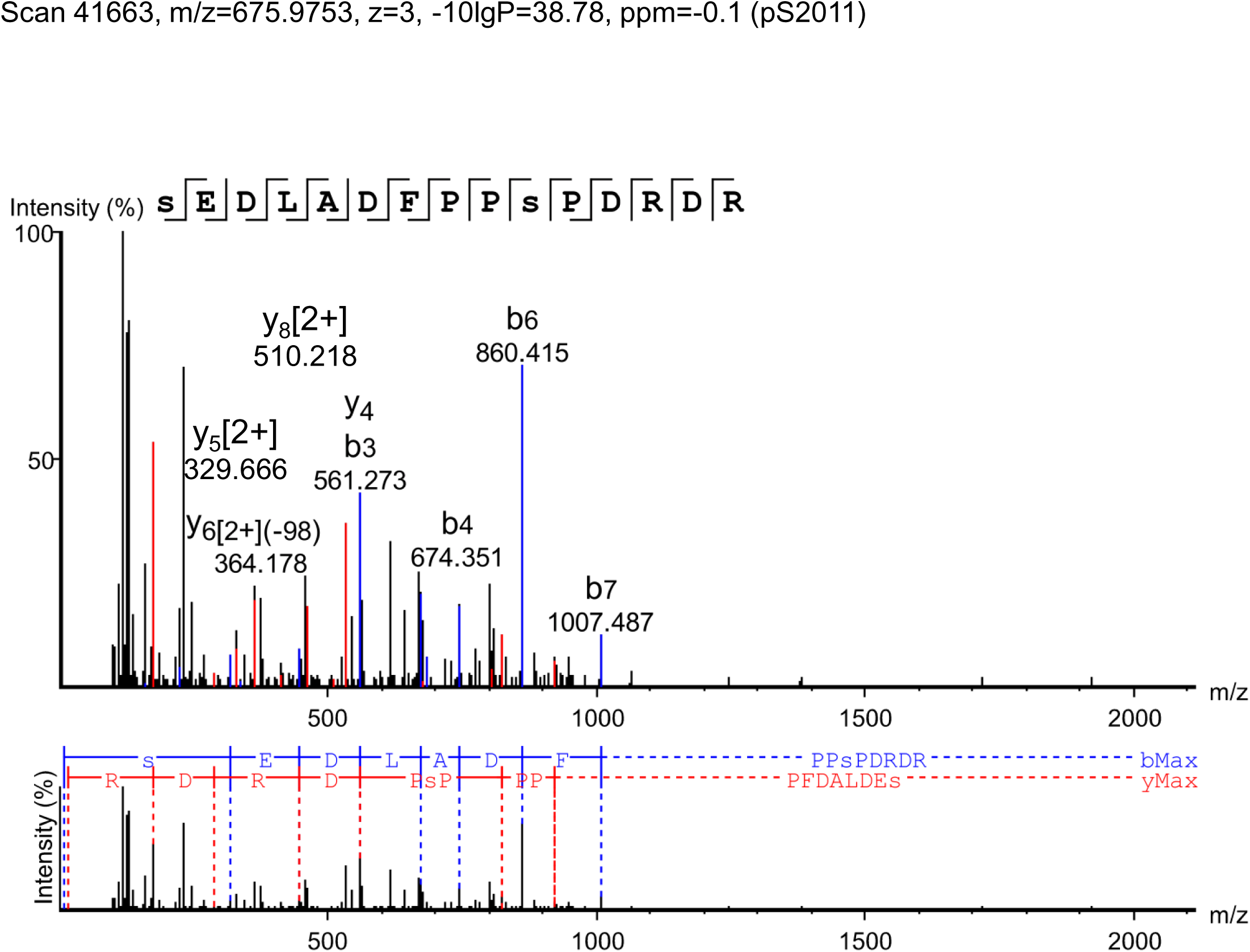
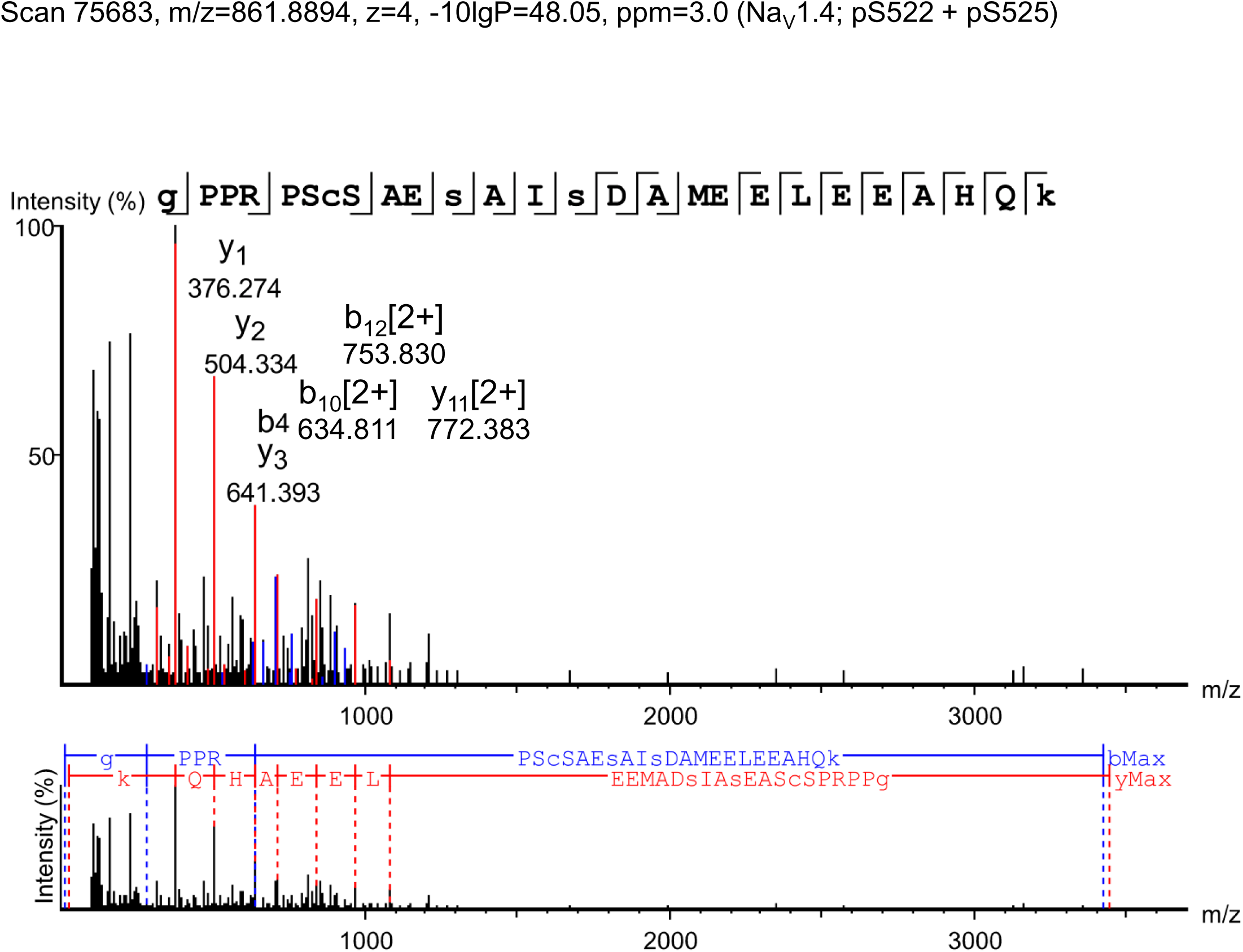
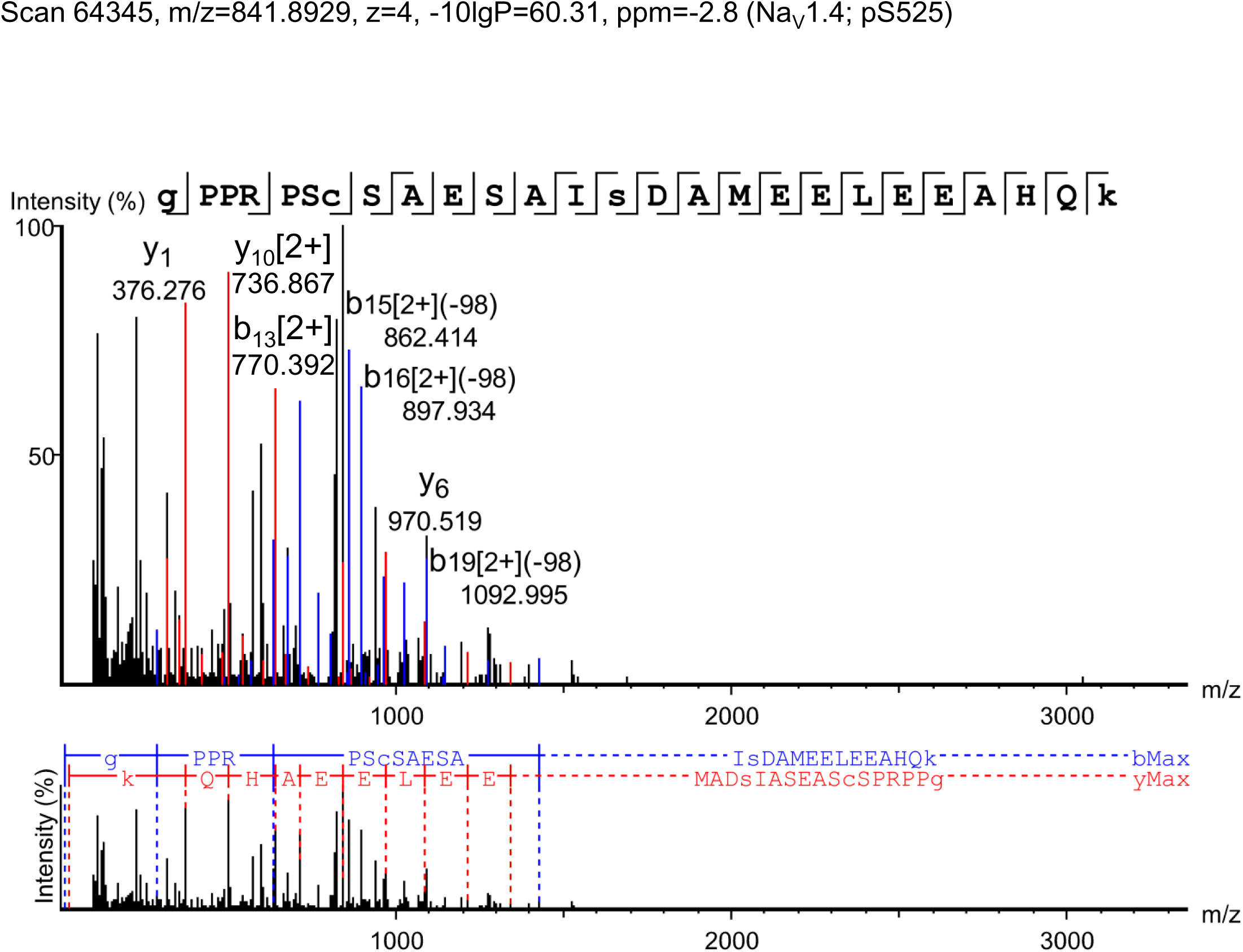
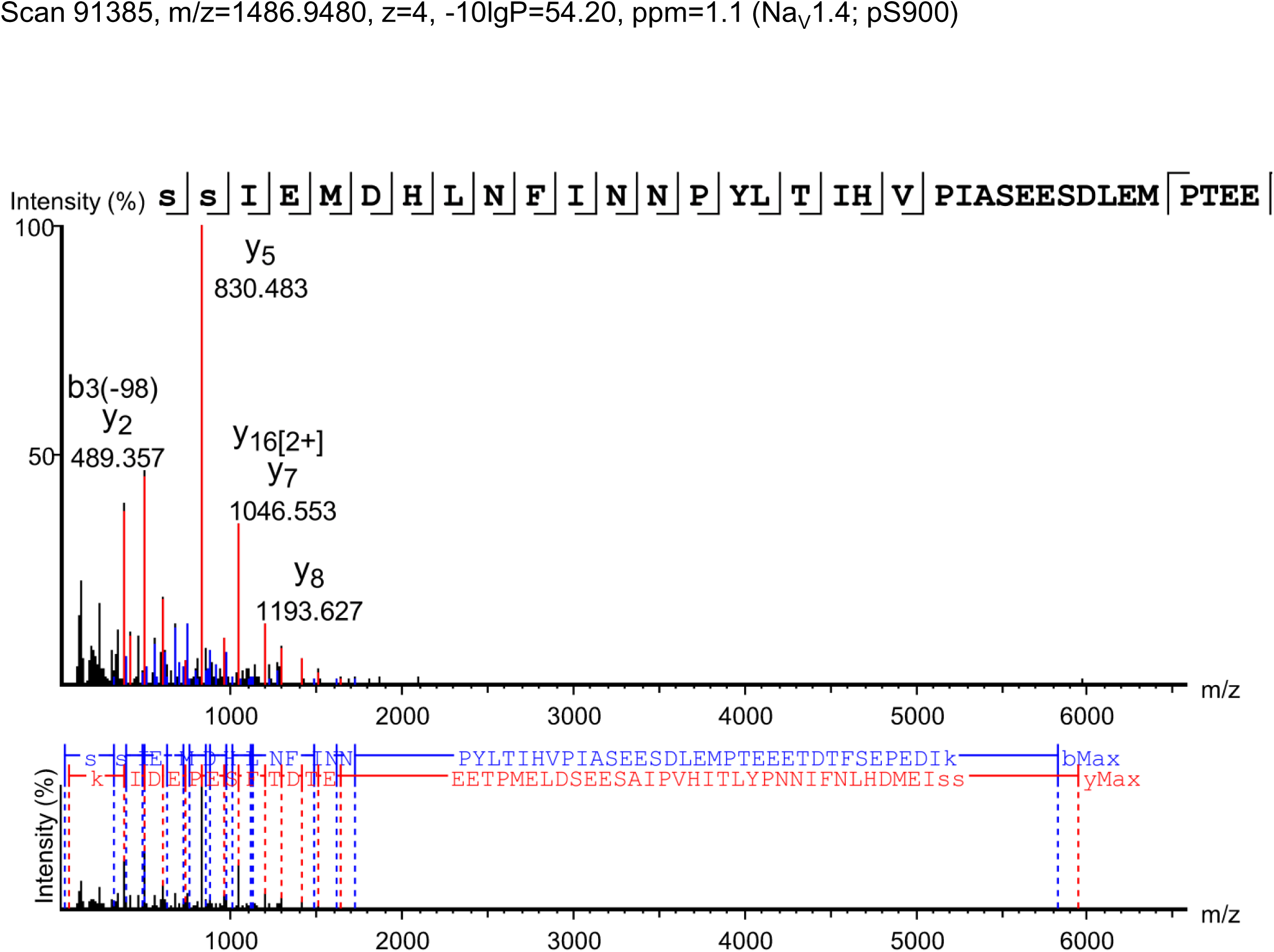
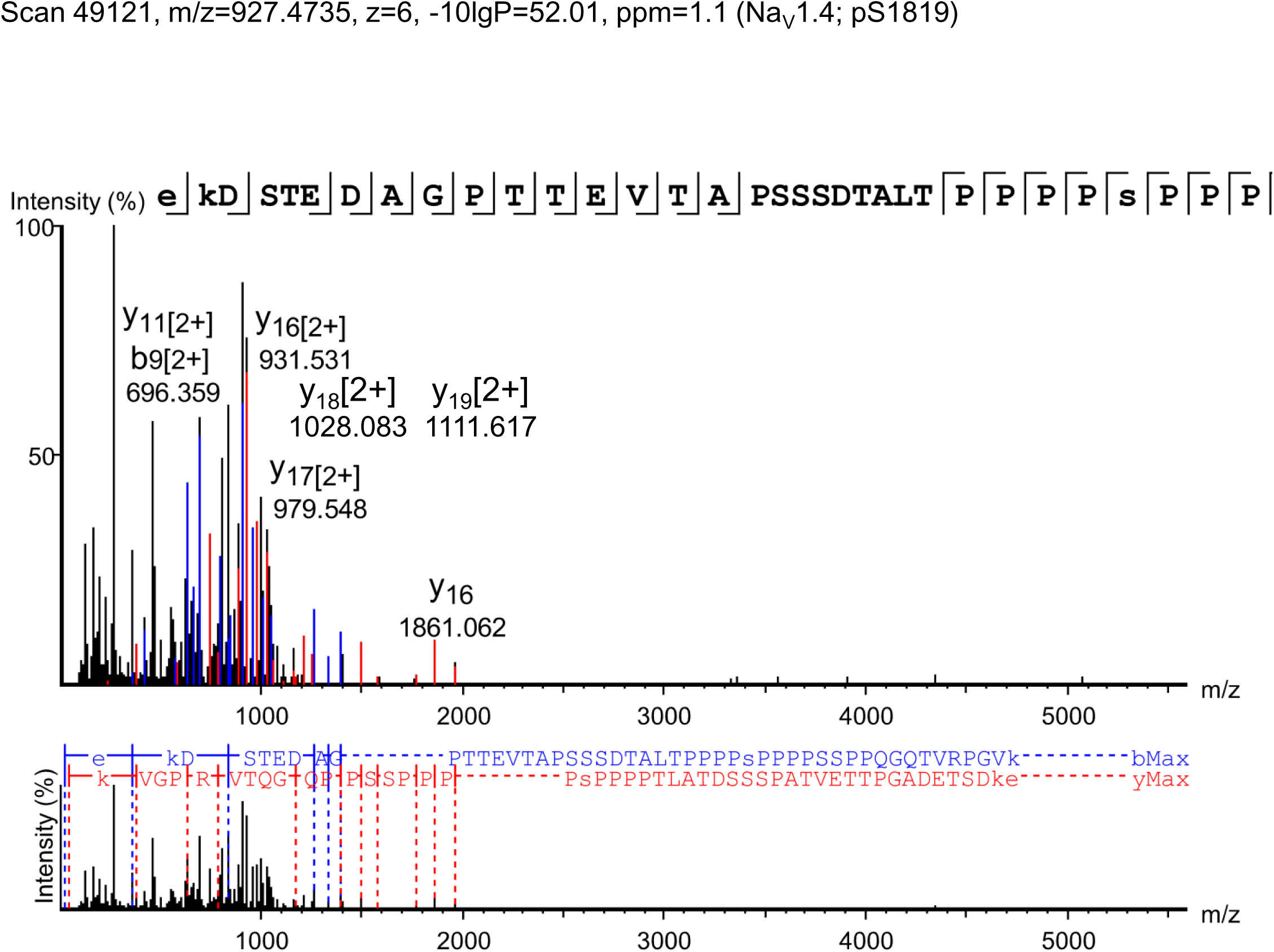
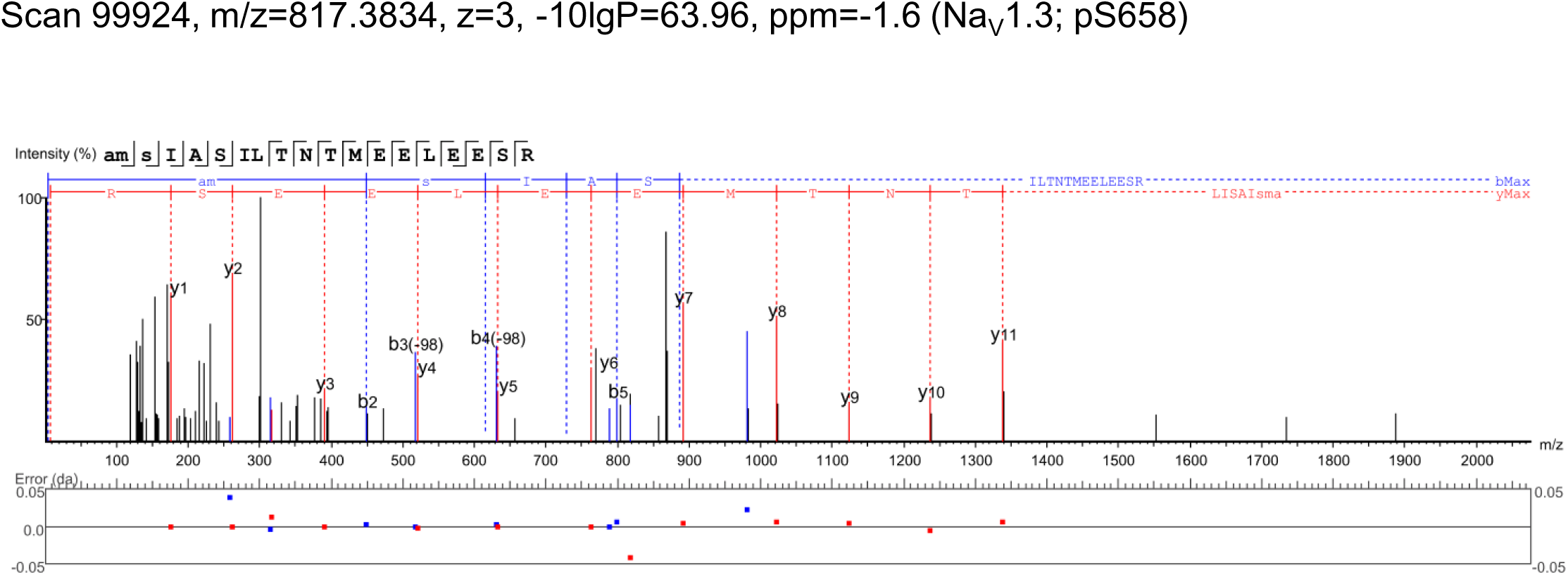
Phosphorylation sites, phosphopeptides and site-discriminating ions identified in co-immunoprecipitated NaV α subunits from Sham and TAC mouse left ventricles using MS

